# HiTaC: a hierarchical taxonomic classifier for fungal ITS sequences compatible with QIIME2

**DOI:** 10.1101/2020.04.24.014852

**Authors:** Fábio M. Miranda, Vasco C. Azevedo, Rommel J. Ramos, Bernhard Y. Renard, Vitor C. Piro

## Abstract

**Background:** Fungi play a key role in several important ecological functions, ranging from organic matter decomposition to symbiotic associations with plants. Moreover, fungi naturally inhabit the human body and can be beneficial when administered as probiotics. In mycology, the internal transcribed spacer (ITS) region was adopted as the universal marker for classifying fungi. Hence, an accurate and robust method for ITS classification is not only desired for the purpose of better diversity estimation, but it can also help us gain a deeper insight into the dynamics of environmental communities and ultimately comprehend whether the abundance of certain species correlate with health and disease. Although many methods have been proposed for taxonomic classification, to the best of our knowledge, none of them fully explore the taxonomic tree hierarchy when building their models. This in turn, leads to lower generalization power and higher risk of committing classification errors.

**Results:** Here we introduce HiTaC, a robust hierarchical machine learning model for accurate ITS classification, which requires a small amount of data for training and can handle imbalanced datasets. HiTaC was thoroughly evaluated with the established TAXXI benchmark and could correctly classify fungal ITS sequences of varying lengths and a range of identity differences between the training and test data. HiTaC outperforms state-of-the-art methods when trained over noisy data, consistently achieving higher F1-score and sensitivity across different taxonomic ranks, improving sensitivity by 6.9 percentage points over top methods in the most noisy dataset available on TAXXI.

**Conclusions:** HiTaC is publicly available at the Python package index, BIO-CONDA and Docker Hub. It is released under the new BSD license, allowing free use in academia and industry. Source code and documentation, which includes installation and usage instructions, are available at https://gitlab.com/dacs-hpi/hitac.

## 1 Background

In the last decades, there has been a surge of interest in characterizing the mycoflora of communities. This is due to the fact that fungi are key organisms in several important ecological functions, ranging from nutrient recycling to mycorrhizal associations and decomposition of wood and litter [1]. Moreover, fungi naturally inhabit the human body and may benefit the host when administered as probiotics [2], yet little is known about the fungal microbiota of healthy individuals when compared to the bacterial microbiome [3]. Furthermore, identifying the presence of fungi can be challenging, as they rarely form structures visible to the naked eye, and similar structures are frequently composed of several distinct species [4]. Hence, it became common practice to use DNA sequencing in addition to morphological studies in contemporary mycology [5].

One of the approaches that can be employed in environmental studies is whole-genome shotgun sequencing (WGS), where the purpose is to obtain the genetic material of all organisms present in a given microbiome simultaneously [6]. Nonetheless, in studies where the end goal is only to estimate the diversity or discover the evolutionary distance of organisms in a microbiome, it is cheaper and more convenient to sequence only genetic markers such as 16S rRNA, which is well conserved in all bacteria [7] and also widely used to classify and identify archaea [8]. In mycology, the internal transcribed spacer (ITS) region was adopted as the universal marker for classifying fungi [9].

The main characteristic of 16S rRNA that makes it a good genetic marker is the presence of nine hypervariable regions V1–V9, which are flanked by conserved regions that can be used to amplify target sequences using universal primers [10]. However, 16S rRNA has fewer hypervariable domains in fungi, and consequently, it is more appropriate to use the ITS regions which are composed by 18S–26S rRNAs [9]. Additionally, the ease with which ITS is amplified makes it an appealing target for sequencing samples from environments as challenging as soil and wood, where the initial quantity and quality of DNA is low [4].

From a computational perspective, the taxonomic classification of ITS fungal sequences can be performed by using either similarity-based or machine learning techniques. Similarity-based approaches, such as BLAST [11], require the alignment of a query sequence against all sequences available in a reference database, e.g., Silva [12] and UNITE [13]. However, a major limitation of similarity-based methods is the dependence on homologs in databases, which are not always available, while machine learning methods can learn only the relevant features and criteria for classification and can, therefore, be applied to any new sequence [14].

One of the first machine learning software proposed to perform taxonomic classification was the RDP-Classifier, which uses 8-mer frequencies to train a Naive Bayes classification algorithm [15]. Improving upon the RDP-Classifier, a novel Naive Bayes classifier with multinomial models was developed to increase the accuracy [16]. This multinomial classifier was later reimplemented and optimized on Microclass [17].

Other machine learning approaches have also been proposed in the literature, such as the k-Nearest Neighbor (KNN) algorithm [18]. Given a query sequence, the KNN algorithm identifies the k-most similar sequences in a database and uses the taxonomic information from each of those sequences to determine the consensus taxonomy. Q2 SK [19] implements several supervised learning classifiers from scikit-learn [20], e.g., Random Forest, Support Vector Machines (SVM) and Gradient Boosting. Q2 SK is flexible and allows the selection of features and classifiers.

Despite the advances made in taxonomic classification, there are still opportunities for enhancement due to the complexity of the data. Furthermore, to the best of our knowledge, the machine learning software available may lack in predictive performance since they perform flat classification, which means that they are only trained on leaf node labels and do not fully explore the hierarchical structure of the taxonomic tree when building their models (see Fig. S1 for a depiction of the hierarchical structure of the taxonomic tree). The main disadvantage of this approach is that the information about parent-child class relationships in the hierarchical structure is not entirely considered, which may cause flat classifiers to commit more errors than hierarchical approaches [21]. Therefore, we developed a new hierarchical taxonomic classifier for fungal ITS sequences, denominated HiTaC, which explores the hierarchical structure of the taxonomic tree during the training stage and improves the F1-score and sensitivity of fungal ITS predictions. Furthermore, HiTaC can be easily installed and is compatible with QIIME2, facilitating its adoption in existing ITS sequence analysis pipelines.

## 2 Implementation

### 2.1 The local hierarchical taxonomic classifier

HiTaC performs feature extraction, building of the hierarchical model, taxonomic classification, and reporting of the predictions. Figure 1 illustrates the algorithmic steps of HiTaC, where it starts by decomposing the DNA sequences from both reference and query files into their constituents k-mers. Next, two k-mer frequency matrices are built for the reference and query sequences, respectively, where each row contains the k-mer frequency for a sequence. Logistic regression classifiers implemented in the library scikit-learn [20] are trained for each parent node in the taxonomic hierarchy, using the local classifier per parent node implementation from a library called HiClass that we developed and published previously [22]. The k-mer frequency matrix and taxonomic labels of the reference sequences are used as training data to predict children labels (Figure S2). Predictions are performed using the k-mer frequency matrix computed for the query sequences and are reported in a top-down approach, based on the taxonomic hierarchy. For example, if the classifier at the root node predicts that a query sequence belongs to the phylum Ascomycota, then only that branch of the taxonomic tree will be considered for the remaining taxonomic levels.

**Fig. 1:**
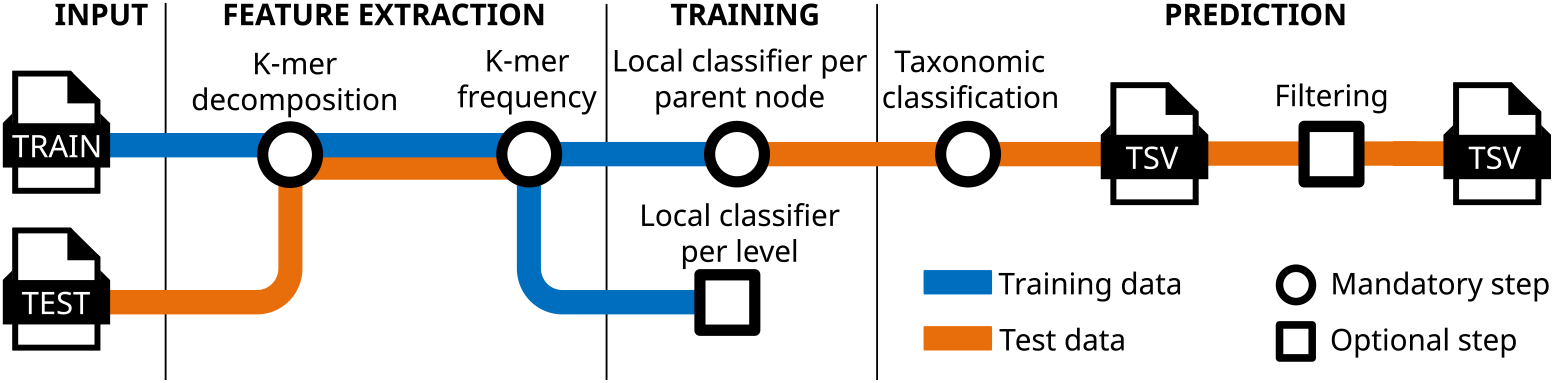
Sketch of the key algorithmic components of HiTaC.

### 2.2 Classification uncertainty

To reduce classification errors when faced with uncertainties, we also construct a filter by training a logistic regression classifier for each level of the hierarchy. This filter computes confidence scores for all taxonomic ranks assigned by the local classifier per parent node and excludes labels that are below a user-defined threshold in a bottom-up approach (Figure S3). The confidence scores are computed using the `predict proba` method implemented in scikit-learn’s logistic regression classifier, which in turn uses a one-vs-rest approach; that is, it calculates the probability of each class using the logistic function and assuming it to be positive, then it normalizes these values across all the classes [23].

### 2.3 Code quality assurance

The code base adheres to the PEP 8 code style [24], which is enforced by `flake8` and the uncompromising code formatter `black` to ensure high code quality. Versioning complies with `SemVer` to increase reproducibility and facilitate dependency management by end users. The code is accompanied by unit tests that cover 98% of the code base and are automatically executed by our *continuous integration workflow* upon commits.

### 2.4 Installation and usage

HiTaC is hosted on GitLab^1^, where there is also documentation with installation and usage instructions. HiTaC is compatible with Python 3.8+ and can be installed on GNU/Linux, Windows and macOS. It can be easily obtained with `pip install hitac`, `conda install-c` `bioconda hitac` or `docker pull mirand863/hitac`. Listing 1 shows a basic example of how to fit a hierarchical model and predict the taxonomy using HiTaC. More elaborate examples can be found in the documentation.

**Listing 1** Example of how to use HiTaC to train a hierarchical classifier and predict taxonomy.

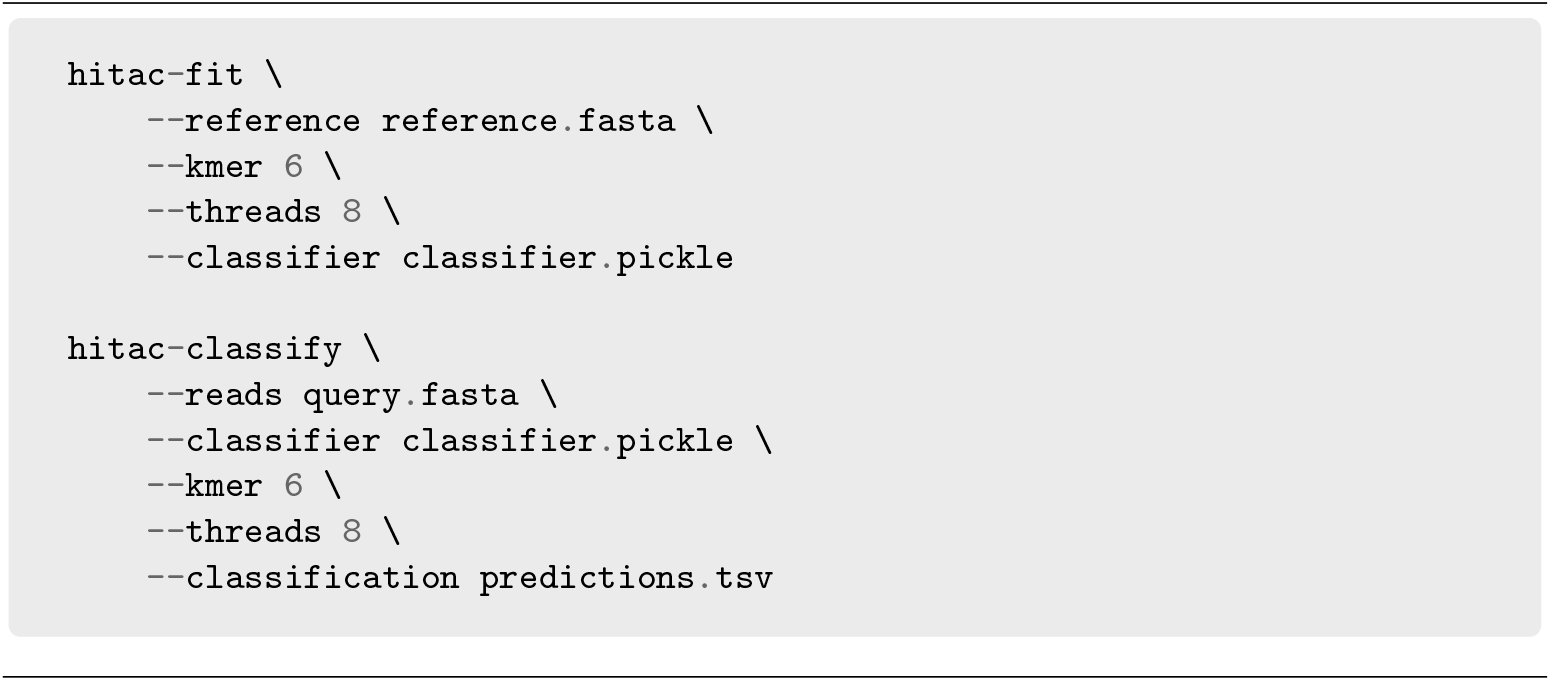

### 2.5 Evaluation

We evaluated HiTaC in five datasets and compared its performance against seventeen taxonomic classifiers using the TAXXI benchmark [25]. All methods evaluated were trained and tested with the same data in order to avoid bias. The commands and parameters executed are summarized in Tables S3-S21.

To measure the performance of our model, we adopted all metrics available in the TAXXI benchmark and extended them by adding standard machine learning metrics. Furthermore, we also computed hierarchical metrics, which can give better insights into which algorithm is better at classifying hierarchical data [21]. The definitions of all these metrics are listed in the supporting information text.

The datasets in the TAXXI benchmark were created using real data, with a strategy known as cross-validation by identity, where varying distances between query sequences and the reference database are accounted for. In practice, this means that in the dataset with 90% identity all species are novel and there is a mix of novel and known taxa at the genus level, making it the most difficult dataset in the benchmark.

As the identity increases, so does the amount of known labels at all taxonomic ranks; hence, the dataset with 100% identity can be considered the easiest in the benchmark.

#### 3 Results and discussion

First, we evaluated HiTaC on the dataset with 90% identity to assess its behavior under uncertainty. In order to do this, we computed the hierarchical precision to measure the proportion of correct predictions, the hierarchical recall to measure the proportion of detected positive samples and the hierarchical F1-score, which is the harmonic mean of the precision and recall. As shown in Figure 2, HiTaC_Filter was one of the top-performing methods in terms of precision, scoring 96.05 while maintaining a recall of 78.86, equivalent to other high-ranking methods. Although other software obtained similar or slightly better precision, it cost them a steeper decline in recall, which is also evidenced by the lower F1 scores. This result shows that HiTaC performed well in the presence of uncertainty.

**Fig. 2:**
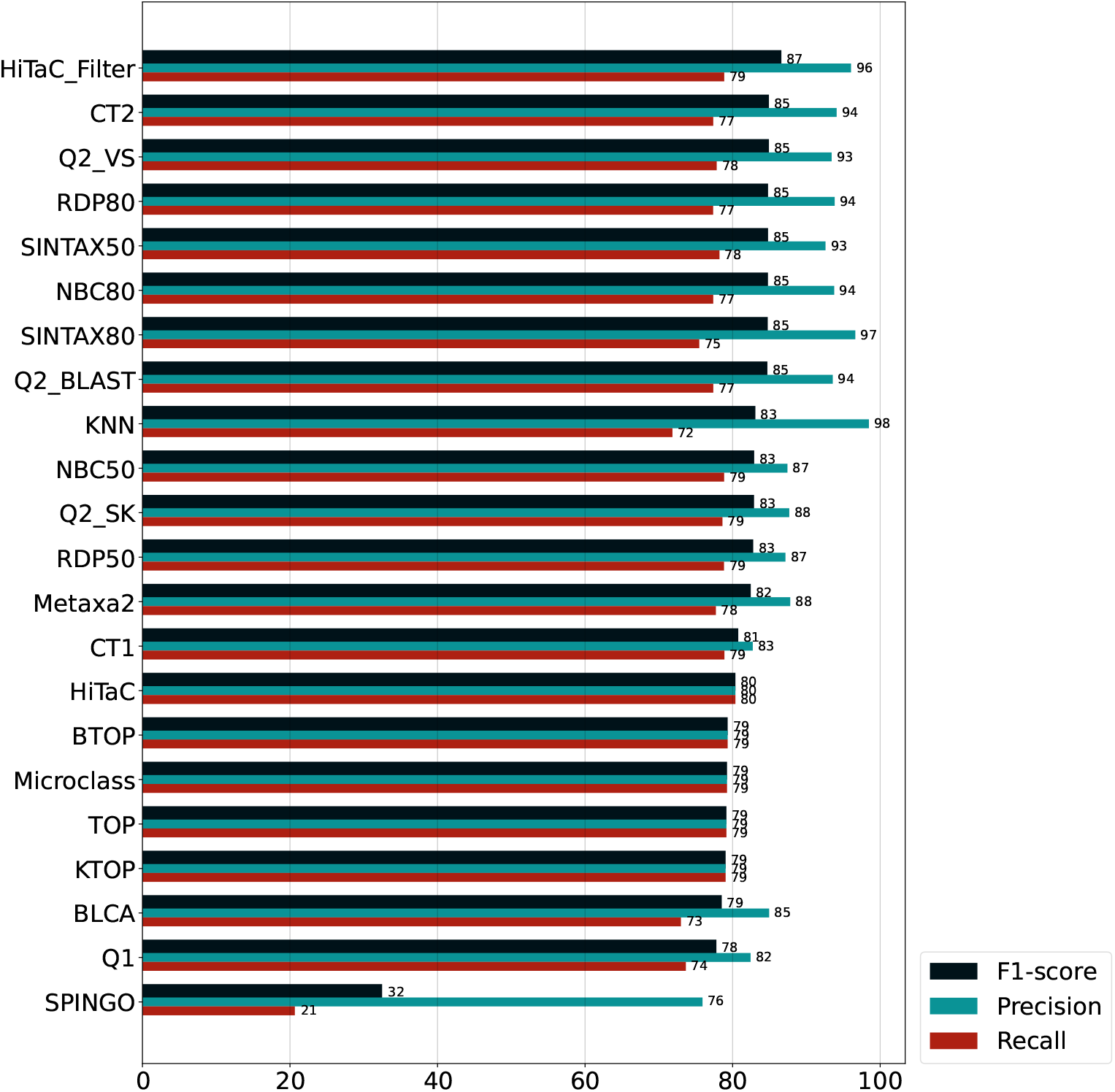
Hierarchical f1-score, precision and recall computed for all taxonomic ranks for the dataset with 90% identity, sorted by f1-score. HiTaC_Filter was one of the best-performing methods when dealing with uncertainty, achieving high f1-score, precision and recall, while the filter-less version obtained the highest recall of all methods.

To appraise HiTaC’s behavior when classifying known organisms, we evaluated it on the dataset with 100% identity. As shown in Figure 3, HiTaC_Filter achieved a perfect precision score of 100 when the taxonomy was fully annotated in the reference database. Furthermore, the recall persisted at 99.68; that is, there was no prominent decrease in recall when compared with the filter-less version. This result shows the potential of HiTaC in accurately classifying known organisms, given that query sequences have perfect matches in the reference database.

**Fig. 3:**
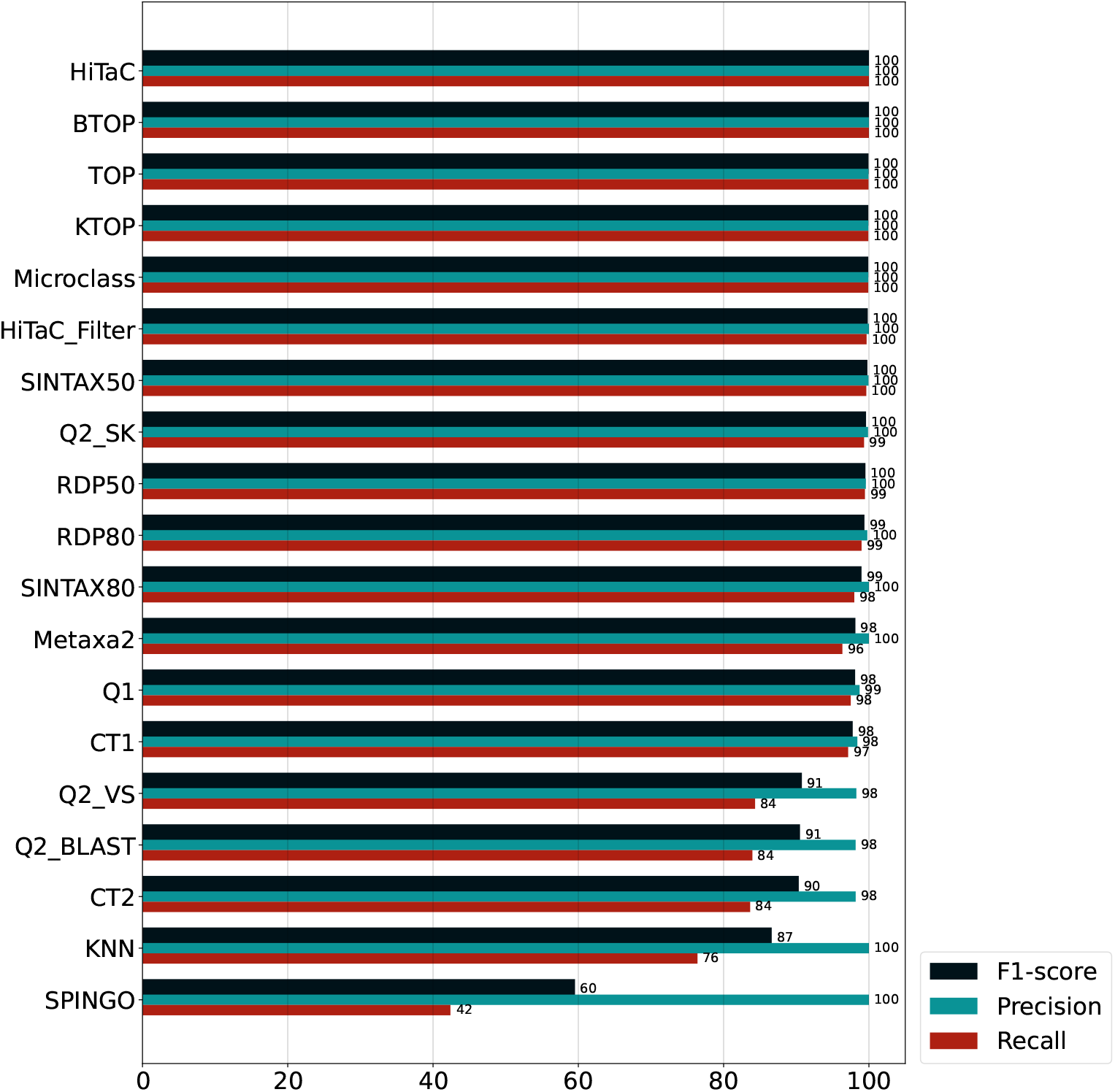
Hierarchical f1-score, precision and recall computed for all taxonomic ranks for the dataset with 100% identity. Both HiTaC and HiTaC_Filter achieved almost perfect scores when all organisms were known.

In Figure 4, we summarize the results achieved for the hierarchical metrics by computing the F1-score for all five datasets and sorting by the average of F1 scores. HiTaC_Filter obtained an F1-score equivalent or higher to the top-performing approaches for all datasets, corroborating the results presented in the previous paragraphs. This result demonstrates that HiTaC_Filter can positively identify organisms annotated in databases while being able to leave most new species unannotated.

**Fig. 4:**
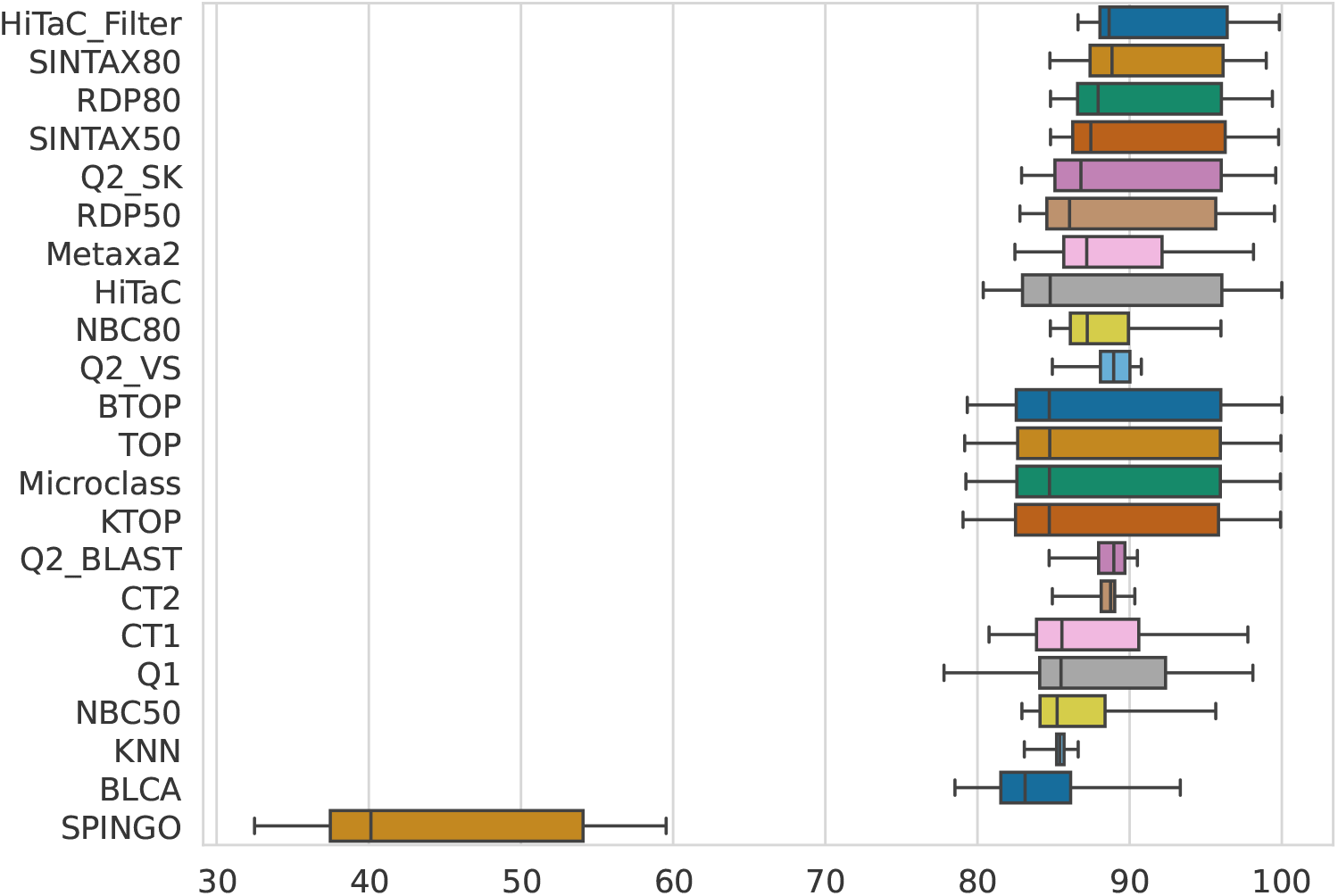
Hierarchical F1-score computed for all taxonomic ranks for the datasets with 90, 95, 97, 99 and 100% identity, sorted by average. HiTaC_Filter achieved F1-scores equal to or above the best methods available in the literature.

Similar conclusions can be drawn from Figure 5, in which we evaluated the percentage of known sequences correctly predicted, i.e., the true positive rate (TPR) for all methods and datasets available in the benchmark. The trend is that HiTaC achieved a TPR higher or equal to top methods. For instance, the best true positive rate at the genus level for the dataset with 90% identity was achieved by HiTaC, which sharply increased the TPR by 6.9 percentage points when compared with BTOP. Regarding the dataset with 95% identity at the genus level, HiTaC obtained a 1.7 percentage points improvement over Microclass, while for the dataset with 97% identity at the genus level, HiTaC surpassed Microclass with a 0.5 increase in percentage points. At the species level, HiTaC achieved a 0.7 percentage points gain over BTOP for the dataset with 99% identity, while for the dataset with 100% identity, HiTaC tied with BTOP, obtaining a perfect score of 100% TPR. These results suggest that the local hierarchical classification approach implemented in HiTaC commits fewer errors than flat classifiers as long as the sequences are previously known.

**Fig. 5:**
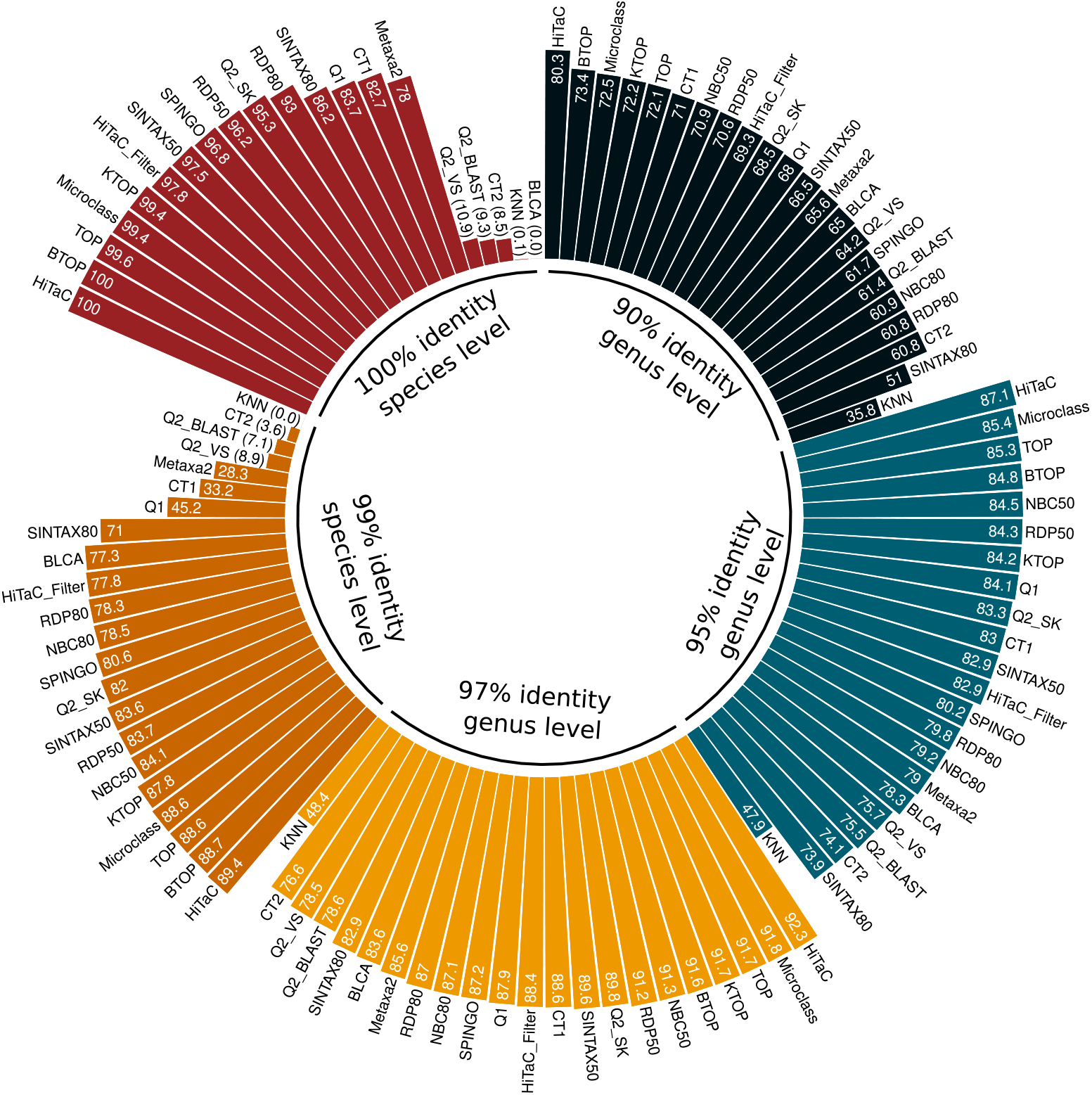
True positive rates for all methods for the datasets with 90%, 95% and 97% identity at the genus level, and 99% and 100% identity at the species level. The trend is that HiTaC achieved a sensitivity higher or equal to top methods. For instance, HiTaC tied with the best method for the dataset with 100% identity, obtaining a perfect score. Moreover, for the datasets with 90-99% identity, HiTaC improved the true positive rates upon top methods.

By default, HiTaC uses 6-mer frequency as a feature extraction method, but the user can change this parameter. Nevertheless, increasing the k-mer size provides small gains in predictive performance and comes with higher computational costs. The k-mer frequency computation method discards substrings with ambiguous nucleotides since they do not often occur in short reads, and accounting for them would require a slower algorithm. For example, a V means that there is either an A, C, or G in that position, and computing all these possibilities would make the method slower. Hence, we opted to disregard ambiguous nucleotides to keep the k-mer counting process as fast as possible. In future releases, we intend to implement a flexible user interface to allow users to use third-party feature extraction methods. Furthermore, we believe allowing for third-party feature extraction methods could also enable accurate taxonomic classification for 16S rRNA.

Training the filter is slower than most methods evaluated in the benchmark (Tables S22-S26). However, it must only be performed once for the reference database, and after training, the filter can be used to estimate the uncertainty quickly. Moreover, some public databases, such as UNITE and SILVA, are not updated frequently. Nevertheless, we provide models pre-trained on the public database UNITE^2^, speeding up the process for users. Some of these pre-trained models contain all eukaryotic ITS sequences available on UNITE, which enable detection and removal of nonfungal sequences mistakenly amplified by polymerase chain reaction (PCR) [26]. Furthermore, HiTaC uses the unique species hypotheses identifiers provided in the database UNITE as the last taxonomic level in the hierarchy during training and reports them to the user, which increases taxonomic reproducibility.

In summary, HiTaC is a Python package optimized to produce accurate taxonomic classification of fungal ITS sequences. Due to the extensive use of the taxonomic hierarchy during training, HiTaC produces fewer errors than other methods compared in the benchmark. Furthermore, its filter is a reliable classifier under uncertainty. HiTaC provides standalone Python scripts and a QIIME 2 plugin that can be quickly adopted into existing analysis pipelines. HiTaC and its dependencies can be easily installed via `pip, conda` or `docker`, which is not the case for most of the taxonomic classifiers available in the literature. HiTaC is released under the new BSD license, allowing free use in academia and industry and free copy, modification, and redistribution of the code as long as a duplicate of the original license is kept and proper acknowledgments are given.

## 4 Conclusion

HiTaC is a Python package for the taxonomic classification of fungal ITS sequences, which applies local hierarchical classifiers for accurate predictions. It provides scripts to train classifiers on specialized datasets and to classify new sequences with the trained models. The standard version has high sensitivity and is ideal for exploratory analysis. HiTaC also implements a filter that can express uncertainties in classifications, indicating if the input sequences are complex to recognize. Thanks to its compatibility with QIIME 2, users can quickly adopt it in existing mycobiome analysis pipelines. HiTaC is an open-source software available at https://gitlab.com/dacs-hpi/hitac.

## Availability and requirements

- **Project name:** HiTaC
- **Project home page:** https://gitlab.com/dacs-hpi/hitac
- **Operating system(s):** Platform independent
- **Programming language:** Python
- **Other requirements:** HiClass v4.4.0 or higher, scikit-learn v1.3.0 or higher
- **License:** new BSD
- **Any restrictions to use by non-academics:** Not applicable

## Supplementary information. Additional file 1

Supporting information text, figures S1-S6 and tables S1-S91.

## Acknowledgements

FMM gratefully acknowledges Marcos Augusto dos Santos (UFMG) for initial inspiration, Robert C. Edgar (independent researcher) for providing the scripts and datasets in the TAXXI benchmark, Sven Giese (HPI) for code improvement suggestions and Elizabeth Yuu (HPI) for proofreading.

## Declarations

### Ethics approval and consent to participate

Not applicable.

### Consent for publication

Not applicable.

### Availability of data and materials

HiTaC is freely available on the Python package index, BIOCONDA and `docker` hub. It is easier to install it by executing `pip install hitac` in the terminal. All datasets and evaluation scripts used in this paper are publicly available in the TAXXI benchmark [25], and a reproducible evaluation pipeline is available on GitLab https://gitlab.com/dacs-hpi/hitac/-/tree/main/benchmark.

### Authors’ contributions

FMM designed and implemented HiTaC’s algorithm, performed analysis, benchmarked the experiments and wrote the manuscript. VAA and RJR assisted with analysis and revised the manuscript. BYR assisted with analysis, visualizations and reviewed the manuscript. VCP oversaw experimental and analysis aspects of the project and reviewed the manuscript. All authors edited and approved the final manuscript.

## Footnotes

1 https://gitlab.com/dacs-hpi/hitac

2 https://gitlab.com/dacs-hpi/hitac#pre-trained-models

