## Supplementary Material for "HiTaC: a hierarchical taxonomic classifier for fungal ITS sequences compatible with QIIME2"

bioRχiv

S21 Commands and parameters to run TOP. 33
S22 Resources benchmark computed for the dataset SP RDP ITS 90. 34
S23 Resources benchmark computed for the dataset SP RDP ITS 95. 35
S24 Resources benchmark computed for the dataset SP RDP ITS 97. 36
S25 Resources benchmark computed for the dataset SP RDP ITS 99. 37
S26 Resources benchmark computed for the dataset SP RDP ITS 100. 38
S27 Hierarchical metrics computed for the dataset SP RDP ITS 90. 39
S28 Hierarchical metrics computed for the dataset SP RDP ITS 95. 40
S29 Hierarchical metrics computed for the dataset SP RDP ITS 97. 41
S30 Hierarchical metrics computed for the dataset SP RDP ITS 99. 42
S31 Hierarchical metrics computed for the dataset SP RDP ITS 100. 43
S32 TAXXI metrics computed for the dataset SP RDP ITS 90 at the phylum level. 44
S33 TAXXI metrics computed for the dataset SP RDP ITS 90 at the class level. 45
S34 TAXXI metrics computed for the dataset SP RDP ITS 90 at the order level. 46
S35 TAXXI metrics computed for the dataset SP RDP ITS 90 at the family level. 47
S36 TAXXI metrics computed for the dataset SP RDP ITS 90 at the genus level. 48
S37 TAXXI metrics computed for the dataset SP RDP ITS 90 at the species level. 49
S38 TAXXI metrics computed for the dataset SP RDP ITS 95 at the phylum level. 50
S39 TAXXI metrics computed for the dataset SP RDP ITS 95 at the class level. 51
S40 TAXXI metrics computed for the dataset SP RDP ITS 95 at the order level. 52
S41 TAXXI metrics computed for the dataset SP RDP ITS 95 at the family level. 53
S42 TAXXI metrics computed for the dataset SP RDP ITS 95 at the genus level. 54
S43 TAXXI metrics computed for the dataset SP RDP ITS 95 at the species level. 55
S44 TAXXI metrics computed for the dataset SP RDP ITS 97 at the phylum level. 56
S45 TAXXI metrics computed for the dataset SP RDP ITS 97 at the class level. 57
S46 TAXXI metrics computed for the dataset SP RDP ITS 97 at the order level. 58
S47 TAXXI metrics computed for the dataset SP RDP ITS 97 at the family level. 59
S48 TAXXI metrics computed for the dataset SP RDP ITS 97 at the genus level. 60
S49 TAXXI metrics computed for the dataset SP RDP ITS 97 at the species level. 61
S50 TAXXI metrics computed for the dataset SP RDP ITS 99 at the phylum level. 62
S51 TAXXI metrics computed for the dataset SP RDP ITS 99 at the class level. 63
S52 TAXXI metrics computed for the dataset SP RDP ITS 99 at the order level. 64
S53 TAXXI metrics computed for the dataset SP RDP ITS 99 at the family level. 65
S54 TAXXI metrics computed for the dataset SP RDP ITS 99 at the genus level. 66
S55 TAXXI metrics computed for the dataset SP RDP ITS 99 at the species level. 67
S56 TAXXI metrics computed for the dataset SP RDP ITS 100 at the phylum level. 68
S57 TAXXI metrics computed for the dataset SP RDP ITS 100 at the class level. 69
S58 TAXXI metrics computed for the dataset SP RDP ITS 100 at the order level. 70
S59 TAXXI metrics computed for the dataset SP RDP ITS 100 at the family level. 71
S60 TAXXI metrics computed for the dataset SP RDP ITS 100 at the genus level. 72
S61 TAXXI metrics computed for the dataset SP RDP ITS 100 at the species level. 73
S62 Machine learning metrics computed for the dataset SP RDP ITS 90 at the phylum level. 74
S63 Machine learning metrics computed for the dataset SP RDP ITS 90 at the class level. 75
S64 Machine learning metrics computed for the dataset SP RDP ITS 90 at the order level. 76
S65 Machine learning metrics computed for the dataset SP RDP ITS 90 at the family level. 77
S66 Machine learning metrics computed for the dataset SP RDP ITS 90 at the genus level. 78
S67 Machine learning metrics computed for the dataset SP RDP ITS 90 at the species level. 79
S68 Machine learning metrics computed for the dataset SP RDP ITS 95 at the phylum level. 80
S69 Machine learning metrics computed for the dataset SP RDP ITS 95 at the class level. 81
S70 Machine learning metrics computed for the dataset SP RDP ITS 95 at the order level. 82
S71 Machine learning metrics computed for the dataset SP RDP ITS 95 at the family level. 83
S72 Machine learning metrics computed for the dataset SP RDP ITS 95 at the genus level. 84
S73 Machine learning metrics computed for the dataset SP RDP ITS 95 at the species level. 85
S74 Machine learning metrics computed for the dataset SP RDP ITS 97 at the phylum level. 86
S75 Machine learning metrics computed for the dataset SP RDP ITS 97 at the class level. 87
S76 Machine learning metrics computed for the dataset SP RDP ITS 97 at the order level. 88
S77 Machine learning metrics computed for the dataset SP RDP ITS 97 at the family level. 89
S78 Machine learning metrics computed for the dataset SP RDP ITS 97 at the genus level. 90
S79 Machine learning metrics computed for the dataset SP RDP ITS 97 at the species level. 91
S80 Machine learning metrics computed for the dataset SP RDP ITS 99 at the phylum level. 92
S81 Machine learning metrics computed for the dataset SP RDP ITS 99 at the class level. 93

S82 Machine learning metrics computed for the dataset SP RDP ITS 99 at the order level. . . . . . . . . . . . . . . . . 94
S83 Machine learning metrics computed for the dataset SP RDP ITS 99 at the family level. . . . . . . . . . . . . . . 95
S84 Machine learning metrics computed for the dataset SP RDP ITS 99 at the genus level. . . . . . . . . . . . . . . . 96
S85 Machine learning metrics computed for the dataset SP RDP ITS 99 at the species level. . . . . . . . . . . . . . 97
S86 Machine learning metrics computed for the dataset SP RDP ITS 100 at the phylum level. . . . . . . . . . . . . 98
S87 Machine learning metrics computed for the dataset SP RDP ITS 100 at the class level. . . . . . . . . . . . . . . 99
S88 Machine learning metrics computed for the dataset SP RDP ITS 100 at the order level. . . . . . . . . . . . . . 100
S89 Machine learning metrics computed for the dataset SP RDP ITS 100 at the family level. . . . . . . . . . . . . . 101
S90 Machine learning metrics computed for the dataset SP RDP ITS 100 at the genus level. . . . . . . . . . . . . . 102
S91 Machine learning metrics computed for the dataset SP RDP ITS 100 at the species level. . . . . . . . . . . . . 103

#### Supporting Information Text

#### Evaluation

The TAXXI benchmark (1) was employed to facilitate the comparison between HiTaC and seventeen other similar software previously available in the benchmark (Table S1), some of which are similarity based approaches, while others are alignment free. We relied on this benchmark because it mimics more accurately real world scenarios including novel sequences, as well as its diversity in software, reference databases and use of real environmental data. Furthermore, the TAXXI benchmark assesses the accuracy using cross-validation by identity, a strategy which models the variation in distances between query sequences and the closest entry in reference databases. In this approach, a 90% identity enforces sequences to only belong to species that are absent from the training set, while a 95% identity allows for a mix of present and absent species. In other words, the cross-validation by identity enables the creation of datasets whose lowest taxonomic ranks were nonexistent in the training data while still knowing the ground truth, which allows for a better evaluation on how a classifier behaves when trying to predict novel sequences.

**Datasets.** We adopted five datasets containing both training and test data, which were previously available in the TAXXI benchmark (1). These datasets were selected because they were the only ones with taxonomic annotations of fungal ITS sequences up to species level. Although there were other five datasets available in this benchmark possessing fungal ITS sequences, we noticed that they were copies of the same datasets we selected, except that they were missing the species level annotation in the training datasets and ground truth. Thus, there was no added value in using these other datasets in our evaluation.

The fungal ITS datasets selected from the TAXXI benchmark were based on real environmental data, publicly available on NCBI (2), RDP (3), the Warcup fungal ITS training set v2 (WITS) (4), UNITE (5) and other *in vivo* samples. They were created through cross-validation by identity, hence the identity between the query sequences and the closest entry in a reference database varied on them, ranging from 90% to 100% (1) (Table S2). In practice, an identity of 90% means that the training data does not contain any of the labels that are in the test dataset at the species level, but most labels are known at the genus level. The known taxa at the species level increase with identity, i.e., all species are present in the training data when the identity is 100%.

It is also important to mention that the training datasets are extremely imbalanced. For example, among the seven existing phyla on dataset SP RDP ITS 90, Ascomycota and Basidiomycota contain approximately 52% and 44% of total instances, respectively (Fig. S1). A similar pattern can be observed in the other datasets with higher identities.

##### Evaluation Metrics.

***Default metrics from the TAXXI benchmark.*** These metrics, as well as scripts used to compute them, originated from the TAXXI benchmark (1):

- **Accuracy (ACC) -** Accounts for all test sequences predicted by the algorithm with names that are known and/or over-predicted, ranging from 0% (no correct predictions) to 100% (no errors);
- **Misclassification Rate (MCR) -** The percentage of known sequences incorrectly predicted;
- **Over-classification Rate (OCR) -** The percentage of unknown sequences assigned a label;
- **True Positive Rate (TPR) -** The percentage of known sequences correctly predicted;
- **Under-classification Rate (UCR) -** The percentage of known sequences that were not assigned a label.

Defining these metrics in mathematical terms, we have:

$$\text{ACC} = 100 \times \frac{\text{True Positive}}{\text{Known Sequences} + \text{Over-classification Errors}}$$

$$\text{MCR} = 100 \times \frac{\text{Misclassification Errors}}{\text{Known Sequences}}$$

$$\text{OCR} = 100 \times \frac{\text{Over-classification Errors}}{\text{Novel Sequences}}$$

$$\text{TPR} = 100 \times \frac{\text{True Positive}}{\text{Known Sequences}}$$

$$\text{UCR} = 100 \times \frac{\text{Under-classification Errors}}{\text{Known Sequences}}$$

where *known sequences* have labels in the training dataset and *novel sequences* do not have labels in the training dataset. A *true positive* is the number of correctly predicted sequences. *Under-classification errors* occur when too few ranks are predicted, while *over-classification errors* happen when too many ranks are predicted. Finally, *misclassification errors* occur when a known name is incorrectly predicted.

***Standard machine learning metrics.*** These metrics and their respective implementations originated from the package scikit-learn (6):

- **Accuracy -** The fraction of correctly classified samples;
- **Balanced accuracy -** Accuracy counterpart planned to deal with imbalanced datasets, which is defined as the average of recall obtained on each class;
- **Precision -** Intuitively, precision is the ability of the classifier not to label a negative sample as positive, and it ranges from 0 (worst score) to 100 (best value);
- **Recall -** Intuitively, recall is the ability of the classifier to detect all positive samples;
- **F1-score -** The F1-score is the harmonic mean of the precision and recall, ranging from 0 (worst score) to 100 (best value);
- **Jaccard -** Quantifies the similarity between the set of predicted labels and the ground truth;
- **Micro, macro and weighted variants -** The Precision, Recall, F1-score and Jaccard were computed in three different ways, which are described below:

  - **Micro:** Computes metrics globally considering the overall true positives, false negatives and false positives;

  - **Macro:** Computes metrics locally for each label and determines their unweighted average without considering label imbalance;

  - **Weighted:** Computes metrics locally for each label and determines their average weight by support, that is, the total of correct instances for each label. In practice, this modifies "macro" to consider label imbalance, which can result in an F1-score that is not between precision and recall.

Defining these metrics in mathematical terms, we have:

$$\text{Accuracy}(y, \hat{y}) = 100 \times \frac{1}{n_\text{samples}} \sum_{i=0}^{n_{samples}-1} \mathbb{1}(\hat{y}_i = y_i)$$

where $\hat{y}_i$ is the predicted value of the $i$-th sample and $y_i$ is the corresponding ground truth, while $\mathbb{1}$ is the indicator function.

$$\text{Balanced accuracy}(y, \hat{y}, w) = 100 \times \frac{1}{\sum \hat{w}_i} \sum_i \mathbb{1}(\hat{y}_i = y_i)\hat{w}_i$$

where $\hat{w}_i = \dfrac{w_i}{\sum_j \mathbb{1}(y_j = y_i) w_j}$ and $w_i$ is the weight of the $i$-th sample.

$$\text{Precision}(y, \hat{y}) = 100 \times \frac{\text{True Positives}}{\text{True Positives} + \text{False Positives}}$$

$$\text{Recall}(y, \hat{y}) = 100 \times \frac{\text{True Positives}}{\text{True Positives} + \text{False Negatives}}$$

$$\text{F1-score}(y, \hat{y}) = 100 \times \frac{2 \times \text{Precision} \times \text{Recall}}{\text{Precision} + \text{Recall}}$$

$$\text{Jaccard}(y, \hat{y}) = 100 \times \frac{y \cap \hat{y}}{y \cup \hat{y}}$$

$$\text{Micro}(y, \hat{y}) = 100 \times M(y, \hat{y})$$

where $M$ is the metric to compute for, i.e., precision, recall, F1-score or jaccard.

$$\text{Macro}(y, \hat{y}) = 100 \times \frac{1}{|L|} \sum_{l \in L} M(y_l, \hat{y}_l)$$

where $L$ is the set of labels.

$$\text{Weighted}(y, \hat{y}) = 100 \times \frac{1}{\sum_{l \in L} |y_l|} \sum_{l \in L} |y_l| M(y_l, \hat{y}_l)$$

***Hierarchical machine learning metrics.*** These metrics are adaptations of the previously introduced metrics of precision, recall and F1-score, but tailored to the hierarchical classification scenario. Their respective implementations originated from the package HiClass (7):

$$\text{Hierarchical precision } (hP) = 100 \times \frac{\sum_i |\alpha_i \cap \beta_i|}{\sum_i |\alpha_i|}$$

$$\text{Hierarchical recall } (hR) = 100 \times \frac{\sum_i |\alpha_i \cap \beta_i|}{\sum_i |\beta_i|}$$

$$\text{Hierarchical F1-score} = 100 \times \frac{2 \times hP \times hR}{hP + hR}$$

where $\alpha_i$ is the set containing the predicted labels for test example $i$ and their respective ancestors, while $\beta_i$ is the set containing the ground truth of test example $i$ and their respective ancestors, with summations computed over all test examples.

**Resources benchmark.** We enhanced the TAXXI benchmark by porting it to Snakemake (8) in order to make it more reproducible, and we introduced a resources benchmark. The benchmark scripts and instructions on how to run it are available in our GitLab repository[1]. The benchmark was computed on a laptop running GNU/Linux with 16 GB physical memory and 12 cores provided by an AMD Ryzen™ 5 processor, where 12 cores were allocated for each method. The results from the benchmark are available in Tables S22-S26, while the commands and parameters executed for all software are available in Tables S3-S21.

[1] https://gitlab.com/dacs-hpi/hitac/-/tree/main/benchmark

### Number of examples for each taxonomic category

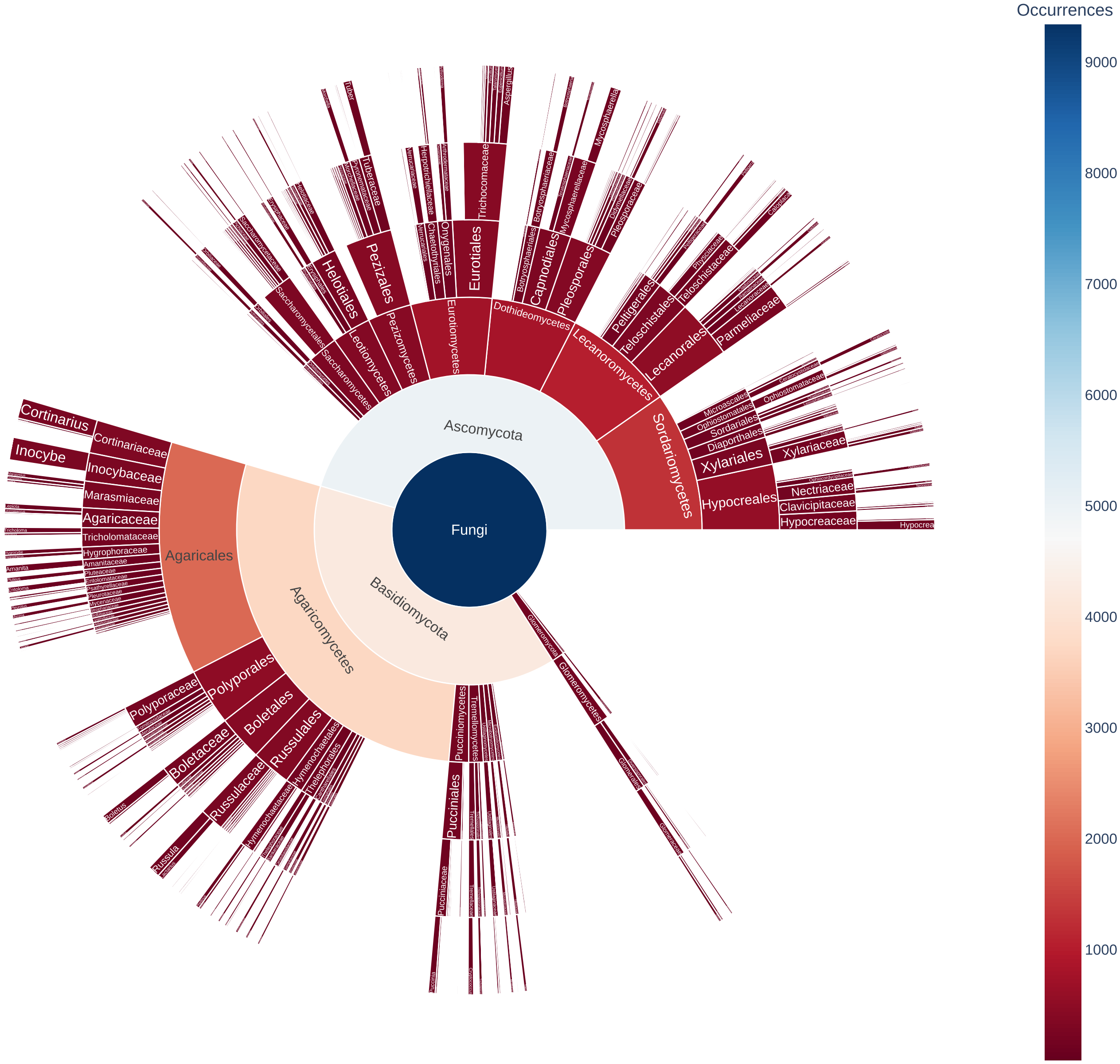

**Fig. S1.** Depiction of imbalanced training set. For instance, the phyla Ascomycota and Basidiomycota together contain approximately 96% of total instances between the seven existing phyla in dataset SP RDP ITS 90. In the remaining taxonomic ranks and datasets we can also observe similar patterns.

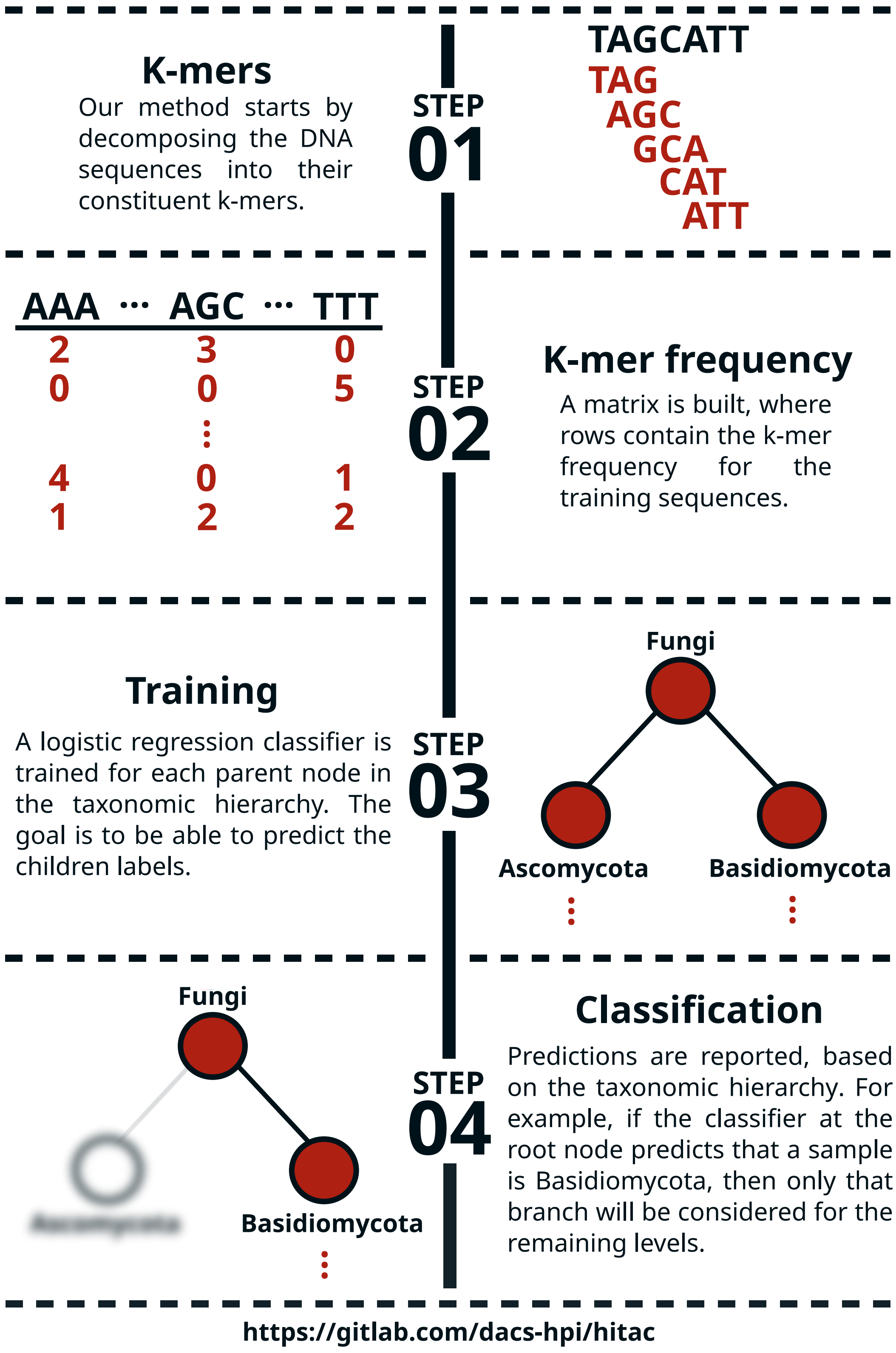

**Fig. S2.** HiTaC algorithm overview. To compute the k-mer frequency of DNA sequences, we first decompose them into their constituent k-mers. In the example shown in step 01, $k = 3$, resulting in 5 sub-strings. Afterwards, HiTaC computes a matrix containing the k-mer frequency for all training sequences, where each row contains the k-mer frequency for a given sequence and the columns are all possible k-mers for a given $k$, i.e., all $k$ nucleotide permutations of A, C, G and T. Next, HiTaC trains a logistic regression classifier for each parent node in the taxonomic tree, where the features are the k-mer frequencies computed in step 2 and the labels are the taxonomic annotations of the training sequences. In step 03, only the first 2 levels are shown for brevity. Lastly, HiTaC makes predictions in a top-down approach, based on the taxonomic hierarchy.

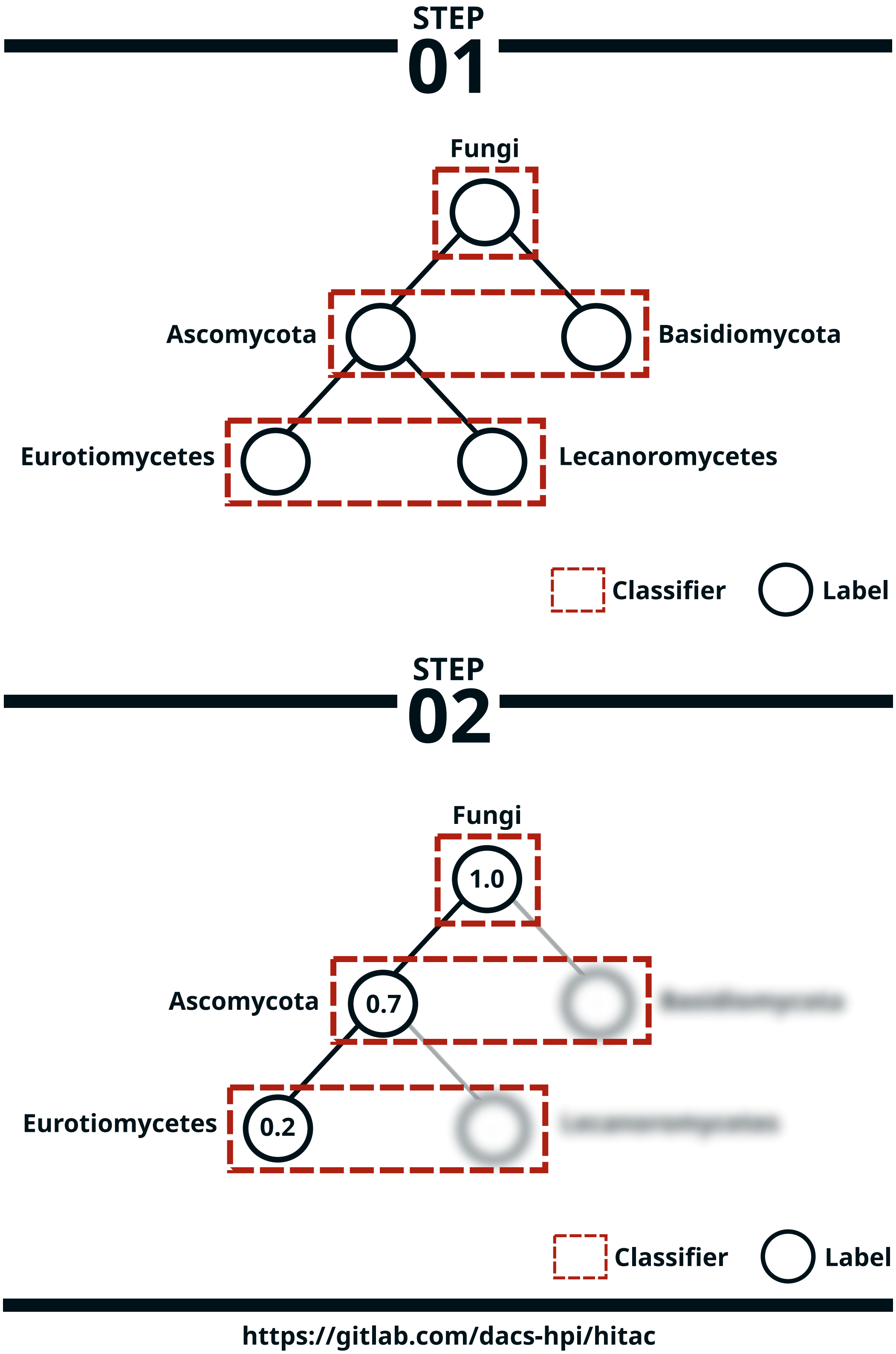

**Fig. S3.** HiTaC filter overview. To build the filter, we first train a logistic regression classifier for each level of the hierarchy, using the same k-mer frequency algorithm introduced previously on Figure S2. In the example shown in step 01, there are only 3 levels in the hierarchy for simplicity. Afterwards, HiTaC computes the confidence score for all taxonomic ranks assigned to a sample by the local classifier per parent node, then on a bottom-up approach it removes labels that are below a certain threshold. In step 02, supposing that the threshold is 0.7 and that the given sample has a confidence score of 0.2 assigned for the lowest rank, then the final prediction for this sample will only contain the labels Fungi and Ascomycota.

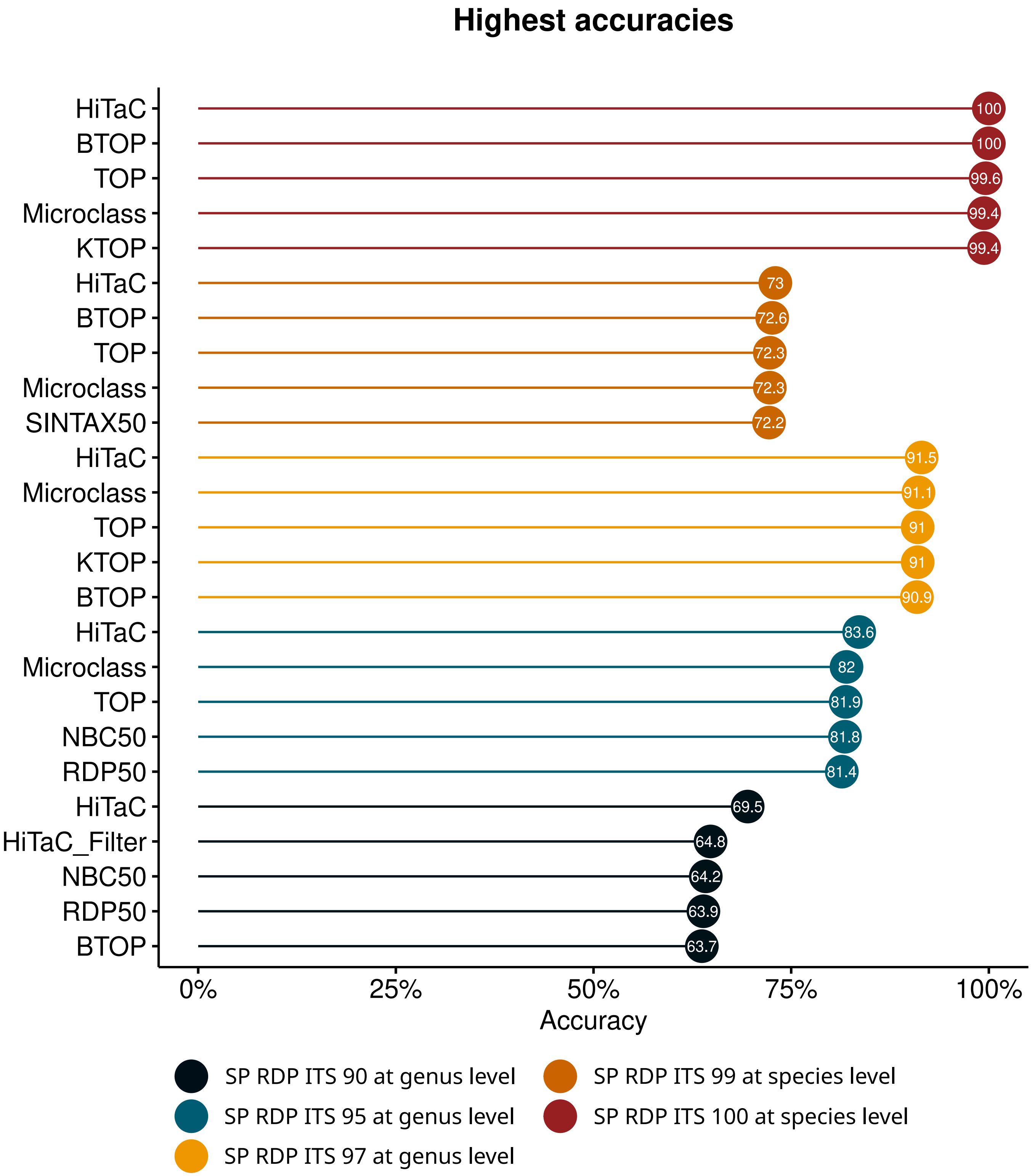

**Fig. S4.** Accuracies for the top 5 methods for datasets SP RDP ITS 90 at the genus level, SP RDP ITS 95 at the genus level, SP RDP ITS 97 at the genus level, SP RDP ITS 99 at the species level and SP RDP ITS 100 at the species level. The trend is that HiTaC achieved an accuracy higher or equal to top methods. For instance, HiTaC tied with the top method for the dataset with 100% identity, obtaining a perfect score. Moreover, for the datasets with 90-99% identity, HiTaC improved the accuracy upon top methods.

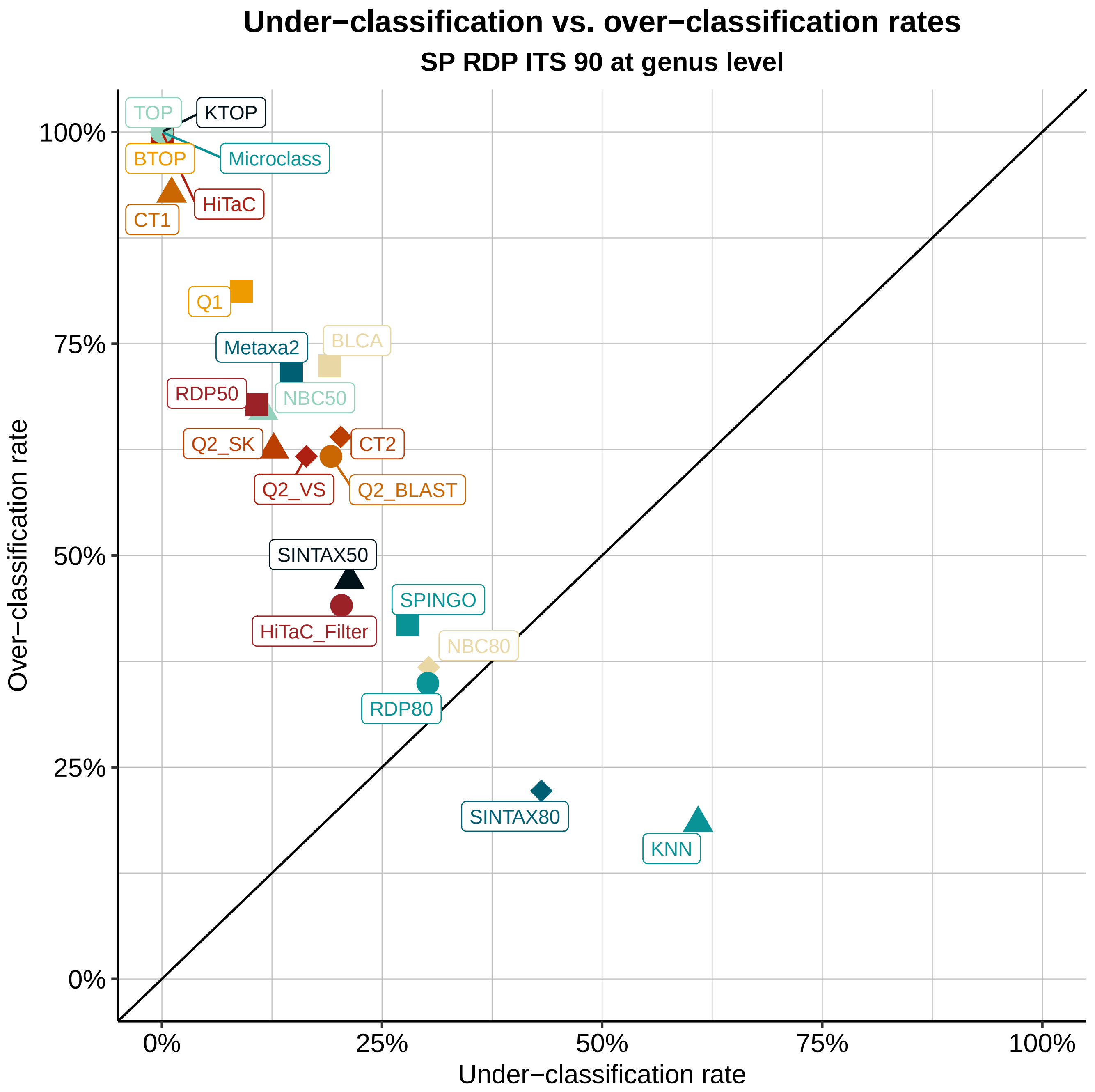

**Fig. S5.** Under-classification versus over-classification rates for dataset SP RDP ITS 90 at genus level. The introduction of the filter in HiTaC proved to be effective by sharply decreasing the over-classification rate from 100% to 44.1%. Contrarily, the filter slightly increased the under-classification rate from 0% to 20.4%, but it still ranked among the methods with the best trade-off between over-classification and under-classification rates, which are the methods closer to the diagonal on the bottom left. An ideal method should be able to classify all known sequences and leave novel sequences unclassified, thus avoiding both over-classification and under-classification errors.

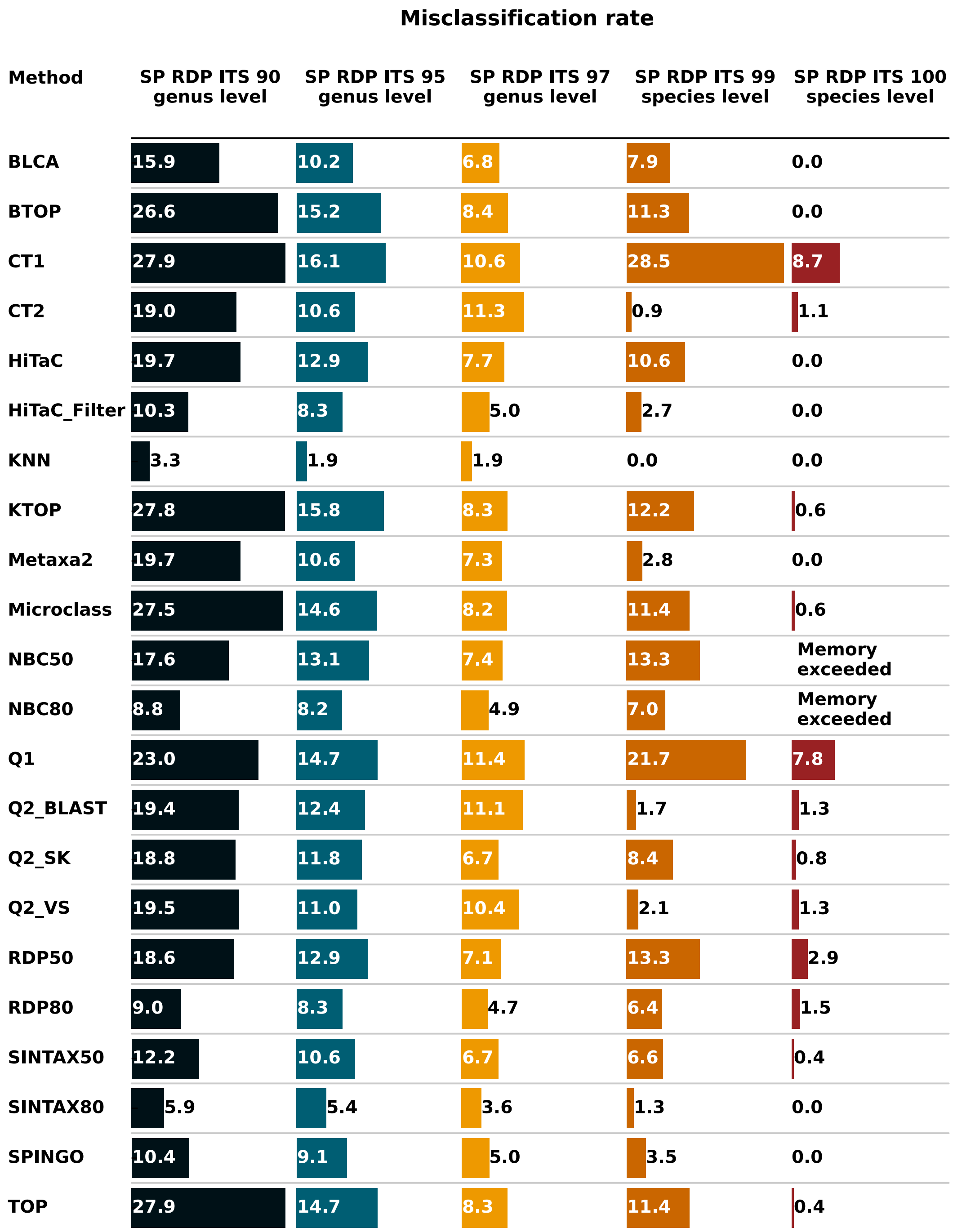

**Fig. S6.** Misclassification rate results for the SP RDP ITS datasets at genus and species level. The misclassification rates for HiTaC, HiTaC_Filter, BTOP, BLCA, SPINGO, SINTAX80, Metaxa2 and KNN were 0% for the dataset SP RDP ITS 100. HiTaC's misclassification rate can be deemed adequate for the datasets with 90–99% identity, while misclassification rate reduced even further when the filter was applied to HiTaC. NBC50 and NBC80 crashed with memory limit exceeded for dataset SP RDP ITS 100, so their results are missing.

**Table S1. List of software available in the TAXXI benchmark (1), which were used for comparison with HiTaC.**

| Method | Software | Algorithm reference |
|---|---|---|
| BLCA | BLCA v2.3-alpha | (9) |
| BTOP | blastn v2.10.1+ | (10) |
| CT1 | usearch v11.0.667 | Re-implements Q1 (1) |
| CT2 | usearch v11.0.667 | Re-implements Q2_VS (1) |
| HiTaC | HiTaC v2.2.2 | (This paper) |
| KNN | mothur v1.48.0 | (11) |
| KTOP | usearch v11.0.667 | (1) |
| Metaxa2 | Metaxa 2.2.3 | (12) |
| Microclass | microclass v1.2 | (13) |
| NBC | usearch v11.0.667 | Re-implements RDP (1) |
| Q1 | QIIME v1.9 | (14) |
| Q2_BLAST | QIIME v2.2022.2 | (15) |
| Q2_SK | QIIME v2.2022.2 | (15) |
| Q2_VS | QIIME v2.2022.2 | (15) |
| RDP | Classifier v2.13 | (16) |
| SINTAX | usearch v11.0.667 | (17) |
| SPINGO | SPINGO v1.3 | (18) |
| TOP | usearch v11.0.667 | (1) |

**Table S2. List of datasets available in the TAXXI benchmark (1), which were selected for possessing taxonomic annotations of fungal ITS sequences up to species level and were used to compare HiTaC with similar software.**

| Name | Lowest Rank | Identity (%) | # Training Sequences | # Test Sequences |
|---|---|---|---|---|
| SP RDP ITS 90 | Species | 90 | 9336 | 1935 |
| SP RDP ITS 95 | Species | 95 | 11439 | 1265 |
| SP RDP ITS 97 | Species | 97 | 12545 | 1420 |
| SP RDP ITS 99 | Species | 99 | 10901 | 3945 |
| SP RDP ITS 100 | Species | 100 | 16055 | 16055 |

**Table S3. Commands and parameters to run BLCA.**

```
python scripts/fasta_utax2_to_blca.py \
    {input.train} \
    {output.reference_reads} \
    {output.reference_taxonomy}

python scripts/fasta_utax2_to_blca.py \
    {input.test} \
    {output.query_reads} \
    {output.query_taxonomy}

makeblastdb \
    -in {input.reference_reads} \
    -dbtype nucl \
    -parse_seqids \
    -out {params.database}

2.blca_main.py \
    -i {input.query_reads} \
    -r {input.reference_taxonomy} \
    -q {params.database} \
    --proc {threads}

python scripts/blca2tab.py \
    {input.predictions} \
    {input.test} \
    > {output.predictions}
```

**Table S4. Commands and parameters to run BTOP.**

```
sed \
    "-es/;.*//" \
    < {input.train} \
    > {output.reference}

makeblastdb \
    -in {input.reference} \
    -dbtype nucl \
    -parse_seqids \
    -out {params.database}

blastn \
    -task megablast \
    -db {params.database} \
    -query {input.test} \
    -num_threads {threads} \
    -max_target_seqs 1 \
    -outfmt "6 qseqid sseqid" \
    -evalue 0.01 \
    > {output.predictions}

python scripts/btop2tab.py \
    {input.predictions} \
    {input.train} \
    {output.predictions}
```

**Table S5. Commands and parameters to run CT1.**

```
usearch \
    -db {input.train} \
    -cons_tax {input.test} \
    -strand plus \
    -tabbedout \
    {output.predictions} \
    -strand plus \
    -id 0.7 \
    -maxaccepts 3 \
    -maxrejects 8 \
    -maj 0.51
```

**Table S6. Commands and parameters to run CT2.**

```
usearch \
    -db {input.train} \
    -cons_tax {input.test} \
    -strand plus \
    -tabbedout \
    {output.predictions} \
    -strand plus \
    -id 0.7 \
    -maxaccepts 10 \
    -maxrejects 32 \
    -maj 0.51
```

**Table S7. Commands and parameters to run HiTaC.**

```
qiime hitac fit \
    --i-reference-reads {input.reference_reads} \
    --i-reference-taxonomy {input.reference_taxonomy} \
    --p-kmer 6 \
    --p-threads {threads} \
    --o-classifier {output.classifier}

qiime hitac classify \
    --i-reads {input.query_reads} \
    --i-classifier {output.classifier} \
    --p-kmer 6 \
    --p-threads {threads} \
    --o-classification {output.predictions}
```

**Table S8. Commands and parameters to run HiTaC_Filter.**

```
qiime hitac fit-filter \
    --i-reference-reads {input.reference_reads} \
    --i-reference-taxonomy {input.reference_taxonomy} \
    --p-kmer 6 \
    --p-threads {threads} \
    --o-filter {output.filter}

qiime hitac filter \
    --i-filter {output.filter} \
    --i-reads {input.query_reads} \
    --i-classification {input.unfiltered_predictions} \
    --p-threshold 0.7 \
    --p-kmer 6 \
    --p-threads {threads} \
    --o-filtered-classification {output.filtered_predictions}
```

**Table S9. Commands and parameters to run KNN.**

```
cp {input.test} {output.query_reads}

python2 scripts/mothur_make_taxtrainfiles.py \
    {input.train} \
    {output.reference_reads} \
    {output.reference_taxonomy}

mothur \
    "#classify.seqs(fasta={input.query_reads}, \
    template={input.reference_reads}, \
    taxonomy={input.reference_taxonomy}, \
    method=knn, processors={threads})"

python scripts/motknn2utax2.py \
    {input.predictions} \
    > {output.predictions}
```

**Table S10. Commands and parameters to run KTOP.**

```
usearch \
    -db {input.train} \
    -sintax {input.test} \
    -strand plus \
    -tabbedout \
    {output.predictions} \
    -strand plus \
    -ktop
```

 

**Table S11. Commands and parameters to run Metaxa2.**

```
python scripts/fasta_utax2_to_metaxa2.py \
    {input.train} \
    {output.reference_reads} \
    {output.reference_taxonomy}

metaxa2_dbb \
    -b {input.reference_reads} \
    -o {params.database} \
    -t {input.reference_taxonomy} \
    --auto_rep T \
    --cpu {threads} \
    --mode divergent

metaxa2 \
    -i {input.test} \
    -d {params.blast} \
    -p {params.hhms} \
    -o {params.predictions} \
    -cpu {threads}

python scripts/metaxa2tab.py \
    {input.predictions} \
    {input.test} \
    > {output.predictions}
```

**Table S12. Commands and parameters to run Microclass.**

```
Rscript scripts/microclass.R \
    {input.train} \
    {input.test} \
    {output.predictions}

python scripts/microclass2tab.py \
    {input.predictions} \
    {output.predictions}
```

**Table S13. Commands and parameters to run NBC.**

```
usearch \
    -nbc_tax \
    {input.test} \
    -db {input.train} \
    -strand plus \
    -tabbedout {output.predictions}

python scripts/bbc_cutoff.py \
    {input.predictions} \
    0.5 \
    > {output.predictions}

python scripts/bbc_cutoff.py \
    {input.predictions} \
    0.8 \
    > {output.predictions}
```

**Table S14. Commands and parameters to run Q1.**

```
python scripts/fasta_utax2qiime.py \
    {input.train} \
    {output.reference_reads} \
    {output.reference_taxonomy}

assign_taxonomy.py \
    -i {input.test} \
    -m uclust \
    -r {input.reference_reads} \
    -t {input.reference_taxonomy} \
    -o {params.tmpdir}

python scripts/qiimetax2tab.py \
    {input.predictions} \
    > {output.predictions}
```

**Table S15. Commands and parameters to run Q2_BLAST.**

```
qiime feature-classifier classify-consensus-blast \
    --i-query {input.query_reads} \
    --i-reference-reads {input.reference_reads} \
    --i-reference-taxonomy {input.reference_taxonomy} \
    --o-classification {output.predictions} \
    --o-search-results {output.search_results}
```

**Table S16. Commands and parameters to run Q2_SK.**

```
qiime feature-classifier fit-classifier-naive-bayes \
    --i-reference-reads {input.reference_reads} \
    --i-reference-taxonomy {input.reference_taxonomy} \
    --o-classifier {output.classifier}

qiime feature-classifier classify-sklearn \
    --i-classifier {output.classifier} \
    --i-reads {input.query_reads} \
    --p-n-jobs {threads} \
    --o-classification {output.predictions}
```

**Table S17. Commands and parameters to run Q2_VS.**

```
qiime feature-classifier classify-consensus-vsearch \
    --i-query {input.query_reads} \
    --p-threads {threads} \
    --i-reference-reads {input.reference_reads} \
    --i-reference-taxonomy {input.reference_taxonomy} \
    --o-classification {output.predictions} \
    --o-search-results {output.search_results}
```

**Table S18. Commands and parameters to run RDP.**

```
python scripts/fasta_utax2rdp.py \
    {input.train} \
    {output.reference_taxonomy} \
    > {output.reference_reads}

workdir=`echo /usr/bin/rdp_classifier_2.13/dist/classifier.jar | sed "-es/e:/\/e/"`

rd=`dirname $workdir | sed "-es/\/dist//"`

props=$(find $rd | grep rRNAClassifier.properties | head -1)

cp $props {output.properties}

java \
    -Xmx8g \
    -cp /usr/bin/rdp_classifier_2.13/dist/classifier.jar \
    edu/msu/cme/rdp/classifier/train/ClassifierTraineeMaker \
    train \
    -t {input.reference_taxonomy} \
    -s {input.reference_reads} \
    -o {params.tmpdir}
java \
    -Xmx1g \
    -jar /usr/bin/rdp_classifier_2.13/dist/classifier.jar \
    -t {output.properties} \
    -q {input.test} \
    -o {output.predictions}

python scripts/rdpc2tab3.py \
    {input.predictions} \
    50 \
    > {output.predictions}

python scripts/rdpc2tab3.py \
    {input.predictions} \
    80 \
    > {output.predictions}
```

**Table S19. Commands and parameters to run SINTAX.**

```
usearch \
    -sintax \
    {input.test} \
    -db {input.train} \
    -strand plus \
    -tabbedout {output.predictions}

python scripts/bbc_cutoff.py \
    {input.predictions} \
    0.5 \
    > {output.predictions}

python scripts/bbc_cutoff.py \
    {input.predictions} \
    0.8 \
    > {output.predictions}
```

**Table S20. Commands and parameters to run SPINGO.**

```
python scripts/fasta_utax2spingo.py \
    {input.train} \
    > {output.reference_reads}

spingo \
    -i {input.test} \
    -d {input.reference_reads} \
    -p {threads} \
    > {output.predictions}

python scripts/spingo2tab.py \
    {input.predictions} \
    > {output.predictions}
```

**Table S21. Commands and parameters to run TOP.**

```
usearch \
    -db {input.train} \
    -cons_tax {input.test} \
    -strand plus \
    -tabbedout {output.predictions} \
    -strand plus \
    -id 0.7 \
    -maxaccepts 3 \
    -maxrejects 16 \
    -top_hit_only
```

**Table S22. Resources benchmark computed for the dataset SP RDP ITS 90.**

| Method | Training Time (hh:mm:ss) | Training Memory (MB) | Classification Time (hh:mm:ss) | Classification Memory (MB) |
|---|---|---|---|---|
| BTOP | 00:00:00 | 28.28 | 00:00:17 | 21.16 |
| BLCA | 00:00:00 | 22.51 | 00:22:22 | 27.44 |
| RDP | 00:00:20 | 18.15 | 00:00:17 | 26.23 |
| Q2_SK | 00:00:37 | 26.05 | 00:00:39 | 25.40 |
| HiTaC | 00:03:18 | 22.84 | 00:00:03 | 18.24 |
| HiTaC_Filter | 00:51:39 | 22.94 | 00:00:05 | 23.63 |
| Metaxa2 | 01:40:34 | 17.42 | 00:01:18 | 21.11 |
| TOP | - | - | 00:00:00 | 20.84 |
| CT1 | - | - | 00:00:00 | 21.75 |
| KTOP | - | - | 00:00:00 | 21.83 |
| CT2 | - | - | 00:00:00 | 21.86 |
| KNN | - | - | 00:00:01 | 24.03 |
| SINTAX | - | - | 00:00:01 | 24.64 |
| Q1 | - | - | 00:00:03 | 21.52 |
| SPINGO | - | - | 00:00:04 | 24.29 |
| Microclass | - | - | 00:00:14 | 26.50 |
| NBC | - | - | 00:00:15 | 14.87 |
| Q2_BLAST | - | - | 00:02:21 | 20.94 |
| Q2_VS | - | - | 00:02:23 | 25.89 |

**Table S23. Resources benchmark computed for the dataset SP RDP ITS 95.**

| Method | Training Time (hh:mm:ss) | Training Memory (MB) | Classification Time (hh:mm:ss) | Classification Memory (MB) |
|---|---|---|---|---|
| BTOP | 00:00:00 | 17.47 | 00:00:14 | 23.90 |
| BLCA | 00:00:00 | 15.71 | 00:14:10 | 19.42 |
| RDP | 00:00:30 | 20.00 | 00:00:14 | 24.22 |
| Q2_SK | 00:00:49 | 24.25 | 00:00:43 | 25.95 |
| HiTaC | 00:04:00 | 20.32 | 00:00:03 | 19.25 |
| HiTaC_Filter | 01:18:19 | 21.29 | 00:00:04 | 25.39 |
| Metaxa2 | 02:44:28 | 19.03 | 00:01:03 | 18.50 |
| TOP | - | - | 00:00:00 | 16.29 |
| CT2 | - | - | 00:00:00 | 18.63 |
| CT1 | - | - | 00:00:00 | 19.71 |
| KTOP | - | - | 00:00:00 | 20.18 |
| SINTAX | - | - | 00:00:01 | 15.03 |
| KNN | - | - | 00:00:01 | 17.00 |
| Q1 | - | - | 00:00:03 | 27.53 |
| SPINGO | - | - | 00:00:05 | 24.41 |
| NBC | - | - | 00:00:16 | 21.65 |
| Microclass | - | - | 00:00:17 | 23.75 |
| Q2_VS | - | - | 00:01:42 | 27.21 |
| Q2_BLAST | - | - | 00:01:53 | 23.95 |

**Table S24. Resources benchmark computed for the dataset SP RDP ITS 97.**

| Method | Training Time (hh:mm:ss) | Training Memory (MB) | Classification Time (hh:mm:ss) | Classification Memory (MB) |
|---|---|---|---|---|
| BTOP | 00:00:00 | 23.84 | 00:00:17 | 21.43 |
| BLCA | 00:00:00 | 13.42 | 00:14:08 | 19.36 |
| RDP | 00:00:35 | 23.73 | 00:00:17 | 24.48 |
| Q2_SK | 00:00:55 | 22.33 | 00:00:48 | 27.01 |
| HiTaC | 00:04:28 | 14.47 | 00:00:03 | 20.16 |
| HiTaC_Filter | 01:35:15 | 19.27 | 00:00:05 | 22.26 |
| Metaxa2 | 02:42:24 | 12.98 | 00:01:17 | 14.35 |
| CT2 | - | - | 00:00:00 | 20.02 |
| CT1 | - | - | 00:00:00 | 22.20 |
| TOP | - | - | 00:00:00 | 22.48 |
| KTOP | - | - | 00:00:00 | 28.46 |
| SINTAX | - | - | 00:00:01 | 22.36 |
| KNN | - | - | 00:00:01 | 24.37 |
| Q1 | - | - | 00:00:03 | 17.74 |
| SPINGO | - | - | 00:00:06 | 28.26 |
| NBC | - | - | 00:00:18 | 19.19 |
| Microclass | - | - | 00:00:19 | 20.40 |
| Q2_VS | - | - | 00:01:52 | 15.38 |
| Q2_BLAST | - | - | 00:02:18 | 23.07 |

**Table S25. Resources benchmark computed for the dataset SP RDP ITS 99.**

| Method | Training Time (hh:mm:ss) | Training Memory (MB) | Classification Time (hh:mm:ss) | Classification Memory (MB) |
|---|---|---|---|---|
| BTOP | 00:00:00 | 15.54 | 00:00:42 | 27.00 |
| BLCA | 00:00:00 | 25.02 | 00:27:30 | 22.30 |
| RDP | 00:00:43 | 22.35 | 00:00:51 | 25.66 |
| Q2_SK | 00:00:55 | 23.56 | 00:01:11 | 24.18 |
| HiTaC | 00:04:51 | 21.34 | 00:00:07 | 21.29 |
| HiTaC_Filter | 01:32:13 | 18.90 | 00:00:11 | 25.14 |
| Metaxa2 | 01:57:22 | 20.55 | 00:03:08 | 19.63 |
| CT2 | - | - | 00:00:01 | 19.60 |
| CT1 | - | - | 00:00:01 | 20.47 |
| TOP | - | - | 00:00:01 | 20.86 |
| KTOP | - | - | 00:00:01 | 22.03 |
| KNN | - | - | 00:00:02 | 22.84 |
| SINTAX | - | - | 00:00:03 | 23.03 |
| Q1 | - | - | 00:00:05 | 14.33 |
| SPINGO | - | - | 00:00:06 | 19.82 |
| Microclass | - | - | 00:00:23 | 21.20 |
| NBC | - | - | 00:00:33 | 22.34 |
| Q2_BLAST | - | - | 00:05:23 | 24.81 |
| Q2_VS | - | - | 00:08:10 | 29.31 |

**Table S26. Resources benchmark computed for the dataset SP RDP ITS 100.**

| Method | Training Time (hh:mm:ss) | Training Memory (MB) | Classification Time (hh:mm:ss) | Classification Memory (MB) |
|---|---|---|---|---|
| BTOP | 00:00:00 | 27.71 | 00:04:01 | 20.39 |
| BLCA | 00:00:00 | 19.81 | 00:04:30 | 21.77 |
| RDP | 00:00:50 | 24.66 | 00:03:42 | 20.38 |
| Q2_SK | 00:01:17 | 28.97 | 00:02:52 | 27.91 |
| HiTaC | 00:05:16 | 26.81 | 00:00:26 | 25.20 |
| HiTaC_Filter | 02:32:27 | 20.39 | 00:00:45 | 20.36 |
| Metaxa2 | 02:43:29 | 12.67 | 00:18:10 | 13.79 |
| KTOP | - | - | 00:00:03 | 18.77 |
| CT1 | - | - | 00:00:03 | 22.85 |
| TOP | - | - | 00:00:03 | 25.27 |
| CT2 | - | - | 00:00:04 | 18.37 |
| KNN | - | - | 00:00:05 | 23.22 |
| Q1 | - | - | 00:00:11 | 19.79 |
| SPINGO | - | - | 00:00:16 | 24.14 |
| SINTAX | - | - | 00:00:17 | 20.38 |
| Microclass | - | - | 00:00:38 | 17.61 |
| Q2_BLAST | - | - | 00:30:37 | 21.40 |
| Q2_VS | - | - | 00:41:07 | 25.41 |
| NBC | - | Exceeded | - | Exceeded |

**Table S27. Hierarchical metrics computed for the dataset SP RDP ITS 90.**

| Method | F1-score | Precision | Recall |
|---|---|---|---|
| HiTaC_Filter | 86.61 | 96.05 | 78.86 |
| Q2_VS | 84.92 | 93.42 | 77.84 |
| CT2 | 84.92 | 94.10 | 77.37 |
| SINTAX50 | 84.80 | 92.59 | 78.22 |
| RDP80 | 84.80 | 93.83 | 77.36 |
| NBC80 | 84.79 | 93.77 | 77.37 |
| SINTAX80 | 84.76 | 96.63 | 75.48 |
| Q2_BLAST | 84.71 | 93.57 | 77.38 |
| KNN | 83.08 | 98.47 | 71.84 |
| NBC50 | 82.92 | 87.43 | 78.85 |
| Q2_SK | 82.90 | 87.68 | 78.62 |
| RDP50 | 82.79 | 87.16 | 78.83 |
| Metaxa2 | 82.46 | 87.81 | 77.73 |
| CT1 | 80.76 | 82.73 | 78.89 |
| HiTaC | 80.38 | 80.38 | 80.38 |
| BTOP | 79.33 | 79.33 | 79.33 |
| Microclass | 79.24 | 79.24 | 79.24 |
| TOP | 79.16 | 79.16 | 79.16 |
| KTOP | 79.05 | 79.05 | 79.05 |
| BLCA | 78.52 | 84.95 | 72.99 |
| Q1 | 77.80 | 82.43 | 73.66 |
| SPINGO | 32.49 | 75.92 | 20.66 |

**Table S28. Hierarchical metrics computed for the dataset SP RDP ITS 95.**

| Method | F1-score | Precision | Recall |
|---|---|---|---|
| CT2 | 88.13 | 96.94 | 80.79 |
| Q2_VS | 88.08 | 96.45 | 81.04 |
| HiTaC_Filter | 88.05 | 94.66 | 82.30 |
| Q2_BLAST | 87.96 | 96.28 | 80.97 |
| SINTAX80 | 87.40 | 95.32 | 80.70 |
| RDP80 | 86.58 | 92.09 | 81.69 |
| NBC80 | 86.54 | 92.20 | 81.54 |
| SINTAX50 | 86.26 | 90.72 | 82.22 |
| Metaxa2 | 85.67 | 90.34 | 81.46 |
| KNN | 85.41 | 99.25 | 74.95 |
| Q2_SK | 85.09 | 88.16 | 82.22 |
| RDP50 | 84.56 | 86.77 | 82.45 |
| NBC50 | 84.51 | 86.71 | 82.42 |
| Q1 | 84.09 | 86.03 | 82.24 |
| CT1 | 83.88 | 85.67 | 82.17 |
| HiTaC | 82.96 | 82.96 | 82.96 |
| TOP | 82.64 | 82.64 | 82.64 |
| Microclass | 82.59 | 82.59 | 82.59 |
| BTOP | 82.55 | 82.55 | 82.55 |
| BLCA | 82.54 | 86.90 | 78.60 |
| KTOP | 82.50 | 82.50 | 82.50 |
| SPINGO | 37.47 | 76.18 | 24.84 |

**Table S29. Hierarchical metrics computed for the dataset SP RDP ITS 97.**

| Method | F1-score | Precision | Recall |
|---|---|---|---|
| Q2_BLAST | 88.96 | 96.94 | 82.19 |
| Q2_VS | 88.95 | 96.99 | 82.14 |
| SINTAX80 | 88.84 | 95.36 | 83.16 |
| CT2 | 88.75 | 97.15 | 81.68 |
| HiTaC_Filter | 88.65 | 93.71 | 84.10 |
| RDP80 | 87.93 | 92.28 | 83.96 |
| NBC80 | 87.89 | 92.19 | 83.97 |
| SINTAX50 | 87.45 | 90.79 | 84.34 |
| Metaxa2 | 87.18 | 91.63 | 83.14 |
| Q2_SK | 86.80 | 89.31 | 84.43 |
| RDP50 | 86.05 | 87.52 | 84.63 |
| NBC50 | 85.96 | 87.33 | 84.64 |
| KNN | 85.69 | 99.32 | 75.35 |
| CT1 | 85.55 | 87.44 | 83.74 |
| Q1 | 85.49 | 87.31 | 83.75 |
| HiTaC | 84.78 | 84.78 | 84.78 |
| TOP | 84.75 | 84.75 | 84.75 |
| Microclass | 84.73 | 84.73 | 84.73 |
| KTOP | 84.72 | 84.72 | 84.72 |
| BTOP | 84.72 | 84.72 | 84.72 |
| BLCA | 83.72 | 88.46 | 79.47 |
| SPINGO | 40.14 | 79.08 | 26.89 |

**Table S30. Hierarchical metrics computed for the dataset SP RDP ITS 99.**

| Method | F1-score | Precision | Recall |
|---|---|---|---|
| HiTaC_Filter | 96.41 | 98.30 | 94.59 |
| SINTAX50 | 96.28 | 97.25 | 95.33 |
| SINTAX80 | 96.14 | 98.77 | 93.65 |
| HiTaC | 96.06 | 96.06 | 96.06 |
| RDP80 | 96.03 | 97.45 | 94.65 |
| Q2_SK | 96.02 | 96.92 | 95.14 |
| NBC80 | 96.00 | 97.37 | 94.67 |
| BTOP | 95.99 | 95.99 | 95.99 |
| TOP | 95.96 | 95.96 | 95.96 |
| Microclass | 95.96 | 95.96 | 95.96 |
| KTOP | 95.84 | 95.84 | 95.84 |
| RDP50 | 95.66 | 95.97 | 95.35 |
| NBC50 | 95.66 | 95.92 | 95.39 |
| BLCA | 93.33 | 96.37 | 90.47 |
| Q1 | 92.36 | 94.84 | 90.01 |
| Metaxa2 | 92.13 | 98.32 | 86.67 |
| CT1 | 90.60 | 93.29 | 88.06 |
| Q2_VS | 90.02 | 97.90 | 83.32 |
| Q2_BLAST | 89.69 | 97.78 | 82.84 |
| CT2 | 89.02 | 97.77 | 81.71 |
| KNN | 85.20 | 99.99 | 74.23 |
| SPINGO | 54.08 | 95.03 | 37.80 |

**Table S31. Hierarchical metrics computed for the dataset SP RDP ITS 100.**

| Method | F1-score | Precision | Recall |
|---|---|---|---|
| HiTaC | 100.00 | 100.00 | 100.00 |
| BTOP | 100.00 | 100.00 | 100.00 |
| TOP | 99.94 | 99.94 | 99.94 |
| KTOP | 99.92 | 99.92 | 99.92 |
| Microclass | 99.91 | 99.91 | 99.91 |
| HiTaC_Filter | 99.84 | 100.00 | 99.68 |
| SINTAX50 | 99.79 | 99.94 | 99.64 |
| Q2_SK | 99.60 | 99.88 | 99.33 |
| RDP50 | 99.52 | 99.58 | 99.45 |
| RDP80 | 99.38 | 99.78 | 98.99 |
| SINTAX80 | 98.98 | 100.00 | 97.99 |
| Metaxa2 | 98.14 | 100.00 | 96.35 |
| Q1 | 98.10 | 98.72 | 97.50 |
| CT1 | 97.77 | 98.41 | 97.14 |
| Q2_VS | 90.77 | 98.27 | 84.33 |
| Q2_BLAST | 90.51 | 98.17 | 83.96 |
| CT2 | 90.34 | 98.18 | 83.65 |
| KNN | 86.62 | 100.00 | 76.40 |
| SPINGO | 59.54 | 100.00 | 42.39 |

**Table S32. TAXXI metrics computed for the dataset SP RDP ITS 90 at the phylum level.**

| Method | Accuracy | Misclassification Rate | Over-classification Rate | True Positive Rate | Under-classification Rate |
|---|---|---|---|---|---|
| BTOP | 100.00 | 0.00 | . | 100.00 | 0.00 |
| CT1 | 100.00 | 0.00 | . | 100.00 | 0.00 |
| CT2 | 100.00 | 0.00 | . | 100.00 | 0.00 |
| HiTaC | 100.00 | 0.00 | . | 100.00 | 0.00 |
| HiTaC_Filter | 100.00 | 0.00 | . | 100.00 | 0.00 |
| KTOP | 100.00 | 0.00 | . | 100.00 | 0.00 |
| Metaxa2 | 100.00 | 0.00 | . | 100.00 | 0.00 |
| Microclass | 100.00 | 0.00 | . | 100.00 | 0.00 |
| NBC50 | 100.00 | 0.00 | . | 100.00 | 0.00 |
| NBC80 | 100.00 | 0.00 | . | 100.00 | 0.00 |
| Q2_SK | 100.00 | 0.00 | . | 100.00 | 0.00 |
| RDP50 | 100.00 | 0.00 | . | 100.00 | 0.00 |
| RDP80 | 100.00 | 0.00 | . | 100.00 | 0.00 |
| SINTAX50 | 100.00 | 0.00 | . | 100.00 | 0.00 |
| SINTAX80 | 100.00 | 0.00 | . | 100.00 | 0.00 |
| TOP | 100.00 | 0.00 | . | 100.00 | 0.00 |
| KNN | 99.90 | 0.00 | . | 99.90 | 0.10 |
| Q2_VS | 99.80 | 0.00 | . | 99.80 | 0.20 |
| Q2_BLAST | 99.70 | 0.00 | . | 99.70 | 0.30 |
| Q1 | 92.80 | 0.00 | . | 92.80 | 7.20 |
| BLCA | 92.50 | 0.00 | . | 92.50 | 7.50 |
| SPINGO | 0.00 | 0.00 | . | 0.00 | 100.00 |

**Table S33. TAXXI metrics computed for the dataset SP RDP ITS 90 at the class level.**

| Method | Accuracy | Misclassification Rate | Over-classification Rate | True Positive Rate | Under-classification Rate |
|---|---|---|---|---|---|
| CT2 | 99.90 | 0.10 | . | 99.90 | 0.00 |
| HiTaC | 99.90 | 0.10 | . | 99.90 | 0.00 |
| HiTaC_Filter | 99.90 | 0.10 | . | 99.90 | 0.00 |
| Q2_VS | 99.80 | 0.10 | . | 99.80 | 0.20 |
| BTOP | 99.80 | 0.20 | . | 99.80 | 0.00 |
| Microclass | 99.80 | 0.20 | . | 99.80 | 0.00 |
| Q2_SK | 99.80 | 0.20 | . | 99.80 | 0.00 |
| Q2_BLAST | 99.70 | 0.10 | . | 99.70 | 0.30 |
| Metaxa2 | 99.70 | 0.30 | . | 99.70 | 0.00 |
| TOP | 99.70 | 0.30 | . | 99.70 | 0.00 |
| RDP50 | 99.60 | 0.40 | . | 99.60 | 0.00 |
| NBC80 | 99.50 | 0.10 | . | 99.50 | 0.40 |
| RDP80 | 99.50 | 0.10 | . | 99.50 | 0.40 |
| SINTAX80 | 99.50 | 0.10 | . | 99.50 | 0.40 |
| SINTAX50 | 99.50 | 0.20 | . | 99.50 | 0.30 |
| CT1 | 99.50 | 0.40 | . | 99.50 | 0.10 |
| NBC50 | 99.50 | 0.40 | . | 99.50 | 0.10 |
| KTOP | 99.50 | 0.50 | . | 99.50 | 0.00 |
| KNN | 98.60 | 0.10 | . | 98.60 | 1.30 |
| Q1 | 92.60 | 0.20 | . | 92.60 | 7.20 |
| BLCA | 92.40 | 0.20 | . | 92.40 | 7.50 |
| SPINGO | 0.00 | 0.00 | . | 0.00 | 100.00 |

**Table S34. TAXXI metrics computed for the dataset SP RDP ITS 90 at the order level.**

| Method | Accuracy | Misclassification Rate | Over-classification Rate | True Positive Rate | Under-classification Rate |
|---|---|---|---|---|---|
| HiTaC_Filter | 98.20 | 1.60 | . | 98.20 | 0.20 |
| HiTaC | 98.20 | 1.80 | . | 98.20 | 0.00 |
| Microclass | 98.20 | 1.80 | . | 98.20 | 0.00 |
| Q2_VS | 98.10 | 1.50 | . | 98.10 | 0.40 |
| Q2_SK | 98.10 | 1.60 | . | 98.10 | 0.30 |
| BTOP | 98.10 | 1.90 | . | 98.10 | 0.00 |
| TOP | 98.10 | 1.90 | . | 98.10 | 0.00 |
| Q2_BLAST | 98.00 | 1.60 | . | 98.00 | 0.50 |
| CT2 | 97.90 | 1.80 | . | 97.90 | 0.40 |
| NBC50 | 97.90 | 1.90 | . | 97.90 | 0.30 |
| RDP50 | 97.90 | 2.00 | . | 97.90 | 0.10 |
| KTOP | 97.90 | 2.10 | . | 97.90 | 0.00 |
| CT1 | 97.90 | 2.10 | . | 97.90 | 0.10 |
| RDP80 | 97.80 | 1.30 | . | 97.80 | 0.80 |
| NBC80 | 97.80 | 1.30 | . | 97.80 | 0.90 |
| SINTAX50 | 97.80 | 1.50 | . | 97.80 | 0.70 |
| Metaxa2 | 97.20 | 2.40 | . | 97.20 | 0.40 |
| SINTAX80 | 97.10 | 1.30 | . | 97.10 | 1.60 |
| KNN | 93.20 | 0.90 | . | 93.20 | 5.80 |
| Q1 | 91.40 | 1.40 | . | 91.40 | 7.20 |
| BLCA | 90.40 | 1.80 | . | 90.40 | 7.80 |
| SPINGO | 0.00 | 0.00 | . | 0.00 | 100.00 |

**Table S35. TAXXI metrics computed for the dataset SP RDP ITS 90 at the family level.**

| Method | Accuracy | Misclassification Rate | Over-classification Rate | True Positive Rate | Under-classification Rate |
|---|---|---|---|---|---|
| HiTaC | 95.10 | 4.90 | . | 95.10 | 0.00 |
| HiTaC_Filter | 94.00 | 3.40 | . | 94.00 | 2.60 |
| TOP | 93.90 | 6.00 | . | 94.00 | 0.00 |
| Microclass | 93.90 | 6.10 | . | 93.90 | 0.00 |
| BTOP | 93.70 | 6.20 | . | 93.80 | 0.00 |
| KTOP | 93.40 | 6.50 | . | 93.50 | 0.00 |
| CT1 | 93.30 | 6.60 | . | 93.40 | 0.10 |
| NBC50 | 93.20 | 5.40 | . | 93.20 | 1.40 |
| RDP50 | 93.20 | 5.90 | . | 93.30 | 0.80 |
| Q2_SK | 93.10 | 4.60 | . | 93.20 | 2.30 |
| SINTAX50 | 92.70 | 4.30 | . | 92.70 | 3.00 |
| Q2_VS | 91.70 | 5.40 | . | 91.80 | 2.80 |
| NBC80 | 91.60 | 3.30 | . | 91.70 | 5.00 |
| RDP80 | 91.50 | 3.50 | . | 91.60 | 5.00 |
| Q2_BLAST | 91.40 | 5.60 | . | 91.40 | 2.90 |
| SPINGO | 91.30 | 3.60 | . | 91.30 | 5.10 |
| CT2 | 91.30 | 5.70 | . | 91.30 | 3.00 |
| Metaxa2 | 90.50 | 6.40 | . | 90.50 | 3.10 |
| SINTAX80 | 87.60 | 2.30 | . | 87.70 | 10.00 |
| Q1 | 87.30 | 5.20 | . | 87.30 | 7.40 |
| BLCA | 86.90 | 4.40 | . | 87.00 | 8.60 |
| KNN | 80.20 | 1.40 | . | 80.20 | 18.40 |

**Table S36. TAXXI metrics computed for the dataset SP RDP ITS 90 at the genus level.**

| Method | Accuracy | Misclassification Rate | Over-classification Rate | True Positive Rate | Under-classification Rate |
|---|---|---|---|---|---|
| HiTaC | 69.50 | 19.70 | 100.00 | 80.30 | 0.00 |
| HiTaC_Filter | 64.80 | 10.30 | 44.10 | 69.30 | 20.40 |
| NBC50 | 64.20 | 17.60 | 67.00 | 70.90 | 11.50 |
| RDP50 | 63.90 | 18.60 | 67.80 | 70.60 | 10.80 |
| BTOP | 63.70 | 26.60 | 100.00 | 73.40 | 0.00 |
| Microclass | 62.70 | 27.50 | 100.00 | 72.50 | 0.00 |
| Q2_SK | 62.40 | 18.80 | 62.50 | 68.50 | 12.70 |
| KTOP | 62.40 | 27.80 | 100.00 | 72.20 | 0.00 |
| TOP | 62.40 | 27.90 | 100.00 | 72.10 | 0.00 |
| CT1 | 62.10 | 27.90 | 92.70 | 71.00 | 1.10 |
| SINTAX50 | 61.90 | 12.20 | 47.10 | 66.50 | 21.30 |
| Q1 | 60.30 | 23.00 | 81.20 | 68.00 | 9.00 |
| Metaxa2 | 59.00 | 19.70 | 71.60 | 65.60 | 14.70 |
| Q2_VS | 58.50 | 19.50 | 61.70 | 64.20 | 16.40 |
| BLCA | 58.40 | 15.90 | 72.40 | 65.00 | 19.10 |
| SPINGO | 57.90 | 10.40 | 41.80 | 61.70 | 27.90 |
| RDP80 | 57.70 | 9.00 | 34.90 | 60.80 | 30.20 |
| NBC80 | 57.60 | 8.80 | 36.80 | 60.90 | 30.30 |
| Q2_BLAST | 56.00 | 19.40 | 61.70 | 61.40 | 19.20 |
| CT2 | 55.20 | 19.00 | 64.00 | 60.80 | 20.30 |
| SINTAX80 | 49.20 | 5.90 | 22.20 | 51.00 | 43.10 |
| KNN | 34.80 | 3.30 | 18.40 | 35.80 | 60.90 |

**Table S37. TAXXI metrics computed for the dataset SP RDP ITS 90 at the species level.**

| Method | Accuracy | Misclassification Rate | Over-classification Rate | True Positive Rate | Under-classification Rate |
|---|---|---|---|---|---|
| CT2 | 0.00 | . | 1.30 | . | . |
| HiTaC_Filter | 0.00 | . | 2.70 | . | . |
| Q2_BLAST | 0.00 | . | 4.80 | . | . |
| Q2_VS | 0.00 | . | 6.20 | . | . |
| SINTAX80 | 0.00 | . | 6.60 | . | . |
| RDP80 | 0.00 | . | 18.20 | . | . |
| NBC80 | 0.00 | . | 18.60 | . | . |
| SINTAX50 | 0.00 | . | 20.80 | . | . |
| SPINGO | 0.00 | . | 22.50 | . | . |
| Metaxa2 | 0.00 | . | 39.70 | . | . |
| Q2_SK | 0.00 | . | 46.20 | . | . |
| NBC50 | 0.00 | . | 47.50 | . | . |
| RDP50 | 0.00 | . | 47.70 | . | . |
| BLCA | 0.00 | . | 53.10 | . | . |
| Q1 | 0.00 | . | 64.90 | . | . |
| CT1 | 0.00 | . | 69.50 | . | . |
| BTOP | 0.00 | . | 100.00 | . | . |
| HiTaC | 0.00 | . | 100.00 | . | . |
| KTOP | 0.00 | . | 100.00 | . | . |
| Microclass | 0.00 | . | 100.00 | . | . |
| TOP | 0.00 | . | 100.00 | . | . |
| KNN | . | . | 0.00 | . | . |

**Table S38. TAXXI metrics computed for the dataset SP RDP ITS 95 at the phylum level.**

| Method | Accuracy | Misclassification Rate | Over-classification Rate | True Positive Rate | Under-classification Rate |
|---|---|---|---|---|---|
| BTOP | 100.00 | 0.00 | . | 100.00 | 0.00 |
| CT1 | 100.00 | 0.00 | . | 100.00 | 0.00 |
| CT2 | 100.00 | 0.00 | . | 100.00 | 0.00 |
| HiTaC | 100.00 | 0.00 | . | 100.00 | 0.00 |
| HiTaC_Filter | 100.00 | 0.00 | . | 100.00 | 0.00 |
| KNN | 100.00 | 0.00 | . | 100.00 | 0.00 |
| KTOP | 100.00 | 0.00 | . | 100.00 | 0.00 |
| Metaxa2 | 100.00 | 0.00 | . | 100.00 | 0.00 |
| Microclass | 100.00 | 0.00 | . | 100.00 | 0.00 |
| NBC50 | 100.00 | 0.00 | . | 100.00 | 0.00 |
| NBC80 | 100.00 | 0.00 | . | 100.00 | 0.00 |
| Q2_BLAST | 100.00 | 0.00 | . | 100.00 | 0.00 |
| Q2_SK | 100.00 | 0.00 | . | 100.00 | 0.00 |
| Q2_VS | 100.00 | 0.00 | . | 100.00 | 0.00 |
| RDP50 | 100.00 | 0.00 | . | 100.00 | 0.00 |
| RDP80 | 100.00 | 0.00 | . | 100.00 | 0.00 |
| SINTAX50 | 100.00 | 0.00 | . | 100.00 | 0.00 |
| SINTAX80 | 100.00 | 0.00 | . | 100.00 | 0.00 |
| TOP | 100.00 | 0.00 | . | 100.00 | 0.00 |
| Q1 | 99.80 | 0.00 | . | 99.80 | 0.20 |
| BLCA | 95.90 | 0.00 | . | 95.90 | 4.10 |
| SPINGO | 0.00 | 0.00 | . | 0.00 | 100.00 |

Table S39. **TAXXI metrics computed for the dataset SP RDP ITS 95 at the class level.**

| Method | Accuracy | Misclassification Rate | Over-classification Rate | True Positive Rate | Under-classification Rate |
|---|---|---|---|---|---|
| CT2 | 99.70 | 0.30 | . | 99.70 | 0.00 |
| HiTaC | 99.70 | 0.30 | . | 99.70 | 0.00 |
| HiTaC_Filter | 99.70 | 0.30 | . | 99.70 | 0.00 |
| Q2_BLAST | 99.70 | 0.30 | . | 99.70 | 0.00 |
| Q2_VS | 99.70 | 0.30 | . | 99.70 | 0.00 |
| CT1 | 99.60 | 0.40 | . | 99.60 | 0.00 |
| KTOP | 99.60 | 0.40 | . | 99.60 | 0.00 |
| TOP | 99.60 | 0.40 | . | 99.60 | 0.00 |
| SINTAX50 | 99.50 | 0.40 | . | 99.50 | 0.10 |
| BTOP | 99.50 | 0.50 | . | 99.50 | 0.00 |
| RDP50 | 99.50 | 0.50 | . | 99.50 | 0.00 |
| NBC80 | 99.40 | 0.40 | . | 99.40 | 0.20 |
| Q2_SK | 99.40 | 0.40 | . | 99.40 | 0.20 |
| RDP80 | 99.40 | 0.40 | . | 99.40 | 0.20 |
| SINTAX80 | 99.40 | 0.40 | . | 99.40 | 0.20 |
| NBC50 | 99.40 | 0.50 | . | 99.40 | 0.10 |
| Metaxa2 | 99.40 | 0.50 | . | 99.40 | 0.20 |
| Q1 | 99.40 | 0.50 | . | 99.40 | 0.20 |
| Microclass | 99.40 | 0.60 | . | 99.40 | 0.00 |
| KNN | 98.70 | 0.30 | . | 98.70 | 0.90 |
| BLCA | 95.40 | 0.50 | . | 95.40 | 4.10 |
| SPINGO | 0.00 | 0.00 | . | 0.00 | 100.00 |

**Table S40. TAXXI metrics computed for the dataset SP RDP ITS 95 at the order level.**

| Method | Accuracy | Misclassification Rate | Over-classification Rate | True Positive Rate | Under-classification Rate |
|---|---|---|---|---|---|
| Q2_VS | 99.10 | 0.60 | . | 99.10 | 0.20 |
| CT2 | 99.10 | 0.70 | . | 99.10 | 0.20 |
| Q2_BLAST | 99.10 | 0.70 | . | 99.10 | 0.20 |
| HiTaC_Filter | 99.10 | 0.80 | . | 99.10 | 0.10 |
| HiTaC | 99.10 | 0.90 | . | 99.10 | 0.00 |
| KTOP | 99.10 | 0.90 | . | 99.10 | 0.00 |
| TOP | 99.10 | 0.90 | . | 99.10 | 0.00 |
| SINTAX50 | 99.00 | 0.80 | . | 99.00 | 0.20 |
| RDP50 | 99.00 | 1.00 | . | 99.00 | 0.00 |
| RDP80 | 98.90 | 0.90 | . | 98.90 | 0.20 |
| NBC50 | 98.90 | 1.00 | . | 98.90 | 0.10 |
| BTOP | 98.90 | 1.10 | . | 98.90 | 0.00 |
| CT1 | 98.90 | 1.10 | . | 98.90 | 0.00 |
| Microclass | 98.90 | 1.10 | . | 98.90 | 0.00 |
| Q2_SK | 98.80 | 0.90 | . | 98.80 | 0.20 |
| NBC80 | 98.70 | 0.80 | . | 98.70 | 0.60 |
| Metaxa2 | 98.70 | 1.00 | . | 98.70 | 0.20 |
| Q1 | 98.70 | 1.20 | . | 98.70 | 0.20 |
| SINTAX80 | 98.40 | 0.60 | . | 98.40 | 1.00 |
| KNN | 94.80 | 0.50 | . | 94.80 | 4.70 |
| BLCA | 94.80 | 0.90 | . | 94.80 | 4.30 |
| SPINGO | 0.00 | 0.00 | . | 0.00 | 100.00 |

**Table S41. TAXXI metrics computed for the dataset SP RDP ITS 95 at the family level.**

| Method | Accuracy | Misclassification Rate | Over-classification Rate | True Positive Rate | Under-classification Rate |
|---|---|---|---|---|---|
| HiTaC | 98.00 | 1.90 | . | 98.10 | 0.00 |
| BTOP | 97.80 | 2.10 | . | 97.90 | 0.00 |
| KTOP | 97.80 | 2.10 | . | 97.90 | 0.00 |
| TOP | 97.80 | 2.10 | . | 97.90 | 0.00 |
| HiTaC_Filter | 97.60 | 1.00 | . | 97.60 | 1.30 |
| RDP50 | 97.60 | 2.20 | . | 97.70 | 0.10 |
| Microclass | 97.50 | 2.40 | . | 97.60 | 0.00 |
| SINTAX50 | 97.30 | 1.70 | . | 97.40 | 0.90 |
| NBC50 | 97.30 | 2.20 | . | 97.40 | 0.40 |
| Q2_SK | 97.20 | 1.80 | . | 97.30 | 0.90 |
| Q1 | 97.20 | 2.50 | . | 97.20 | 0.30 |
| CT1 | 97.00 | 2.90 | . | 97.10 | 0.00 |
| RDP80 | 96.80 | 1.40 | . | 96.90 | 1.70 |
| SPINGO | 96.80 | 1.50 | . | 96.90 | 1.60 |
| NBC80 | 96.60 | 1.30 | . | 96.70 | 2.00 |
| Metaxa2 | 96.40 | 2.10 | . | 96.40 | 1.60 |
| SINTAX80 | 96.10 | 1.00 | . | 96.10 | 2.80 |
| Q2_VS | 95.80 | 2.20 | . | 95.80 | 2.00 |
| CT2 | 95.70 | 2.30 | . | 95.70 | 2.00 |
| Q2_BLAST | 95.60 | 2.50 | . | 95.60 | 2.00 |
| BLCA | 93.00 | 1.90 | . | 93.00 | 5.10 |
| KNN | 85.20 | 0.90 | . | 85.20 | 13.80 |

**Table S42. TAXXI metrics computed for the dataset SP RDP ITS 95 at the genus level.**

| Method | Accuracy | Misclassification Rate | Over-classification Rate | True Positive Rate | Under-classification Rate |
|---|---|---|---|---|---|
| HiTaC | 83.60 | 12.90 | 100.00 | 87.10 | 0.00 |
| Microclass | 82.00 | 14.60 | 100.00 | 85.40 | 0.00 |
| TOP | 81.90 | 14.70 | 100.00 | 85.30 | 0.00 |
| NBC50 | 81.80 | 13.10 | 78.40 | 84.50 | 2.40 |
| RDP50 | 81.40 | 12.90 | 82.40 | 84.30 | 2.80 |
| BTOP | 81.30 | 15.20 | 100.00 | 84.80 | 0.00 |
| Q1 | 81.00 | 14.70 | 90.20 | 84.10 | 1.20 |
| KTOP | 80.80 | 15.80 | 100.00 | 84.20 | 0.00 |
| HiTaC_Filter | 80.60 | 8.30 | 70.60 | 82.90 | 8.70 |
| Q2_SK | 80.60 | 11.80 | 80.40 | 83.30 | 4.90 |
| SINTAX50 | 80.50 | 10.60 | 72.50 | 82.90 | 6.40 |
| CT1 | 79.80 | 16.10 | 96.10 | 83.00 | 0.90 |
| SPINGO | 78.10 | 9.10 | 64.70 | 80.20 | 10.60 |
| RDP80 | 77.60 | 8.30 | 66.70 | 79.80 | 11.90 |
| NBC80 | 77.20 | 8.20 | 62.70 | 79.20 | 12.60 |
| Metaxa2 | 76.60 | 10.60 | 74.50 | 79.00 | 10.40 |
| BLCA | 75.90 | 10.20 | 76.50 | 78.30 | 11.40 |
| Q2_VS | 73.30 | 11.00 | 76.50 | 75.70 | 13.30 |
| Q2_BLAST | 73.10 | 12.40 | 76.50 | 75.50 | 12.10 |
| SINTAX80 | 72.40 | 5.40 | 49.00 | 73.90 | 20.80 |
| CT2 | 72.00 | 10.60 | 70.60 | 74.10 | 15.20 |
| KNN | 47.70 | 1.90 | 9.80 | 47.90 | 50.20 |

**Table S43. TAXXI metrics computed for the dataset SP RDP ITS 95 at the species level.**

| Method | Accuracy | Misclassification Rate | Over-classification Rate | True Positive Rate | Under-classification Rate |
|---|---|---|---|---|---|
| Q2_BLAST | 2.00 | . | 3.40 | . | . |
| Q2_VS | 1.80 | . | 4.10 | . | . |
| HiTaC_Filter | 0.80 | . | 19.50 | . | . |
| SINTAX80 | 0.40 | . | 18.60 | . | . |
| SINTAX50 | 0.40 | . | 42.80 | . | . |
| Q2_SK | 0.30 | . | 59.60 | . | . |
| BTOP | 0.30 | . | 100.00 | . | . |
| NBC80 | 0.20 | . | 35.30 | . | . |
| RDP80 | 0.20 | . | 35.60 | . | . |
| SPINGO | 0.20 | . | 39.90 | . | . |
| RDP50 | 0.20 | . | 68.50 | . | . |
| NBC50 | 0.20 | . | 69.00 | . | . |
| HiTaC | 0.20 | . | 100.00 | . | . |
| KTOP | 0.20 | . | 100.00 | . | . |
| Microclass | 0.20 | . | 100.00 | . | . |
| TOP | 0.20 | . | 100.00 | . | . |
| BLCA | 0.10 | . | 62.60 | . | . |
| Q1 | 0.10 | . | 71.40 | . | . |
| CT2 | 0.00 | . | 1.50 | . | . |
| Metaxa2 | 0.00 | . | 44.20 | . | . |
| CT1 | 0.00 | . | 72.50 | . | . |
| KNN | . | . | 0.00 | . | . |

**Table S44. TAXXI metrics computed for the dataset SP RDP ITS 97 at the phylum level.**

| Method | Accuracy | Misclassification Rate | Over-classification Rate | True Positive Rate | Under-classification Rate |
|---|---|---|---|---|---|
| BTOP | 100.00 | 0.00 | . | 100.00 | 0.00 |
| CT1 | 100.00 | 0.00 | . | 100.00 | 0.00 |
| CT2 | 100.00 | 0.00 | . | 100.00 | 0.00 |
| HiTaC | 100.00 | 0.00 | . | 100.00 | 0.00 |
| HiTaC_Filter | 100.00 | 0.00 | . | 100.00 | 0.00 |
| KTOP | 100.00 | 0.00 | . | 100.00 | 0.00 |
| Microclass | 100.00 | 0.00 | . | 100.00 | 0.00 |
| NBC50 | 100.00 | 0.00 | . | 100.00 | 0.00 |
| NBC80 | 100.00 | 0.00 | . | 100.00 | 0.00 |
| Q2_BLAST | 100.00 | 0.00 | . | 100.00 | 0.00 |
| Q2_SK | 100.00 | 0.00 | . | 100.00 | 0.00 |
| RDP50 | 100.00 | 0.00 | . | 100.00 | 0.00 |
| RDP80 | 100.00 | 0.00 | . | 100.00 | 0.00 |
| SINTAX50 | 100.00 | 0.00 | . | 100.00 | 0.00 |
| SINTAX80 | 100.00 | 0.00 | . | 100.00 | 0.00 |
| TOP | 100.00 | 0.00 | . | 100.00 | 0.00 |
| KNN | 99.90 | 0.00 | . | 99.90 | 0.10 |
| Metaxa2 | 99.90 | 0.00 | . | 99.90 | 0.10 |
| Q1 | 99.90 | 0.00 | . | 99.90 | 0.10 |
| Q2_VS | 99.90 | 0.00 | . | 99.90 | 0.10 |
| BLCA | 94.50 | 0.00 | . | 94.50 | 5.50 |
| SPINGO | 0.00 | 0.00 | . | 0.00 | 100.00 |

**Table S45. TAXXI metrics computed for the dataset SP RDP ITS 97 at the class level.**

| Method | Accuracy | Misclassification Rate | Over-classification Rate | True Positive Rate | Under-classification Rate |
|---|---|---|---|---|---|
| BTOP | 99.90 | 0.10 | . | 99.90 | 0.00 |
| CT2 | 99.90 | 0.10 | . | 99.90 | 0.00 |
| HiTaC | 99.90 | 0.10 | . | 99.90 | 0.00 |
| HiTaC_Filter | 99.90 | 0.10 | . | 99.90 | 0.00 |
| KTOP | 99.90 | 0.10 | . | 99.90 | 0.00 |
| Microclass | 99.90 | 0.10 | . | 99.90 | 0.00 |
| NBC50 | 99.90 | 0.10 | . | 99.90 | 0.00 |
| NBC80 | 99.90 | 0.10 | . | 99.90 | 0.00 |
| Q2_BLAST | 99.90 | 0.10 | . | 99.90 | 0.00 |
| Q2_SK | 99.90 | 0.10 | . | 99.90 | 0.00 |
| RDP50 | 99.90 | 0.10 | . | 99.90 | 0.00 |
| RDP80 | 99.90 | 0.10 | . | 99.90 | 0.00 |
| SINTAX50 | 99.90 | 0.10 | . | 99.90 | 0.00 |
| TOP | 99.90 | 0.10 | . | 99.90 | 0.00 |
| CT1 | 99.90 | 0.10 | . | 99.90 | 0.10 |
| Q2_VS | 99.90 | 0.10 | . | 99.90 | 0.10 |
| SINTAX80 | 99.90 | 0.10 | . | 99.90 | 0.10 |
| Q1 | 99.80 | 0.10 | . | 99.80 | 0.10 |
| Metaxa2 | 99.70 | 0.10 | . | 99.70 | 0.20 |
| KNN | 98.10 | 0.10 | . | 98.10 | 1.80 |
| BLCA | 94.40 | 0.10 | . | 94.40 | 5.50 |
| SPINGO | 0.00 | 0.00 | . | 0.00 | 100.00 |

**Table S46. TAXXI metrics computed for the dataset SP RDP ITS 97 at the order level.**

| Method | Accuracy | Misclassification Rate | Over-classification Rate | True Positive Rate | Under-classification Rate |
|---|---|---|---|---|---|
| BTOP | 99.40 | 0.60 | . | 99.40 | 0.00 |
| HiTaC | 99.40 | 0.60 | . | 99.40 | 0.00 |
| KTOP | 99.40 | 0.60 | . | 99.40 | 0.00 |
| Microclass | 99.40 | 0.60 | . | 99.40 | 0.00 |
| NBC50 | 99.40 | 0.60 | . | 99.40 | 0.00 |
| NBC80 | 99.40 | 0.60 | . | 99.40 | 0.00 |
| Q2_SK | 99.40 | 0.60 | . | 99.40 | 0.00 |
| RDP50 | 99.40 | 0.60 | . | 99.40 | 0.00 |
| RDP80 | 99.40 | 0.60 | . | 99.40 | 0.00 |
| TOP | 99.40 | 0.60 | . | 99.40 | 0.00 |
| HiTaC_Filter | 99.40 | 0.60 | . | 99.40 | 0.10 |
| SINTAX50 | 99.40 | 0.60 | . | 99.40 | 0.10 |
| Q2_VS | 99.30 | 0.50 | . | 99.30 | 0.20 |
| SINTAX80 | 99.30 | 0.50 | . | 99.30 | 0.20 |
| Q1 | 99.30 | 0.60 | . | 99.30 | 0.10 |
| Q2_BLAST | 99.30 | 0.60 | . | 99.30 | 0.10 |
| CT2 | 99.20 | 0.60 | . | 99.20 | 0.10 |
| CT1 | 99.20 | 0.70 | . | 99.20 | 0.10 |
| Metaxa2 | 99.00 | 0.50 | . | 99.00 | 0.50 |
| KNN | 95.80 | 0.40 | . | 95.80 | 3.70 |
| BLCA | 93.90 | 0.60 | . | 93.90 | 5.60 |
| SPINGO | 0.00 | 0.00 | . | 0.00 | 100.00 |

**Table S47. TAXXI metrics computed for the dataset SP RDP ITS 97 at the family level.**

| Method | Accuracy | Misclassification Rate | Over-classification Rate | True Positive Rate | Under-classification Rate |
|---|---|---|---|---|---|
| TOP | 99.20 | 0.70 | . | 99.30 | 0.00 |
| BTOP | 99.20 | 0.80 | . | 99.20 | 0.00 |
| KTOP | 99.20 | 0.80 | . | 99.20 | 0.00 |
| RDP80 | 99.10 | 0.70 | . | 99.20 | 0.10 |
| Microclass | 99.10 | 0.80 | . | 99.20 | 0.00 |
| NBC50 | 99.10 | 0.80 | . | 99.20 | 0.00 |
| Q2_SK | 99.10 | 0.80 | . | 99.20 | 0.00 |
| RDP50 | 99.10 | 0.80 | . | 99.20 | 0.00 |
| NBC80 | 99.00 | 0.60 | . | 99.10 | 0.30 |
| SINTAX50 | 98.90 | 0.70 | . | 99.00 | 0.30 |
| HiTaC_Filter | 98.90 | 0.80 | . | 98.90 | 0.30 |
| HiTaC | 98.90 | 1.00 | . | 99.00 | 0.00 |
| SPINGO | 98.70 | 0.70 | . | 98.80 | 0.50 |
| CT1 | 98.70 | 1.10 | . | 98.70 | 0.10 |
| Q1 | 98.50 | 1.30 | . | 98.50 | 0.20 |
| SINTAX80 | 98.40 | 0.70 | . | 98.40 | 0.90 |
| Metaxa2 | 98.20 | 0.70 | . | 98.20 | 1.10 |
| Q2_BLAST | 97.50 | 1.80 | . | 97.50 | 0.80 |
| Q2_VS | 97.50 | 1.90 | . | 97.50 | 0.60 |
| CT2 | 96.50 | 2.30 | . | 96.50 | 1.20 |
| BLCA | 93.40 | 0.80 | . | 93.40 | 5.80 |
| KNN | 85.70 | 0.60 | . | 85.70 | 13.70 |

**Table S48. TAXXI metrics computed for the dataset SP RDP ITS 97 at the genus level.**

| Method | Accuracy | Misclassification Rate | Over-classification Rate | True Positive Rate | Under-classification Rate |
|---|---|---|---|---|---|
| HiTaC | 91.50 | 7.70 | 100.00 | 92.30 | 0.00 |
| Microclass | 91.10 | 8.20 | 100.00 | 91.80 | 0.00 |
| KTOP | 91.00 | 8.30 | 100.00 | 91.70 | 0.00 |
| TOP | 91.00 | 8.30 | 100.00 | 91.70 | 0.00 |
| BTOP | 90.90 | 8.40 | 100.00 | 91.60 | 0.00 |
| NBC50 | 90.60 | 7.40 | 100.00 | 91.30 | 1.30 |
| RDP50 | 90.50 | 7.10 | 100.00 | 91.20 | 1.70 |
| SINTAX50 | 89.10 | 6.70 | 81.80 | 89.60 | 3.60 |
| Q2_SK | 89.10 | 6.70 | 100.00 | 89.80 | 3.50 |
| CT1 | 87.90 | 10.60 | 90.90 | 88.60 | 0.90 |
| HiTaC_Filter | 87.80 | 5.00 | 81.80 | 88.40 | 6.60 |
| Q1 | 87.30 | 11.40 | 81.80 | 87.90 | 0.70 |
| SPINGO | 86.60 | 5.00 | 81.80 | 87.20 | 7.90 |
| RDP80 | 86.50 | 4.70 | 81.80 | 87.00 | 8.30 |
| NBC80 | 86.50 | 4.90 | 81.80 | 87.10 | 8.00 |
| Metaxa2 | 85.00 | 7.30 | 90.90 | 85.60 | 7.10 |
| BLCA | 83.10 | 6.80 | 81.80 | 83.60 | 9.60 |
| SINTAX80 | 82.50 | 3.60 | 54.50 | 82.90 | 13.50 |
| Q2_BLAST | 78.10 | 11.10 | 81.80 | 78.60 | 10.20 |
| Q2_VS | 77.90 | 10.40 | 90.90 | 78.50 | 11.10 |
| CT2 | 76.10 | 11.30 | 81.80 | 76.60 | 12.10 |
| KNN | 48.20 | 1.90 | 54.50 | 48.40 | 49.70 |

**Table S49. TAXXI metrics computed for the dataset SP RDP ITS 97 at the species level.**

| Method | Accuracy | Misclassification Rate | Over-classification Rate | True Positive Rate | Under-classification Rate |
|---|---|---|---|---|---|
| Q2_BLAST | 9.40 | 0.00 | 4.20 | 18.30 | 81.70 |
| SINTAX80 | 8.80 | 1.70 | 24.00 | 56.70 | 41.70 |
| Q2_VS | 8.30 | 1.70 | 4.40 | 16.70 | 81.70 |
| HiTaC_Filter | 7.90 | 1.70 | 33.80 | 68.30 | 30.00 |
| NBC80 | 6.50 | 6.70 | 44.60 | 71.70 | 21.70 |
| SPINGO | 6.40 | 3.30 | 44.70 | 71.70 | 25.00 |
| RDP80 | 6.40 | 8.30 | 44.00 | 70.00 | 21.70 |
| SINTAX50 | 5.80 | 11.70 | 52.90 | 75.00 | 13.30 |
| Q2_SK | 5.30 | 10.00 | 64.00 | 81.70 | 8.30 |
| RDP50 | 4.40 | 13.30 | 77.90 | 81.70 | 5.00 |
| NBC50 | 4.30 | 13.30 | 79.10 | 81.70 | 5.00 |
| BLCA | 4.20 | 5.00 | 60.50 | 61.70 | 33.30 |
| HiTaC | 3.70 | 13.30 | 100.00 | 86.70 | 0.00 |
| TOP | 3.70 | 13.30 | 100.00 | 86.70 | 0.00 |
| BTOP | 3.70 | 14.80 | 100.00 | 85.20 | 0.00 |
| Microclass | 3.60 | 15.00 | 100.00 | 85.00 | 0.00 |
| KTOP | 3.50 | 16.70 | 100.00 | 83.30 | 0.00 |
| CT2 | 3.40 | 0.00 | 2.00 | 5.00 | 95.00 |
| Q1 | 2.30 | 23.30 | 73.40 | 40.00 | 36.70 |
| CT1 | 0.80 | 38.30 | 72.60 | 13.30 | 48.30 |
| Metaxa2 | 0.70 | 0.00 | 45.90 | 8.30 | 91.70 |
| KNN | 0.00 | 0.00 | 0.20 | 0.00 | 100.00 |

**Table S50. TAXXI metrics computed for the dataset SP RDP ITS 99 at the phylum level.**

| Method | Accuracy | Misclassification Rate | Over-classification Rate | True Positive Rate | Under-classification Rate |
|---|---|---|---|---|---|
| BTOP | 100.00 | 0.00 | . | 100.00 | 0.00 |
| CT1 | 100.00 | 0.00 | . | 100.00 | 0.00 |
| CT2 | 100.00 | 0.00 | . | 100.00 | 0.00 |
| HiTaC | 100.00 | 0.00 | . | 100.00 | 0.00 |
| HiTaC_Filter | 100.00 | 0.00 | . | 100.00 | 0.00 |
| KTOP | 100.00 | 0.00 | . | 100.00 | 0.00 |
| Metaxa2 | 100.00 | 0.00 | . | 100.00 | 0.00 |
| Microclass | 100.00 | 0.00 | . | 100.00 | 0.00 |
| NBC50 | 100.00 | 0.00 | . | 100.00 | 0.00 |
| NBC80 | 100.00 | 0.00 | . | 100.00 | 0.00 |
| Q2_SK | 100.00 | 0.00 | . | 100.00 | 0.00 |
| RDP50 | 100.00 | 0.00 | . | 100.00 | 0.00 |
| RDP80 | 100.00 | 0.00 | . | 100.00 | 0.00 |
| SINTAX50 | 100.00 | 0.00 | . | 100.00 | 0.00 |
| SINTAX80 | 100.00 | 0.00 | . | 100.00 | 0.00 |
| TOP | 100.00 | 0.00 | . | 100.00 | 0.00 |
| Q2_BLAST | 99.90 | 0.00 | . | 99.90 | 0.10 |
| Q2_VS | 99.90 | 0.00 | . | 99.90 | 0.10 |
| KNN | 99.80 | 0.00 | . | 99.80 | 0.20 |
| Q1 | 99.70 | 0.00 | . | 99.70 | 0.30 |
| BLCA | 95.20 | 0.00 | . | 95.20 | 4.80 |
| SPINGO | 0.00 | 0.00 | . | 0.00 | 100.00 |

**Table S51. TAXXI metrics computed for the dataset SP RDP ITS 99 at the class level.**

| Method | Accuracy | Misclassification Rate | Over-classification Rate | True Positive Rate | Under-classification Rate |
|---|---|---|---|---|---|
| BTOP | 100.00 | 0.00 | . | 100.00 | 0.00 |
| HiTaC | 100.00 | 0.00 | . | 100.00 | 0.00 |
| HiTaC_Filter | 100.00 | 0.00 | . | 100.00 | 0.00 |
| KTOP | 100.00 | 0.00 | . | 100.00 | 0.00 |
| Microclass | 100.00 | 0.00 | . | 100.00 | 0.00 |
| NBC50 | 100.00 | 0.00 | . | 100.00 | 0.00 |
| NBC80 | 100.00 | 0.00 | . | 100.00 | 0.00 |
| Q2_SK | 100.00 | 0.00 | . | 100.00 | 0.00 |
| RDP50 | 100.00 | 0.00 | . | 100.00 | 0.00 |
| RDP80 | 100.00 | 0.00 | . | 100.00 | 0.00 |
| SINTAX50 | 100.00 | 0.00 | . | 100.00 | 0.00 |
| SINTAX80 | 100.00 | 0.00 | . | 100.00 | 0.00 |
| TOP | 100.00 | 0.00 | . | 100.00 | 0.00 |
| CT1 | 99.90 | 0.10 | . | 99.90 | 0.00 |
| Q1 | 99.70 | 0.10 | . | 99.70 | 0.30 |
| Q2_BLAST | 99.70 | 0.20 | . | 99.70 | 0.10 |
| Q2_VS | 99.70 | 0.20 | . | 99.70 | 0.10 |
| CT2 | 99.70 | 0.30 | . | 99.70 | 0.10 |
| Metaxa2 | 99.60 | 0.00 | . | 99.60 | 0.40 |
| KNN | 97.80 | 0.00 | . | 97.80 | 2.20 |
| BLCA | 95.20 | 0.00 | . | 95.20 | 4.80 |
| SPINGO | 0.00 | 0.00 | . | 0.00 | 100.00 |

**Table S52. TAXXI metrics computed for the dataset SP RDP ITS 99 at the order level.**

| Method | Accuracy | Misclassification Rate | Over-classification Rate | True Positive Rate | Under-classification Rate |
|---|---|---|---|---|---|
| BTOP | 100.00 | 0.00 | . | 100.00 | 0.00 |
| HiTaC | 100.00 | 0.00 | . | 100.00 | 0.00 |
| HiTaC_Filter | 100.00 | 0.00 | . | 100.00 | 0.00 |
| Microclass | 100.00 | 0.00 | . | 100.00 | 0.00 |
| Q2_SK | 99.90 | 0.00 | . | 99.90 | 0.00 |
| SINTAX50 | 99.90 | 0.00 | . | 99.90 | 0.00 |
| SINTAX80 | 99.90 | 0.00 | . | 99.90 | 0.10 |
| KTOP | 99.90 | 0.10 | . | 99.90 | 0.00 |
| NBC50 | 99.90 | 0.10 | . | 99.90 | 0.00 |
| NBC80 | 99.90 | 0.10 | . | 99.90 | 0.00 |
| RDP50 | 99.90 | 0.10 | . | 99.90 | 0.00 |
| RDP80 | 99.90 | 0.10 | . | 99.90 | 0.00 |
| TOP | 99.90 | 0.10 | . | 99.90 | 0.00 |
| Q1 | 99.40 | 0.30 | . | 99.40 | 0.30 |
| CT1 | 99.30 | 0.60 | . | 99.30 | 0.10 |
| Q2_VS | 99.00 | 0.70 | . | 99.00 | 0.30 |
| Q2_BLAST | 98.90 | 0.80 | . | 98.90 | 0.30 |
| Metaxa2 | 98.70 | 0.00 | . | 98.70 | 1.30 |
| CT2 | 98.50 | 1.00 | . | 98.50 | 0.60 |
| BLCA | 95.20 | 0.00 | . | 95.20 | 4.80 |
| KNN | 93.20 | 0.00 | . | 93.20 | 6.80 |
| SPINGO | 0.00 | 0.00 | . | 0.00 | 100.00 |

**Table S53. TAXXI metrics computed for the dataset SP RDP ITS 99 at the family level.**

| Method | Accuracy | Misclassification Rate | Over-classification Rate | True Positive Rate | Under-classification Rate |
|---|---|---|---|---|---|
| BTOP | 100.00 | 0.00 | . | 100.00 | 0.00 |
| HiTaC | 100.00 | 0.00 | . | 100.00 | 0.00 |
| HiTaC_Filter | 99.90 | 0.00 | . | 99.90 | 0.00 |
| NBC80 | 99.90 | 0.00 | . | 99.90 | 0.10 |
| SINTAX50 | 99.90 | 0.00 | . | 99.90 | 0.10 |
| SPINGO | 99.90 | 0.00 | . | 99.90 | 0.10 |
| KTOP | 99.90 | 0.10 | . | 99.90 | 0.00 |
| Microclass | 99.90 | 0.10 | . | 99.90 | 0.00 |
| NBC50 | 99.90 | 0.10 | . | 99.90 | 0.00 |
| RDP50 | 99.90 | 0.10 | . | 99.90 | 0.00 |
| TOP | 99.90 | 0.10 | . | 99.90 | 0.00 |
| Q2_SK | 99.90 | 0.10 | . | 99.90 | 0.10 |
| RDP80 | 99.90 | 0.10 | . | 99.90 | 0.10 |
| SINTAX80 | 99.80 | 0.00 | . | 99.80 | 0.20 |
| Q1 | 99.10 | 0.60 | . | 99.10 | 0.30 |
| CT1 | 97.90 | 1.70 | . | 97.90 | 0.30 |
| Metaxa2 | 96.20 | 0.00 | . | 96.20 | 3.80 |
| Q2_VS | 96.10 | 2.30 | . | 96.10 | 1.60 |
| Q2_BLAST | 95.60 | 2.30 | . | 95.60 | 2.10 |
| BLCA | 95.10 | 0.00 | . | 95.10 | 4.80 |
| CT2 | 93.80 | 2.90 | . | 93.80 | 3.20 |
| KNN | 80.80 | 0.00 | . | 80.80 | 19.10 |

**Table S54. TAXXI metrics computed for the dataset SP RDP ITS 99 at the genus level.**

| Method | Accuracy | Misclassification Rate | Over-classification Rate | True Positive Rate | Under-classification Rate |
|---|---|---|---|---|---|
| HiTaC | 99.50 | 0.50 | . | 99.50 | 0.00 |
| TOP | 99.50 | 0.50 | . | 99.50 | 0.00 |
| BTOP | 99.40 | 0.60 | . | 99.40 | 0.00 |
| KTOP | 99.40 | 0.60 | . | 99.40 | 0.00 |
| Microclass | 99.40 | 0.60 | . | 99.40 | 0.00 |
| RDP50 | 99.30 | 0.60 | . | 99.30 | 0.10 |
| NBC50 | 99.30 | 0.70 | . | 99.30 | 0.10 |
| Q2_SK | 99.20 | 0.40 | . | 99.20 | 0.30 |
| SINTAX50 | 99.20 | 0.40 | . | 99.20 | 0.40 |
| SPINGO | 99.00 | 0.20 | . | 99.00 | 0.80 |
| RDP80 | 98.90 | 0.30 | . | 98.90 | 0.90 |
| HiTaC_Filter | 98.80 | 0.10 | . | 98.80 | 1.10 |
| NBC80 | 98.80 | 0.20 | . | 98.80 | 1.00 |
| SINTAX80 | 98.00 | 0.10 | . | 98.00 | 2.00 |
| Q1 | 95.50 | 3.30 | . | 95.50 | 1.20 |
| BLCA | 94.30 | 0.30 | . | 94.30 | 5.40 |
| CT1 | 92.20 | 6.00 | . | 92.20 | 1.70 |
| Metaxa2 | 89.00 | 0.30 | . | 89.00 | 10.60 |
| Q2_VS | 81.40 | 6.80 | . | 81.40 | 11.80 |
| Q2_BLAST | 80.20 | 7.70 | . | 80.20 | 12.20 |
| CT2 | 77.10 | 7.60 | . | 77.10 | 15.40 |
| KNN | 48.00 | 0.00 | . | 48.00 | 52.00 |

**Table S55. TAXXI metrics computed for the dataset SP RDP ITS 99 at the species level.**

| Method | Accuracy | Misclassification Rate | Over-classification Rate | True Positive Rate | Under-classification Rate |
|---|---|---|---|---|---|
| HiTaC | 73.00 | 10.60 | 100.00 | 89.40 | 0.00 |
| BTOP | 72.60 | 11.30 | 100.00 | 88.70 | 0.00 |
| Microclass | 72.30 | 11.40 | 100.00 | 88.60 | 0.00 |
| TOP | 72.30 | 11.40 | 100.00 | 88.60 | 0.00 |
| SINTAX50 | 72.20 | 6.60 | 70.40 | 83.60 | 9.70 |
| KTOP | 71.60 | 12.20 | 100.00 | 87.80 | 0.00 |
| SPINGO | 71.30 | 3.50 | 57.60 | 80.60 | 15.90 |
| Q2_SK | 70.10 | 8.40 | 74.90 | 82.00 | 9.60 |
| HiTaC_Filter | 70.00 | 2.70 | 49.00 | 77.80 | 19.50 |
| NBC50 | 69.70 | 13.30 | 91.20 | 84.10 | 2.60 |
| RDP50 | 69.60 | 13.30 | 89.50 | 83.70 | 3.00 |
| NBC80 | 68.60 | 7.00 | 64.30 | 78.50 | 14.50 |
| RDP80 | 68.50 | 6.40 | 63.60 | 78.30 | 15.30 |
| BLCA | 67.20 | 7.90 | 66.50 | 77.30 | 14.80 |
| SINTAX80 | 65.40 | 1.30 | 37.70 | 71.00 | 27.70 |
| Q1 | 39.40 | 21.70 | 65.70 | 45.20 | 33.10 |
| CT1 | 28.80 | 28.50 | 68.20 | 33.20 | 38.30 |
| Metaxa2 | 25.90 | 2.80 | 42.00 | 28.30 | 68.90 |
| Q2_VS | 8.80 | 2.10 | 4.50 | 8.90 | 89.10 |
| Q2_BLAST | 7.00 | 1.70 | 4.70 | 7.10 | 91.30 |
| CT2 | 3.60 | 0.90 | 3.30 | 3.60 | 95.50 |
| KNN | 0.00 | 0.00 | 0.00 | 0.00 | 100.00 |

**Table S56. TAXXI metrics computed for the dataset SP RDP ITS 100 at the phylum level.**

| Method | Accuracy | Misclassification Rate | Over-classification Rate | True Positive Rate | Under-classification Rate |
|---|---|---|---|---|---|
| BTOP | 100.00 | 0.00 | . | 100.00 | 0.00 |
| CT1 | 100.00 | 0.00 | . | 100.00 | 0.00 |
| CT2 | 100.00 | 0.00 | . | 100.00 | 0.00 |
| HiTaC | 100.00 | 0.00 | . | 100.00 | 0.00 |
| HiTaC_Filter | 100.00 | 0.00 | . | 100.00 | 0.00 |
| KTOP | 100.00 | 0.00 | . | 100.00 | 0.00 |
| Metaxa2 | 100.00 | 0.00 | . | 100.00 | 0.00 |
| Microclass | 100.00 | 0.00 | . | 100.00 | 0.00 |
| Q1 | 100.00 | 0.00 | . | 100.00 | 0.00 |
| Q2_BLAST | 100.00 | 0.00 | . | 100.00 | 0.00 |
| Q2_SK | 100.00 | 0.00 | . | 100.00 | 0.00 |
| Q2_VS | 100.00 | 0.00 | . | 100.00 | 0.00 |
| RDP50 | 100.00 | 0.00 | . | 100.00 | 0.00 |
| RDP80 | 100.00 | 0.00 | . | 100.00 | 0.00 |
| SINTAX50 | 100.00 | 0.00 | . | 100.00 | 0.00 |
| SINTAX80 | 100.00 | 0.00 | . | 100.00 | 0.00 |
| TOP | 100.00 | 0.00 | . | 100.00 | 0.00 |
| KNN | 99.80 | 0.00 | . | 99.80 | 0.20 |
| BLCA | 0.00 | 0.00 | . | 0.00 | 100.00 |
| SPINGO | 0.00 | 0.00 | . | 0.00 | 100.00 |

**Table S57. TAXXI metrics computed for the dataset SP RDP ITS 100 at the class level.**

| Method | Accuracy | Misclassification Rate | Over-classification Rate | True Positive Rate | Under-classification Rate |
|---|---|---|---|---|---|
| BTOP | 100.00 | 0.00 | . | 100.00 | 0.00 |
| HiTaC | 100.00 | 0.00 | . | 100.00 | 0.00 |
| HiTaC_Filter | 100.00 | 0.00 | . | 100.00 | 0.00 |
| KTOP | 100.00 | 0.00 | . | 100.00 | 0.00 |
| Microclass | 100.00 | 0.00 | . | 100.00 | 0.00 |
| Q1 | 100.00 | 0.00 | . | 100.00 | 0.00 |
| Q2_SK | 100.00 | 0.00 | . | 100.00 | 0.00 |
| RDP50 | 100.00 | 0.00 | . | 100.00 | 0.00 |
| RDP80 | 100.00 | 0.00 | . | 100.00 | 0.00 |
| SINTAX50 | 100.00 | 0.00 | . | 100.00 | 0.00 |
| SINTAX80 | 100.00 | 0.00 | . | 100.00 | 0.00 |
| TOP | 100.00 | 0.00 | . | 100.00 | 0.00 |
| Metaxa2 | 99.90 | 0.00 | . | 99.90 | 0.10 |
| CT1 | 99.90 | 0.10 | . | 99.90 | 0.00 |
| Q2_BLAST | 99.70 | 0.20 | . | 99.70 | 0.00 |
| Q2_VS | 99.70 | 0.20 | . | 99.70 | 0.00 |
| CT2 | 99.70 | 0.30 | . | 99.70 | 0.00 |
| KNN | 98.20 | 0.00 | . | 98.20 | 1.80 |
| BLCA | 0.00 | 0.00 | . | 0.00 | 100.00 |
| SPINGO | 0.00 | 0.00 | . | 0.00 | 100.00 |

**Table S58. TAXXI metrics computed for the dataset SP RDP ITS 100 at the order level.**

| Method | Accuracy | Misclassification Rate | Over-classification Rate | True Positive Rate | Under-classification Rate |
|---|---|---|---|---|---|
| BTOP | 100.00 | 0.00 | . | 100.00 | 0.00 |
| HiTaC | 100.00 | 0.00 | . | 100.00 | 0.00 |
| HiTaC_Filter | 100.00 | 0.00 | . | 100.00 | 0.00 |
| KTOP | 100.00 | 0.00 | . | 100.00 | 0.00 |
| Microclass | 100.00 | 0.00 | . | 100.00 | 0.00 |
| Q1 | 100.00 | 0.00 | . | 100.00 | 0.00 |
| Q2_SK | 100.00 | 0.00 | . | 100.00 | 0.00 |
| RDP50 | 100.00 | 0.00 | . | 100.00 | 0.00 |
| RDP80 | 100.00 | 0.00 | . | 100.00 | 0.00 |
| SINTAX50 | 100.00 | 0.00 | . | 100.00 | 0.00 |
| SINTAX80 | 100.00 | 0.00 | . | 100.00 | 0.00 |
| TOP | 100.00 | 0.00 | . | 100.00 | 0.00 |
| CT1 | 99.80 | 0.20 | . | 99.80 | 0.00 |
| Metaxa2 | 99.70 | 0.00 | . | 99.70 | 0.30 |
| Q2_VS | 99.00 | 0.60 | . | 99.00 | 0.30 |
| Q2_BLAST | 99.00 | 0.60 | . | 99.00 | 0.40 |
| CT2 | 98.70 | 0.80 | . | 98.70 | 0.50 |
| KNN | 94.40 | 0.00 | . | 94.40 | 5.60 |
| BLCA | 0.00 | 0.00 | . | 0.00 | 100.00 |
| SPINGO | 0.00 | 0.00 | . | 0.00 | 100.00 |

**Table S59. TAXXI metrics computed for the dataset SP RDP ITS 100 at the family level.**

| Method | Accuracy | Misclassification Rate | Over-classification Rate | True Positive Rate | Under-classification Rate |
|---|---|---|---|---|---|
| BTOP | 100.00 | 0.00 | . | 100.00 | 0.00 |
| HiTaC | 100.00 | 0.00 | . | 100.00 | 0.00 |
| HiTaC_Filter | 100.00 | 0.00 | . | 100.00 | 0.00 |
| KTOP | 100.00 | 0.00 | . | 100.00 | 0.00 |
| Microclass | 100.00 | 0.00 | . | 100.00 | 0.00 |
| Q2_SK | 100.00 | 0.00 | . | 100.00 | 0.00 |
| RDP50 | 100.00 | 0.00 | . | 100.00 | 0.00 |
| RDP80 | 100.00 | 0.00 | . | 100.00 | 0.00 |
| SINTAX50 | 100.00 | 0.00 | . | 100.00 | 0.00 |
| SINTAX80 | 100.00 | 0.00 | . | 100.00 | 0.00 |
| SPINGO | 100.00 | 0.00 | . | 100.00 | 0.00 |
| TOP | 100.00 | 0.00 | . | 100.00 | 0.00 |
| Q1 | 99.80 | 0.20 | . | 99.80 | 0.00 |
| CT1 | 99.50 | 0.50 | . | 99.50 | 0.10 |
| Metaxa2 | 99.20 | 0.00 | . | 99.20 | 0.80 |
| Q2_VS | 96.40 | 2.00 | . | 96.40 | 1.60 |
| Q2_BLAST | 96.10 | 2.10 | . | 96.10 | 1.80 |
| CT2 | 95.80 | 2.20 | . | 95.80 | 2.10 |
| KNN | 84.80 | 0.00 | . | 84.80 | 15.20 |
| BLCA | 0.00 | 0.00 | . | 0.00 | 100.00 |

**Table S60. TAXXI metrics computed for the dataset SP RDP ITS 100 at the genus level.**

| Method | Accuracy | Misclassification Rate | Over-classification Rate | True Positive Rate | Under-classification Rate |
|---|---|---|---|---|---|
| BTOP | 100.00 | 0.00 | . | 100.00 | 0.00 |
| HiTaC | 100.00 | 0.00 | . | 100.00 | 0.00 |
| HiTaC_Filter | 100.00 | 0.00 | . | 100.00 | 0.00 |
| KTOP | 100.00 | 0.00 | . | 100.00 | 0.00 |
| Microclass | 100.00 | 0.00 | . | 100.00 | 0.00 |
| Q2_SK | 100.00 | 0.00 | . | 100.00 | 0.00 |
| RDP50 | 100.00 | 0.00 | . | 100.00 | 0.00 |
| SINTAX50 | 100.00 | 0.00 | . | 100.00 | 0.00 |
| SPINGO | 100.00 | 0.00 | . | 100.00 | 0.00 |
| TOP | 100.00 | 0.00 | . | 100.00 | 0.00 |
| RDP80 | 99.90 | 0.00 | . | 99.90 | 0.10 |
| SINTAX80 | 99.80 | 0.00 | . | 99.80 | 0.20 |
| Q1 | 99.00 | 0.80 | . | 99.00 | 0.20 |
| CT1 | 98.20 | 1.50 | . | 98.20 | 0.30 |
| Metaxa2 | 97.70 | 0.00 | . | 97.70 | 2.30 |
| Q2_VS | 84.30 | 6.20 | . | 84.30 | 9.50 |
| Q2_BLAST | 83.60 | 6.70 | . | 83.60 | 9.70 |
| CT2 | 82.90 | 6.60 | . | 82.90 | 10.50 |
| KNN | 57.60 | 0.00 | . | 57.60 | 42.40 |
| BLCA | 0.00 | 0.00 | . | 0.00 | 100.00 |

**Table S61. TAXXI metrics computed for the dataset SP RDP ITS 100 at the species level.**

| Method | Accuracy | Misclassification Rate | Over-classification Rate | True Positive Rate | Under-classification Rate |
|---|---|---|---|---|---|
| BTOP | 100.00 | 0.00 | . | 100.00 | 0.00 |
| HiTaC | 100.00 | 0.00 | . | 100.00 | 0.00 |
| TOP | 99.60 | 0.40 | . | 99.60 | 0.00 |
| KTOP | 99.40 | 0.60 | . | 99.40 | 0.00 |
| Microclass | 99.40 | 0.60 | . | 99.40 | 0.00 |
| HiTaC_Filter | 97.80 | 0.00 | . | 97.80 | 2.20 |
| SINTAX50 | 97.50 | 0.40 | . | 97.50 | 2.10 |
| SPINGO | 96.80 | 0.00 | . | 96.80 | 3.20 |
| RDP50 | 96.20 | 2.90 | . | 96.20 | 0.90 |
| Q2_SK | 95.30 | 0.80 | . | 95.30 | 3.80 |
| RDP80 | 93.00 | 1.50 | . | 93.00 | 5.50 |
| SINTAX80 | 86.20 | 0.00 | . | 86.20 | 13.80 |
| Q1 | 83.70 | 7.80 | . | 83.70 | 8.50 |
| CT1 | 82.70 | 8.70 | . | 82.70 | 8.60 |
| Metaxa2 | 78.00 | 0.00 | . | 78.00 | 22.00 |
| Q2_VS | 10.90 | 1.30 | . | 10.90 | 87.80 |
| Q2_BLAST | 9.30 | 1.30 | . | 9.30 | 89.40 |
| CT2 | 8.50 | 1.10 | . | 8.50 | 90.40 |
| KNN | 0.10 | 0.00 | . | 0.10 | 99.90 |
| BLCA | 0.00 | 0.00 | . | 0.00 | 100.00 |

**Table S62. Machine learning metrics computed for the dataset SP RDP ITS 90 at the phylum level.**

| Method | Accuracy | Balanced Accuracy | F1-score Micro | F1-score Macro | F1-score Weighted | Precision Micro | Precision Macro | Precision Weighted | Recall Micro | Recall Macro | Recall Weighted | Jaccard Micro | Jaccard Macro | Jaccard Weighted |
|---|---|---|---|---|---|---|---|---|---|---|---|---|---|---|
| BTOP | 100.00 | 100.00 | 100.00 | 100.00 | 100.00 | 100.00 | 100.00 | 100.00 | 100.00 | 100.00 | 100.00 | 100.00 | 100.00 | 100.00 |
| CT1 | 100.00 | 100.00 | 100.00 | 100.00 | 100.00 | 100.00 | 100.00 | 100.00 | 100.00 | 100.00 | 100.00 | 100.00 | 100.00 | 100.00 |
| CT2 | 100.00 | 100.00 | 100.00 | 100.00 | 100.00 | 100.00 | 100.00 | 100.00 | 100.00 | 100.00 | 100.00 | 100.00 | 100.00 | 100.00 |
| HiTaC | 100.00 | 100.00 | 100.00 | 100.00 | 100.00 | 100.00 | 100.00 | 100.00 | 100.00 | 100.00 | 100.00 | 100.00 | 100.00 | 100.00 |
| HiTaC_Filter | 100.00 | 100.00 | 100.00 | 100.00 | 100.00 | 100.00 | 100.00 | 100.00 | 100.00 | 100.00 | 100.00 | 100.00 | 100.00 | 100.00 |
| KTOP | 100.00 | 100.00 | 100.00 | 100.00 | 100.00 | 100.00 | 100.00 | 100.00 | 100.00 | 100.00 | 100.00 | 100.00 | 100.00 | 100.00 |
| Metaxa2 | 100.00 | 100.00 | 100.00 | 100.00 | 100.00 | 100.00 | 100.00 | 100.00 | 100.00 | 100.00 | 100.00 | 100.00 | 100.00 | 100.00 |
| Microclass | 100.00 | 100.00 | 100.00 | 100.00 | 100.00 | 100.00 | 100.00 | 100.00 | 100.00 | 100.00 | 100.00 | 100.00 | 100.00 | 100.00 |
| NBC50 | 100.00 | 100.00 | 100.00 | 100.00 | 100.00 | 100.00 | 100.00 | 100.00 | 100.00 | 100.00 | 100.00 | 100.00 | 100.00 | 100.00 |
| NBC80 | 100.00 | 100.00 | 100.00 | 100.00 | 100.00 | 100.00 | 100.00 | 100.00 | 100.00 | 100.00 | 100.00 | 100.00 | 100.00 | 100.00 |
| Q2_SK | 100.00 | 100.00 | 100.00 | 100.00 | 100.00 | 100.00 | 100.00 | 100.00 | 100.00 | 100.00 | 100.00 | 100.00 | 100.00 | 100.00 |
| RDP50 | 100.00 | 100.00 | 100.00 | 100.00 | 100.00 | 100.00 | 100.00 | 100.00 | 100.00 | 100.00 | 100.00 | 100.00 | 100.00 | 100.00 |
| RDP80 | 100.00 | 100.00 | 100.00 | 100.00 | 100.00 | 100.00 | 100.00 | 100.00 | 100.00 | 100.00 | 100.00 | 100.00 | 100.00 | 100.00 |
| SINTAX50 | 100.00 | 100.00 | 100.00 | 100.00 | 100.00 | 100.00 | 100.00 | 100.00 | 100.00 | 100.00 | 100.00 | 100.00 | 100.00 | 100.00 |
| SINTAX80 | 100.00 | 100.00 | 100.00 | 100.00 | 100.00 | 100.00 | 100.00 | 100.00 | 100.00 | 100.00 | 100.00 | 100.00 | 100.00 | 100.00 |
| TOP | 100.00 | 100.00 | 100.00 | 100.00 | 100.00 | 100.00 | 100.00 | 100.00 | 100.00 | 100.00 | 100.00 | 100.00 | 100.00 | 100.00 |
| KNN | 99.90 | 98.15 | 99.90 | 74.29 | 99.95 | 99.90 | 75.00 | 100.00 | 99.90 | 73.61 | 99.90 | 99.79 | 73.61 | 99.90 |
| Q2_VS | 99.84 | 99.91 | 99.84 | 74.97 | 99.92 | 99.84 | 75.00 | 100.00 | 99.84 | 74.93 | 99.84 | 99.69 | 74.93 | 99.84 |
| Q2_BLAST | 99.74 | 99.85 | 99.74 | 74.94 | 99.87 | 99.74 | 75.00 | 100.00 | 99.74 | 74.89 | 99.74 | 99.48 | 74.89 | 99.74 |
| Q1 | 92.76 | 95.16 | 92.76 | 73.12 | 96.24 | 92.76 | 75.00 | 100.00 | 92.76 | 71.37 | 92.76 | 86.51 | 71.37 | 92.76 |
| BLCA | 92.51 | 82.79 | 92.51 | 67.31 | 96.03 | 92.51 | 75.00 | 100.00 | 92.51 | 62.09 | 92.51 | 86.06 | 62.09 | 92.51 |
| SPINGO | 0.00 | 0.00 | 0.00 | 0.00 | 0.00 | 0.00 | 0.00 | 0.00 | 0.00 | 0.00 | 0.00 | 0.00 | 0.00 | 0.00 |

**Table S63. Machine learning metrics computed for the dataset SP RDP ITS 90 at the class level.**

| Method | Accuracy | Balanced Accuracy | F1-score Micro | F1-score Macro | F1-score Weighted | Precision Micro | Precision Macro | Precision Weighted | Recall Micro | Recall Macro | Recall Weighted | Jaccard Micro | Jaccard Macro | Jaccard Weighted |
|---|---|---|---|---|---|---|---|---|---|---|---|---|---|---|
| HiTaC | 99.95 | 99.97 | 99.95 | 99.97 | 99.95 | 99.95 | 99.98 | 99.95 | 99.95 | 99.97 | 99.95 | 99.90 | 99.95 | 99.90 |
| HiTaC_Filter | 99.95 | 99.97 | 99.95 | 99.97 | 99.95 | 99.95 | 99.98 | 99.95 | 99.95 | 99.97 | 99.95 | 99.90 | 99.95 | 99.90 |
| CT2 | 99.90 | 99.95 | 99.90 | 99.95 | 99.90 | 99.90 | 99.95 | 99.90 | 99.90 | 99.95 | 99.90 | 99.79 | 99.90 | 99.79 |
| BTOP | 99.85 | 99.04 | 99.85 | 99.12 | 99.85 | 99.85 | 99.33 | 99.86 | 99.85 | 99.04 | 99.85 | 99.70 | 98.37 | 99.71 |
| Microclass | 99.84 | 99.04 | 99.84 | 99.12 | 99.84 | 99.84 | 99.33 | 99.86 | 99.84 | 99.04 | 99.84 | 99.69 | 98.37 | 99.70 |
| Q2_SK | 99.84 | 99.04 | 99.84 | 99.12 | 99.84 | 99.84 | 99.33 | 99.86 | 99.84 | 99.04 | 99.84 | 99.69 | 98.37 | 99.70 |
| Q2_VS | 99.79 | 99.92 | 99.79 | 94.69 | 99.87 | 99.79 | 94.72 | 99.95 | 99.79 | 94.66 | 99.79 | 99.59 | 94.64 | 99.74 |
| TOP | 99.74 | 99.01 | 99.74 | 99.02 | 99.74 | 99.74 | 99.17 | 99.76 | 99.74 | 99.01 | 99.74 | 99.48 | 98.17 | 99.50 |
| Q2_BLAST | 99.69 | 99.88 | 99.69 | 94.67 | 99.82 | 99.69 | 94.72 | 99.95 | 99.69 | 94.62 | 99.69 | 99.38 | 94.61 | 99.64 |
| Metaxa2 | 99.69 | 98.95 | 99.69 | 99.05 | 99.69 | 99.69 | 99.28 | 99.70 | 99.69 | 98.95 | 99.69 | 99.38 | 98.23 | 99.39 |
| RDP50 | 99.59 | 98.93 | 99.59 | 98.83 | 99.59 | 99.59 | 98.88 | 99.61 | 99.59 | 98.93 | 99.59 | 99.18 | 97.81 | 99.20 |
| KTOP | 99.53 | 98.90 | 99.53 | 98.90 | 99.53 | 99.53 | 99.04 | 99.55 | 99.53 | 98.90 | 99.53 | 99.07 | 97.94 | 99.09 |
| NBC80 | 99.53 | 98.90 | 99.53 | 94.17 | 99.74 | 99.53 | 94.72 | 99.95 | 99.53 | 93.69 | 99.53 | 99.07 | 93.68 | 99.48 |
| RDP80 | 99.53 | 98.90 | 99.53 | 94.17 | 99.74 | 99.53 | 94.72 | 99.95 | 99.53 | 93.69 | 99.53 | 99.07 | 93.68 | 99.48 |
| SINTAX80 | 99.53 | 98.90 | 99.53 | 94.17 | 99.74 | 99.53 | 94.72 | 99.95 | 99.53 | 93.69 | 99.53 | 99.07 | 93.68 | 99.48 |
| SINTAX50 | 99.53 | 98.90 | 99.53 | 94.14 | 99.66 | 99.53 | 94.67 | 99.79 | 99.53 | 93.69 | 99.53 | 99.07 | 93.63 | 99.33 |
| CT1 | 99.53 | 98.90 | 99.53 | 93.79 | 99.56 | 99.53 | 94.00 | 99.60 | 99.53 | 93.69 | 99.53 | 99.07 | 92.96 | 99.14 |
| NBC50 | 99.53 | 98.90 | 99.53 | 93.78 | 99.59 | 99.53 | 93.96 | 99.65 | 99.53 | 93.69 | 99.53 | 99.07 | 92.92 | 99.19 |
| KNN | 98.60 | 91.56 | 98.60 | 88.95 | 99.15 | 98.60 | 94.72 | 99.95 | 98.60 | 86.74 | 98.60 | 97.25 | 86.73 | 98.55 |
| Q1 | 92.61 | 92.46 | 92.61 | 90.40 | 96.03 | 92.61 | 94.10 | 99.86 | 92.61 | 87.59 | 92.61 | 86.24 | 86.96 | 92.47 |
| BLCA | 92.35 | 91.80 | 92.35 | 90.00 | 95.71 | 92.35 | 94.10 | 99.85 | 92.35 | 86.96 | 92.35 | 85.79 | 86.33 | 92.21 |
| SPINGO | 0.00 | 0.00 | 0.00 | 0.00 | 0.00 | 0.00 | 0.00 | 0.00 | 0.00 | 0.00 | 0.00 | 0.00 | 0.00 | 0.00 |

**Table S64. Machine learning metrics computed for the dataset SP RDP ITS 90 at the order level.**

| Method | Accuracy | Balanced Accuracy | F1-score Micro | F1-score Macro | F1-score Weighted | Precision Micro | Precision Macro | Precision Weighted | Recall Micro | Recall Macro | Recall Weighted | Jaccard Micro | Jaccard Macro | Jaccard Weighted |
|---|---|---|---|---|---|---|---|---|---|---|---|---|---|---|
| Microclass | 98.24 | 94.21 | 98.24 | 92.80 | 98.29 | 98.24 | 93.34 | 98.46 | 98.24 | 92.66 | 98.24 | 96.55 | 91.38 | 97.24 |
| HiTaC | 98.19 | 90.49 | 98.19 | 89.42 | 98.07 | 98.19 | 90.38 | 98.07 | 98.19 | 89.01 | 98.19 | 96.45 | 88.04 | 97.10 |
| HiTaC_Filter | 98.19 | 90.49 | 98.19 | 88.01 | 98.17 | 98.19 | 88.99 | 98.26 | 98.19 | 87.57 | 98.19 | 96.45 | 86.69 | 97.29 |
| Q2_VS | 98.14 | 89.79 | 98.14 | 87.08 | 98.20 | 98.14 | 87.39 | 98.30 | 98.14 | 86.89 | 98.14 | 96.35 | 86.03 | 97.34 |
| TOP | 98.09 | 94.09 | 98.09 | 92.65 | 98.14 | 98.09 | 93.17 | 98.31 | 98.09 | 92.55 | 98.09 | 96.25 | 91.09 | 96.95 |
| Q2_SK | 98.09 | 93.78 | 98.09 | 91.13 | 98.28 | 98.09 | 91.93 | 98.59 | 98.09 | 90.76 | 98.09 | 96.25 | 89.59 | 97.22 |
| BTOP | 98.07 | 93.84 | 98.07 | 92.57 | 98.10 | 98.07 | 93.31 | 98.28 | 98.07 | 92.30 | 98.07 | 96.21 | 90.99 | 96.89 |
| Q2_BLAST | 97.98 | 91.02 | 97.98 | 88.47 | 98.11 | 97.98 | 89.06 | 98.30 | 97.98 | 88.08 | 97.98 | 96.05 | 87.27 | 97.13 |
| KTOP | 97.93 | 94.11 | 97.93 | 92.47 | 97.98 | 97.93 | 92.83 | 98.19 | 97.93 | 92.56 | 97.93 | 95.95 | 90.77 | 96.67 |
| RDP50 | 97.93 | 94.04 | 97.93 | 91.07 | 98.02 | 97.93 | 91.55 | 98.26 | 97.93 | 91.00 | 97.93 | 95.95 | 89.46 | 96.74 |
| CT1 | 97.88 | 94.02 | 97.88 | 90.99 | 97.95 | 97.88 | 91.42 | 98.16 | 97.88 | 90.99 | 97.88 | 95.85 | 89.32 | 96.63 |
| NBC50 | 97.88 | 94.01 | 97.88 | 91.14 | 98.04 | 97.88 | 91.70 | 98.35 | 97.88 | 90.98 | 97.88 | 95.85 | 89.59 | 96.78 |
| CT2 | 97.88 | 86.64 | 97.88 | 84.42 | 97.88 | 97.88 | 85.58 | 98.01 | 97.88 | 83.85 | 97.88 | 95.85 | 82.78 | 96.84 |
| RDP80 | 97.83 | 93.74 | 97.83 | 91.34 | 98.25 | 97.83 | 92.35 | 98.80 | 97.83 | 90.71 | 97.83 | 95.75 | 89.97 | 97.18 |
| SINTAX50 | 97.83 | 93.62 | 97.83 | 91.20 | 98.18 | 97.83 | 92.22 | 98.67 | 97.83 | 90.60 | 97.83 | 95.75 | 89.73 | 97.05 |
| NBC80 | 97.78 | 93.32 | 97.78 | 91.11 | 98.22 | 97.78 | 92.35 | 98.80 | 97.78 | 90.31 | 97.78 | 95.65 | 89.56 | 97.13 |
| Metaxa2 | 97.21 | 92.23 | 97.21 | 89.45 | 97.18 | 97.21 | 91.13 | 97.88 | 97.21 | 89.26 | 97.21 | 94.57 | 87.38 | 95.69 |
| SINTAX80 | 97.11 | 89.60 | 97.11 | 87.75 | 97.75 | 97.11 | 89.12 | 98.55 | 97.11 | 86.71 | 97.11 | 94.37 | 85.97 | 96.46 |
| KNN | 93.23 | 68.33 | 93.23 | 68.02 | 94.70 | 93.23 | 72.18 | 96.81 | 93.23 | 66.12 | 93.23 | 87.32 | 65.74 | 92.80 |
| Q1 | 91.37 | 87.92 | 91.37 | 87.73 | 94.68 | 91.37 | 92.02 | 98.75 | 91.37 | 85.09 | 91.37 | 84.11 | 84.08 | 90.71 |
| BLCA | 90.44 | 86.70 | 90.44 | 86.87 | 93.71 | 90.44 | 91.73 | 98.39 | 90.44 | 83.90 | 90.44 | 82.55 | 82.73 | 89.49 |
| SPINGO | 0.00 | 0.00 | 0.00 | 0.00 | 0.00 | 0.00 | 0.00 | 0.00 | 0.00 | 0.00 | 0.00 | 0.00 | 0.00 | 0.00 |

Table S65. Machine learning metrics computed for the dataset SP RDP ITS 90 at the family level.

| Method | Accuracy | Balanced Accuracy | F1-score Micro | F1-score Macro | F1-score Weighted | Precision Micro | Precision Macro | Precision Weighted | Recall Micro | Recall Macro | Recall Weighted | Jaccard Micro | Jaccard Macro | Jaccard Weighted |
|---|---|---|---|---|---|---|---|---|---|---|---|---|---|---|
| HiTaC | 95.09 | 87.88 | 95.09 | 84.95 | 94.86 | 95.09 | 85.89 | 95.51 | 95.09 | 85.68 | 95.09 | 90.64 | 82.74 | 92.60 |
| HiTaC_Filter | 93.95 | 86.27 | 93.95 | 83.33 | 94.47 | 93.95 | 85.37 | 96.39 | 93.95 | 83.60 | 93.95 | 88.60 | 81.18 | 92.40 |
| TOP | 93.90 | 89.21 | 93.90 | 84.49 | 93.77 | 93.90 | 85.44 | 95.14 | 93.90 | 85.91 | 93.90 | 88.50 | 82.01 | 90.90 |
| Microclass | 93.85 | 89.90 | 93.85 | 84.35 | 93.92 | 93.85 | 84.74 | 95.02 | 93.85 | 85.52 | 93.85 | 88.41 | 81.67 | 90.87 |
| BTOP | 93.75 | 88.95 | 93.75 | 84.30 | 93.87 | 93.75 | 85.58 | 95.38 | 93.75 | 85.65 | 93.75 | 88.23 | 81.39 | 90.89 |
| KTOP | 93.44 | 88.40 | 93.44 | 83.12 | 93.18 | 93.44 | 84.02 | 94.61 | 93.44 | 84.60 | 93.44 | 87.68 | 80.41 | 89.98 |
| CT1 | 93.33 | 87.73 | 93.33 | 81.73 | 93.23 | 93.33 | 82.41 | 94.63 | 93.33 | 83.45 | 93.33 | 87.50 | 78.84 | 90.00 |
| RDP50 | 93.23 | 88.85 | 93.23 | 84.01 | 93.31 | 93.23 | 85.26 | 95.12 | 93.23 | 85.03 | 93.23 | 87.32 | 81.34 | 90.13 |
| NBC50 | 93.18 | 88.83 | 93.18 | 84.08 | 93.40 | 93.18 | 85.72 | 95.46 | 93.18 | 85.02 | 93.18 | 87.23 | 81.54 | 90.37 |
| Q2_SK | 93.13 | 89.02 | 93.13 | 85.43 | 93.92 | 93.13 | 87.32 | 96.14 | 93.13 | 85.73 | 93.13 | 87.14 | 82.64 | 90.88 |
| SINTAX50 | 92.66 | 86.44 | 92.66 | 82.90 | 93.46 | 92.66 | 85.14 | 95.97 | 92.66 | 83.24 | 92.66 | 86.33 | 80.39 | 90.64 |
| Q2_VS | 91.73 | 81.38 | 91.73 | 76.48 | 92.00 | 91.73 | 78.97 | 94.33 | 91.73 | 77.89 | 91.73 | 84.73 | 73.63 | 89.32 |
| NBC80 | 91.63 | 86.39 | 91.63 | 83.79 | 92.90 | 91.63 | 86.47 | 96.00 | 91.63 | 83.71 | 91.63 | 84.55 | 81.19 | 90.04 |
| RDP80 | 91.52 | 86.51 | 91.52 | 83.78 | 92.80 | 91.52 | 86.42 | 95.97 | 91.52 | 83.82 | 91.52 | 84.37 | 81.30 | 89.97 |
| Q2_BLAST | 91.37 | 82.22 | 91.37 | 77.09 | 91.90 | 91.37 | 78.94 | 93.71 | 91.37 | 78.21 | 91.37 | 84.11 | 74.17 | 88.85 |
| SPINGO | 91.27 | 86.17 | 91.27 | 83.28 | 92.74 | 91.27 | 86.36 | 95.99 | 91.27 | 82.98 | 91.27 | 83.94 | 80.34 | 89.64 |
| CT2 | 91.27 | 79.08 | 91.27 | 74.12 | 91.51 | 91.27 | 76.16 | 93.54 | 91.27 | 75.22 | 91.27 | 83.94 | 71.10 | 88.62 |
| Metaxa2 | 90.49 | 83.06 | 90.49 | 79.90 | 90.85 | 90.49 | 82.14 | 93.72 | 90.49 | 80.48 | 90.49 | 82.63 | 76.74 | 87.22 |
| SINTAX80 | 87.65 | 77.13 | 87.65 | 77.70 | 90.33 | 87.65 | 83.05 | 95.64 | 87.65 | 76.15 | 87.65 | 78.01 | 74.28 | 86.71 |
| Q1 | 87.29 | 81.03 | 87.29 | 79.45 | 90.37 | 87.29 | 83.41 | 95.34 | 87.29 | 78.03 | 87.29 | 77.44 | 75.12 | 85.07 |
| BLCA | 86.93 | 81.64 | 86.93 | 79.39 | 89.98 | 86.93 | 83.12 | 95.21 | 86.93 | 78.61 | 86.93 | 76.87 | 75.13 | 84.88 |
| KNN | 80.16 | 53.99 | 80.16 | 55.66 | 83.55 | 80.16 | 61.78 | 90.70 | 80.16 | 53.31 | 80.16 | 66.88 | 52.70 | 79.56 |

**Table S66. Machine learning metrics computed for the dataset SP RDP ITS 90 at the genus level.**

| Method | Accuracy | Balanced Accuracy | F1-score Micro | F1-score Macro | F1-score Weighted | Precision Micro | Precision Macro | Precision Weighted | Recall Micro | Recall Macro | Recall Weighted | Jaccard Micro | Jaccard Macro | Jaccard Weighted |
|---|---|---|---|---|---|---|---|---|---|---|---|---|---|---|
| HiTaC | 69.46 | 55.89 | 69.46 | 43.88 | 65.82 | 69.46 | 43.74 | 65.51 | 69.46 | 47.45 | 69.46 | 53.21 | 41.25 | 61.51 |
| BTOP | 63.65 | 57.26 | 63.65 | 43.26 | 63.08 | 63.65 | 44.23 | 68.20 | 63.65 | 45.81 | 63.65 | 46.68 | 40.44 | 57.69 |
| Microclass | 62.74 | 56.93 | 62.74 | 42.74 | 61.97 | 62.74 | 43.30 | 64.98 | 62.74 | 45.45 | 62.74 | 45.71 | 39.92 | 56.97 |
| KTOP | 62.43 | 56.09 | 62.43 | 42.29 | 61.26 | 62.43 | 43.13 | 64.00 | 62.43 | 45.14 | 62.43 | 45.38 | 39.43 | 56.14 |
| TOP | 62.38 | 56.63 | 62.38 | 42.43 | 61.28 | 62.38 | 43.03 | 64.04 | 62.38 | 45.30 | 62.38 | 45.32 | 39.73 | 56.32 |
| CT1 | 61.45 | 51.53 | 61.45 | 38.34 | 59.95 | 61.45 | 38.46 | 61.71 | 61.45 | 41.55 | 61.45 | 44.35 | 35.57 | 54.99 |
| NBC50 | 61.34 | 55.04 | 61.34 | 44.13 | 60.65 | 61.34 | 45.57 | 64.31 | 61.34 | 46.25 | 61.34 | 44.24 | 41.25 | 55.92 |
| RDP50 | 61.09 | 54.83 | 61.09 | 44.11 | 60.59 | 61.09 | 45.41 | 64.37 | 61.09 | 46.27 | 61.09 | 43.97 | 41.23 | 55.70 |
| HiTaC_Filter | 59.95 | 46.92 | 59.95 | 42.51 | 58.84 | 59.95 | 43.94 | 63.32 | 59.95 | 44.36 | 59.95 | 42.80 | 39.97 | 54.80 |
| Q2_SK | 59.28 | 53.63 | 59.28 | 43.23 | 59.27 | 59.28 | 44.92 | 64.77 | 59.28 | 45.25 | 59.28 | 42.12 | 40.56 | 54.52 |
| Q1 | 58.81 | 52.45 | 58.81 | 41.91 | 59.78 | 58.81 | 44.87 | 67.48 | 58.81 | 43.20 | 58.81 | 41.65 | 38.49 | 53.64 |
| SINTAX50 | 57.52 | 49.40 | 57.52 | 42.74 | 57.55 | 57.52 | 44.55 | 61.66 | 57.52 | 44.58 | 57.52 | 40.37 | 40.05 | 53.01 |
| Metaxa2 | 56.74 | 47.63 | 56.74 | 39.35 | 56.82 | 56.74 | 41.43 | 62.19 | 56.74 | 41.02 | 56.74 | 39.61 | 36.75 | 51.90 |
| BLCA | 56.23 | 49.12 | 56.23 | 40.17 | 57.93 | 56.23 | 43.20 | 66.30 | 56.23 | 41.03 | 56.23 | 39.11 | 36.97 | 52.27 |
| Q2_VS | 55.50 | 42.52 | 55.50 | 34.92 | 53.30 | 55.50 | 34.73 | 54.64 | 55.50 | 38.45 | 55.50 | 38.41 | 32.55 | 49.45 |
| SPINGO | 53.39 | 47.62 | 53.39 | 41.90 | 54.43 | 53.39 | 44.16 | 60.34 | 53.39 | 43.07 | 53.39 | 36.41 | 39.25 | 49.70 |
| Q2_BLAST | 53.13 | 40.53 | 53.13 | 33.79 | 51.69 | 53.13 | 34.25 | 54.26 | 53.13 | 36.73 | 53.13 | 36.17 | 31.23 | 47.16 |
| NBC80 | 52.66 | 46.40 | 52.66 | 41.73 | 53.56 | 52.66 | 44.34 | 60.25 | 52.66 | 42.71 | 52.66 | 35.74 | 39.22 | 49.31 |
| RDP80 | 52.61 | 46.95 | 52.61 | 42.37 | 53.64 | 52.61 | 44.98 | 60.72 | 52.61 | 43.21 | 52.61 | 35.69 | 39.98 | 49.42 |
| CT2 | 52.56 | 37.55 | 52.56 | 31.00 | 50.70 | 52.56 | 31.10 | 52.81 | 52.56 | 34.26 | 52.56 | 35.65 | 28.57 | 46.47 |
| SINTAX80 | 44.08 | 34.72 | 44.08 | 33.29 | 45.93 | 44.08 | 36.60 | 53.02 | 44.08 | 33.28 | 44.08 | 28.27 | 31.16 | 41.95 |
| KNN | 31.01 | 16.11 | 31.01 | 16.20 | 33.47 | 31.01 | 18.32 | 39.74 | 31.01 | 15.80 | 31.01 | 18.35 | 14.80 | 29.76 |

**Table S67. Machine learning metrics computed for the dataset SP RDP ITS 90 at the species level.**

| Method | Accuracy | Balanced Accuracy | F1-score Micro | F1-score Macro | F1-score Weighted | Precision Micro | Precision Macro | Precision Weighted | Recall Micro | Recall Macro | Recall Weighted | Jaccard Micro | Jaccard Macro | Jaccard Weighted |
|---|---|---|---|---|---|---|---|---|---|---|---|---|---|---|
| BLCA | 0.00 | 0.00 | 0.00 | 0.00 | 0.00 | 0.00 | 0.00 | 0.00 | 0.00 | 0.00 | 0.00 | 0.00 | 0.00 | 0.00 |
| BTOP | 0.00 | 0.00 | 0.00 | 0.00 | 0.00 | 0.00 | 0.00 | 0.00 | 0.00 | 0.00 | 0.00 | 0.00 | 0.00 | 0.00 |
| CT1 | 0.00 | 0.00 | 0.00 | 0.00 | 0.00 | 0.00 | 0.00 | 0.00 | 0.00 | 0.00 | 0.00 | 0.00 | 0.00 | 0.00 |
| CT2 | 0.00 | 0.00 | 0.00 | 0.00 | 0.00 | 0.00 | 0.00 | 0.00 | 0.00 | 0.00 | 0.00 | 0.00 | 0.00 | 0.00 |
| HiTaC | 0.00 | 0.00 | 0.00 | 0.00 | 0.00 | 0.00 | 0.00 | 0.00 | 0.00 | 0.00 | 0.00 | 0.00 | 0.00 | 0.00 |
| HiTaC_Filter | 0.00 | 0.00 | 0.00 | 0.00 | 0.00 | 0.00 | 0.00 | 0.00 | 0.00 | 0.00 | 0.00 | 0.00 | 0.00 | 0.00 |
| KNN | 0.00 | 0.00 | 0.00 | 0.00 | 0.00 | 0.00 | 0.00 | 0.00 | 0.00 | 0.00 | 0.00 | 0.00 | 0.00 | 0.00 |
| KTOP | 0.00 | 0.00 | 0.00 | 0.00 | 0.00 | 0.00 | 0.00 | 0.00 | 0.00 | 0.00 | 0.00 | 0.00 | 0.00 | 0.00 |
| Metaxa2 | 0.00 | 0.00 | 0.00 | 0.00 | 0.00 | 0.00 | 0.00 | 0.00 | 0.00 | 0.00 | 0.00 | 0.00 | 0.00 | 0.00 |
| Microclass | 0.00 | 0.00 | 0.00 | 0.00 | 0.00 | 0.00 | 0.00 | 0.00 | 0.00 | 0.00 | 0.00 | 0.00 | 0.00 | 0.00 |
| NBC50 | 0.00 | 0.00 | 0.00 | 0.00 | 0.00 | 0.00 | 0.00 | 0.00 | 0.00 | 0.00 | 0.00 | 0.00 | 0.00 | 0.00 |
| NBC80 | 0.00 | 0.00 | 0.00 | 0.00 | 0.00 | 0.00 | 0.00 | 0.00 | 0.00 | 0.00 | 0.00 | 0.00 | 0.00 | 0.00 |
| Q1 | 0.00 | 0.00 | 0.00 | 0.00 | 0.00 | 0.00 | 0.00 | 0.00 | 0.00 | 0.00 | 0.00 | 0.00 | 0.00 | 0.00 |
| Q2_BLAST | 0.00 | 0.00 | 0.00 | 0.00 | 0.00 | 0.00 | 0.00 | 0.00 | 0.00 | 0.00 | 0.00 | 0.00 | 0.00 | 0.00 |
| Q2_SK | 0.00 | 0.00 | 0.00 | 0.00 | 0.00 | 0.00 | 0.00 | 0.00 | 0.00 | 0.00 | 0.00 | 0.00 | 0.00 | 0.00 |
| Q2_VS | 0.00 | 0.00 | 0.00 | 0.00 | 0.00 | 0.00 | 0.00 | 0.00 | 0.00 | 0.00 | 0.00 | 0.00 | 0.00 | 0.00 |
| RDP50 | 0.00 | 0.00 | 0.00 | 0.00 | 0.00 | 0.00 | 0.00 | 0.00 | 0.00 | 0.00 | 0.00 | 0.00 | 0.00 | 0.00 |
| RDP80 | 0.00 | 0.00 | 0.00 | 0.00 | 0.00 | 0.00 | 0.00 | 0.00 | 0.00 | 0.00 | 0.00 | 0.00 | 0.00 | 0.00 |
| SINTAX50 | 0.00 | 0.00 | 0.00 | 0.00 | 0.00 | 0.00 | 0.00 | 0.00 | 0.00 | 0.00 | 0.00 | 0.00 | 0.00 | 0.00 |
| SINTAX80 | 0.00 | 0.00 | 0.00 | 0.00 | 0.00 | 0.00 | 0.00 | 0.00 | 0.00 | 0.00 | 0.00 | 0.00 | 0.00 | 0.00 |
| SPINGO | 0.00 | 0.00 | 0.00 | 0.00 | 0.00 | 0.00 | 0.00 | 0.00 | 0.00 | 0.00 | 0.00 | 0.00 | 0.00 | 0.00 |
| TOP | 0.00 | 0.00 | 0.00 | 0.00 | 0.00 | 0.00 | 0.00 | 0.00 | 0.00 | 0.00 | 0.00 | 0.00 | 0.00 | 0.00 |

**Table S68. Machine learning metrics computed for the dataset SP RDP ITS 95 at the phylum level.**

| Method | Accuracy | Balanced Accuracy | F1-score Micro | F1-score Macro | F1-score Weighted | Precision Micro | Precision Macro | Precision Weighted | Recall Micro | Recall Macro | Recall Weighted | Jaccard Micro | Jaccard Macro | Jaccard Weighted |
|---|---|---|---|---|---|---|---|---|---|---|---|---|---|---|
| BTOP | 100.00 | 100.00 | 100.00 | 100.00 | 100.00 | 100.00 | 100.00 | 100.00 | 100.00 | 100.00 | 100.00 | 100.00 | 100.00 | 100.00 |
| CT1 | 100.00 | 100.00 | 100.00 | 100.00 | 100.00 | 100.00 | 100.00 | 100.00 | 100.00 | 100.00 | 100.00 | 100.00 | 100.00 | 100.00 |
| CT2 | 100.00 | 100.00 | 100.00 | 100.00 | 100.00 | 100.00 | 100.00 | 100.00 | 100.00 | 100.00 | 100.00 | 100.00 | 100.00 | 100.00 |
| HiTaC | 100.00 | 100.00 | 100.00 | 100.00 | 100.00 | 100.00 | 100.00 | 100.00 | 100.00 | 100.00 | 100.00 | 100.00 | 100.00 | 100.00 |
| HiTaC_Filter | 100.00 | 100.00 | 100.00 | 100.00 | 100.00 | 100.00 | 100.00 | 100.00 | 100.00 | 100.00 | 100.00 | 100.00 | 100.00 | 100.00 |
| KNN | 100.00 | 100.00 | 100.00 | 100.00 | 100.00 | 100.00 | 100.00 | 100.00 | 100.00 | 100.00 | 100.00 | 100.00 | 100.00 | 100.00 |
| KTOP | 100.00 | 100.00 | 100.00 | 100.00 | 100.00 | 100.00 | 100.00 | 100.00 | 100.00 | 100.00 | 100.00 | 100.00 | 100.00 | 100.00 |
| Metaxa2 | 100.00 | 100.00 | 100.00 | 100.00 | 100.00 | 100.00 | 100.00 | 100.00 | 100.00 | 100.00 | 100.00 | 100.00 | 100.00 | 100.00 |
| Microclass | 100.00 | 100.00 | 100.00 | 100.00 | 100.00 | 100.00 | 100.00 | 100.00 | 100.00 | 100.00 | 100.00 | 100.00 | 100.00 | 100.00 |
| NBC50 | 100.00 | 100.00 | 100.00 | 100.00 | 100.00 | 100.00 | 100.00 | 100.00 | 100.00 | 100.00 | 100.00 | 100.00 | 100.00 | 100.00 |
| NBC80 | 100.00 | 100.00 | 100.00 | 100.00 | 100.00 | 100.00 | 100.00 | 100.00 | 100.00 | 100.00 | 100.00 | 100.00 | 100.00 | 100.00 |
| Q2_BLAST | 100.00 | 100.00 | 100.00 | 100.00 | 100.00 | 100.00 | 100.00 | 100.00 | 100.00 | 100.00 | 100.00 | 100.00 | 100.00 | 100.00 |
| Q2_SK | 100.00 | 100.00 | 100.00 | 100.00 | 100.00 | 100.00 | 100.00 | 100.00 | 100.00 | 100.00 | 100.00 | 100.00 | 100.00 | 100.00 |
| Q2_VS | 100.00 | 100.00 | 100.00 | 100.00 | 100.00 | 100.00 | 100.00 | 100.00 | 100.00 | 100.00 | 100.00 | 100.00 | 100.00 | 100.00 |
| RDP50 | 100.00 | 100.00 | 100.00 | 100.00 | 100.00 | 100.00 | 100.00 | 100.00 | 100.00 | 100.00 | 100.00 | 100.00 | 100.00 | 100.00 |
| RDP80 | 100.00 | 100.00 | 100.00 | 100.00 | 100.00 | 100.00 | 100.00 | 100.00 | 100.00 | 100.00 | 100.00 | 100.00 | 100.00 | 100.00 |
| SINTAX50 | 100.00 | 100.00 | 100.00 | 100.00 | 100.00 | 100.00 | 100.00 | 100.00 | 100.00 | 100.00 | 100.00 | 100.00 | 100.00 | 100.00 |
| SINTAX80 | 100.00 | 100.00 | 100.00 | 100.00 | 100.00 | 100.00 | 100.00 | 100.00 | 100.00 | 100.00 | 100.00 | 100.00 | 100.00 | 100.00 |
| TOP | 100.00 | 100.00 | 100.00 | 100.00 | 100.00 | 100.00 | 100.00 | 100.00 | 100.00 | 100.00 | 100.00 | 100.00 | 100.00 | 100.00 |
| Q1 | 99.84 | 99.91 | 99.84 | 74.97 | 99.92 | 99.84 | 75.00 | 100.00 | 99.84 | 74.94 | 99.84 | 99.68 | 74.94 | 99.84 |
| BLCA | 95.89 | 92.69 | 95.89 | 72.07 | 97.87 | 95.89 | 75.00 | 100.00 | 95.89 | 69.51 | 95.89 | 92.10 | 69.51 | 95.89 |
| SPINGO | 0.00 | 0.00 | 0.00 | 0.00 | 0.00 | 0.00 | 0.00 | 0.00 | 0.00 | 0.00 | 0.00 | 0.00 | 0.00 | 0.00 |

Table S69. Machine learning metrics computed for the dataset SP RDP ITS 95 at the class level.

| Method | Accuracy | Balanced Accuracy | F1-score Micro | F1-score Macro | F1-score Weighted | Precision Micro | Precision Macro | Precision Weighted | Recall Micro | Recall Macro | Recall Weighted | Jaccard Micro | Jaccard Macro | Jaccard Weighted |
|---|---|---|---|---|---|---|---|---|---|---|---|---|---|---|
| CT2 | 99.68 | 99.12 | 99.68 | 98.65 | 99.69 | 99.68 | 98.45 | 99.71 | 99.68 | 99.12 | 99.68 | 99.37 | 97.57 | 99.39 |
| HiTaC | 99.68 | 99.12 | 99.68 | 98.65 | 99.69 | 99.68 | 98.45 | 99.71 | 99.68 | 99.12 | 99.68 | 99.37 | 97.57 | 99.39 |
| HiTaC_Filter | 99.68 | 99.12 | 99.68 | 98.65 | 99.69 | 99.68 | 98.45 | 99.71 | 99.68 | 99.12 | 99.68 | 99.37 | 97.57 | 99.39 |
| Q2_BLAST | 99.68 | 99.12 | 99.68 | 98.65 | 99.69 | 99.68 | 98.45 | 99.71 | 99.68 | 99.12 | 99.68 | 99.37 | 97.57 | 99.39 |
| Q2_VS | 99.68 | 99.12 | 99.68 | 98.65 | 99.69 | 99.68 | 98.45 | 99.71 | 99.68 | 99.12 | 99.68 | 99.37 | 97.57 | 99.39 |
| CT1 | 99.60 | 99.00 | 99.60 | 98.58 | 99.61 | 99.60 | 98.42 | 99.63 | 99.60 | 99.00 | 99.60 | 99.21 | 97.42 | 99.23 |
| KTOP | 99.60 | 99.00 | 99.60 | 98.57 | 99.61 | 99.60 | 98.41 | 99.63 | 99.60 | 99.00 | 99.60 | 99.21 | 97.40 | 99.23 |
| TOP | 99.60 | 99.00 | 99.60 | 98.57 | 99.61 | 99.60 | 98.41 | 99.63 | 99.60 | 99.00 | 99.60 | 99.21 | 97.40 | 99.23 |
| BTOP | 99.53 | 98.88 | 99.53 | 98.48 | 99.53 | 99.53 | 98.36 | 99.55 | 99.53 | 98.88 | 99.53 | 99.06 | 97.24 | 99.08 |
| RDP50 | 99.53 | 98.88 | 99.53 | 98.48 | 99.53 | 99.53 | 98.36 | 99.55 | 99.53 | 98.88 | 99.53 | 99.06 | 97.24 | 99.08 |
| SINTAX50 | 99.53 | 98.88 | 99.53 | 93.04 | 99.57 | 99.53 | 92.94 | 99.63 | 99.53 | 93.38 | 99.53 | 99.06 | 91.88 | 99.15 |
| Microclass | 99.45 | 98.76 | 99.45 | 98.40 | 99.45 | 99.45 | 98.31 | 99.47 | 99.45 | 98.76 | 99.45 | 98.90 | 97.07 | 98.92 |
| Q2_SK | 99.45 | 98.76 | 99.45 | 92.98 | 99.52 | 99.45 | 92.94 | 99.63 | 99.45 | 93.27 | 99.45 | 98.90 | 91.77 | 99.07 |
| RDP80 | 99.45 | 98.76 | 99.45 | 92.98 | 99.52 | 99.45 | 92.94 | 99.63 | 99.45 | 93.27 | 99.45 | 98.90 | 91.77 | 99.07 |
| SINTAX80 | 99.45 | 98.76 | 99.45 | 92.98 | 99.52 | 99.45 | 92.94 | 99.63 | 99.45 | 93.27 | 99.45 | 98.90 | 91.77 | 99.07 |
| NBC50 | 99.45 | 98.76 | 99.45 | 92.95 | 99.49 | 99.45 | 92.89 | 99.55 | 99.45 | 93.27 | 99.45 | 98.90 | 91.72 | 99.00 |
| Q1 | 99.37 | 98.79 | 99.37 | 92.97 | 99.45 | 99.37 | 92.89 | 99.55 | 99.37 | 93.30 | 99.37 | 98.74 | 91.75 | 98.92 |
| NBC80 | 99.37 | 98.63 | 99.37 | 92.92 | 99.48 | 99.37 | 92.94 | 99.63 | 99.37 | 93.16 | 99.37 | 98.74 | 91.65 | 99.00 |
| Metaxa2 | 99.37 | 98.63 | 99.37 | 92.89 | 99.44 | 99.37 | 92.89 | 99.55 | 99.37 | 93.16 | 99.37 | 98.74 | 91.60 | 98.92 |
| KNN | 98.74 | 94.95 | 98.74 | 91.00 | 99.17 | 98.74 | 92.52 | 99.69 | 98.74 | 89.68 | 98.74 | 97.50 | 88.68 | 98.46 |
| BLCA | 95.42 | 93.36 | 95.42 | 90.08 | 97.21 | 95.42 | 92.41 | 99.49 | 95.42 | 88.17 | 95.42 | 91.23 | 87.10 | 95.02 |
| SPINGO | 0.00 | 0.00 | 0.00 | 0.00 | 0.00 | 0.00 | 0.00 | 0.00 | 0.00 | 0.00 | 0.00 | 0.00 | 0.00 | 0.00 |

**Table S70. Machine learning metrics computed for the dataset SP RDP ITS 95 at the order level.**

| Method | Accuracy | Balanced Accuracy | F1-score Micro | F1-score Macro | F1-score Weighted | Precision Micro | Precision Macro | Precision Weighted | Recall Micro | Recall Macro | Recall Weighted | Jaccard Micro | Jaccard Macro | Jaccard Weighted |
|---|---|---|---|---|---|---|---|---|---|---|---|---|---|---|
| HiTaC | 99.13 | 97.59 | 99.13 | 95.78 | 99.17 | 99.13 | 96.49 | 99.30 | 99.13 | 95.75 | 99.13 | 98.28 | 94.13 | 98.44 |
| HiTaC_Filter | 99.13 | 97.59 | 99.13 | 94.04 | 99.20 | 99.13 | 94.78 | 99.38 | 99.13 | 93.98 | 99.13 | 98.28 | 92.46 | 98.51 |
| Q2_VS | 99.13 | 96.34 | 99.13 | 94.31 | 99.21 | 99.13 | 94.71 | 99.38 | 99.13 | 94.52 | 99.13 | 98.28 | 93.00 | 98.59 |
| Q2_BLAST | 99.13 | 96.27 | 99.13 | 94.24 | 99.17 | 99.13 | 94.64 | 99.30 | 99.13 | 94.45 | 99.13 | 98.28 | 92.86 | 98.52 |
| KTOP | 99.05 | 97.20 | 99.05 | 95.80 | 99.06 | 99.05 | 96.93 | 99.19 | 99.05 | 95.36 | 99.05 | 98.12 | 94.19 | 98.25 |
| TOP | 99.05 | 97.20 | 99.05 | 95.80 | 99.06 | 99.05 | 96.93 | 99.19 | 99.05 | 95.36 | 99.05 | 98.12 | 94.19 | 98.25 |
| CT2 | 99.05 | 95.35 | 99.05 | 93.64 | 99.12 | 99.05 | 94.67 | 99.30 | 99.05 | 93.55 | 99.05 | 98.12 | 91.99 | 98.44 |
| SINTAX50 | 98.97 | 97.12 | 98.97 | 94.02 | 99.10 | 98.97 | 95.20 | 99.34 | 98.97 | 93.52 | 98.97 | 97.97 | 92.43 | 98.32 |
| RDP50 | 98.97 | 97.00 | 98.97 | 95.49 | 98.97 | 98.97 | 96.62 | 99.13 | 98.97 | 95.17 | 98.97 | 97.97 | 93.68 | 98.11 |
| CT1 | 98.89 | 97.21 | 98.89 | 95.00 | 98.92 | 98.89 | 95.66 | 99.08 | 98.89 | 95.38 | 98.89 | 97.81 | 92.93 | 97.99 |
| BTOP | 98.89 | 96.93 | 98.89 | 95.44 | 98.89 | 98.89 | 96.58 | 99.05 | 98.89 | 95.10 | 98.89 | 97.81 | 93.58 | 97.96 |
| Microclass | 98.89 | 96.93 | 98.89 | 95.31 | 98.89 | 98.89 | 96.40 | 99.09 | 98.89 | 95.10 | 98.89 | 97.81 | 93.38 | 97.98 |
| RDP80 | 98.89 | 96.93 | 98.89 | 93.87 | 99.00 | 98.89 | 95.17 | 99.27 | 98.89 | 93.34 | 98.89 | 97.81 | 92.21 | 98.17 |
| NBC50 | 98.89 | 96.93 | 98.89 | 93.68 | 98.92 | 98.89 | 94.83 | 99.13 | 98.89 | 93.34 | 98.89 | 97.81 | 91.87 | 98.03 |
| Q2_SK | 98.81 | 96.85 | 98.81 | 93.81 | 98.92 | 98.81 | 95.14 | 99.20 | 98.81 | 93.26 | 98.81 | 97.66 | 92.11 | 98.02 |
| Metaxa2 | 98.74 | 96.77 | 98.74 | 93.50 | 98.85 | 98.74 | 94.68 | 99.16 | 98.74 | 93.19 | 98.74 | 97.50 | 91.58 | 97.90 |
| Q1 | 98.66 | 96.87 | 98.66 | 93.35 | 98.74 | 98.66 | 94.30 | 99.02 | 98.66 | 93.28 | 98.66 | 97.35 | 91.29 | 97.68 |
| NBC80 | 98.66 | 96.74 | 98.66 | 93.79 | 98.91 | 98.66 | 95.20 | 99.34 | 98.66 | 93.16 | 98.66 | 97.35 | 92.07 | 98.00 |
| SINTAX80 | 98.42 | 96.47 | 98.42 | 93.72 | 98.90 | 98.42 | 95.34 | 99.57 | 98.42 | 92.89 | 98.42 | 96.89 | 91.94 | 97.99 |
| BLCA | 94.78 | 90.84 | 94.78 | 88.81 | 96.50 | 94.78 | 91.17 | 98.98 | 94.78 | 87.47 | 94.78 | 90.08 | 86.08 | 94.10 |
| KNN | 94.78 | 75.55 | 94.78 | 76.32 | 96.07 | 94.78 | 82.04 | 98.38 | 94.78 | 74.13 | 94.78 | 90.08 | 73.15 | 94.36 |
| SPINGO | 0.00 | 0.00 | 0.00 | 0.00 | 0.00 | 0.00 | 0.00 | 0.00 | 0.00 | 0.00 | 0.00 | 0.00 | 0.00 | 0.00 |

**Table S71. Machine learning metrics computed for the dataset SP RDP ITS 95 at the family level.**

| Method | Accuracy | Balanced Accuracy | F1-score Micro | F1-score Macro | F1-score Weighted | Precision Micro | Precision Macro | Precision Weighted | Recall Micro | Recall Macro | Recall Weighted | Jaccard Micro | Jaccard Macro | Jaccard Weighted |
|---|---|---|---|---|---|---|---|---|---|---|---|---|---|---|
| HiTaC | 98.02 | 96.05 | 98.02 | 94.06 | 98.03 | 98.02 | 94.26 | 98.37 | 98.02 | 94.76 | 98.02 | 96.12 | 92.48 | 96.74 |
| TOP | 97.79 | 96.39 | 97.79 | 93.45 | 97.91 | 97.79 | 93.92 | 98.44 | 97.79 | 93.84 | 97.79 | 95.67 | 91.89 | 96.56 |
| BTOP | 97.79 | 96.33 | 97.79 | 93.35 | 97.89 | 97.79 | 94.00 | 98.56 | 97.79 | 93.78 | 97.79 | 95.67 | 91.75 | 96.52 |
| KTOP | 97.79 | 96.10 | 97.79 | 92.59 | 97.92 | 97.79 | 93.32 | 98.57 | 97.79 | 92.94 | 97.79 | 95.67 | 90.92 | 96.57 |
| RDP50 | 97.63 | 95.81 | 97.63 | 92.73 | 97.64 | 97.63 | 93.22 | 98.21 | 97.63 | 93.27 | 97.63 | 95.37 | 91.17 | 96.28 |
| Microclass | 97.55 | 95.82 | 97.55 | 92.86 | 97.67 | 97.55 | 93.45 | 98.26 | 97.55 | 93.28 | 97.55 | 95.22 | 91.02 | 96.21 |
| HiTaC_Filter | 97.55 | 94.83 | 97.55 | 93.82 | 98.05 | 97.55 | 94.81 | 98.94 | 97.55 | 93.56 | 97.55 | 95.22 | 92.40 | 96.81 |
| SINTAX50 | 97.31 | 95.75 | 97.31 | 93.28 | 97.74 | 97.31 | 94.44 | 98.74 | 97.31 | 93.21 | 97.31 | 94.77 | 91.68 | 96.28 |
| NBC50 | 97.31 | 95.36 | 97.31 | 91.89 | 97.50 | 97.31 | 92.63 | 98.26 | 97.31 | 92.23 | 97.31 | 94.77 | 90.18 | 96.03 |
| Q2_SK | 97.23 | 95.26 | 97.23 | 92.74 | 97.55 | 97.23 | 93.75 | 98.44 | 97.23 | 92.74 | 97.23 | 94.62 | 91.19 | 96.15 |
| Q1 | 97.15 | 94.13 | 97.15 | 91.42 | 97.25 | 97.15 | 92.48 | 97.98 | 97.15 | 91.64 | 97.15 | 94.47 | 89.47 | 95.51 |
| CT1 | 97.00 | 92.06 | 97.00 | 89.15 | 96.81 | 97.00 | 89.34 | 97.17 | 97.00 | 90.22 | 97.00 | 94.17 | 86.98 | 95.06 |
| SPINGO | 96.84 | 95.05 | 96.84 | 92.54 | 97.44 | 96.84 | 93.63 | 98.66 | 96.84 | 92.53 | 96.84 | 93.87 | 90.86 | 95.96 |
| RDP80 | 96.84 | 94.92 | 96.84 | 93.07 | 97.43 | 96.84 | 94.23 | 98.65 | 96.84 | 93.03 | 96.84 | 93.87 | 91.36 | 95.97 |
| NBC80 | 96.60 | 94.80 | 96.60 | 93.06 | 97.32 | 96.60 | 94.32 | 98.69 | 96.60 | 92.91 | 96.60 | 93.43 | 91.28 | 95.74 |
| Metaxa2 | 96.28 | 91.68 | 96.28 | 89.61 | 96.69 | 96.28 | 90.60 | 97.87 | 96.28 | 89.84 | 96.28 | 92.84 | 87.79 | 94.98 |
| SINTAX80 | 96.05 | 93.22 | 96.05 | 92.44 | 97.02 | 96.05 | 93.97 | 98.66 | 96.05 | 91.97 | 96.05 | 92.40 | 90.69 | 95.39 |
| Q2_VS | 95.73 | 87.19 | 95.73 | 85.93 | 95.95 | 95.73 | 86.73 | 96.83 | 95.73 | 86.60 | 95.73 | 91.81 | 83.74 | 94.17 |
| CT2 | 95.65 | 84.34 | 95.65 | 82.79 | 95.80 | 95.65 | 83.82 | 96.60 | 95.65 | 83.20 | 95.65 | 91.67 | 80.74 | 94.05 |
| Q2_BLAST | 95.49 | 86.87 | 95.49 | 85.42 | 95.76 | 95.49 | 86.44 | 96.73 | 95.49 | 86.28 | 95.49 | 91.38 | 83.05 | 93.90 |
| BLCA | 92.96 | 90.80 | 92.96 | 88.47 | 94.77 | 92.96 | 90.51 | 97.92 | 92.96 | 87.81 | 92.96 | 86.85 | 86.27 | 91.80 |
| KNN | 85.14 | 59.25 | 85.14 | 61.04 | 87.49 | 85.14 | 65.84 | 91.92 | 85.14 | 58.85 | 85.14 | 74.12 | 58.15 | 84.56 |

**Table S72. Machine learning metrics computed for the dataset SP RDP ITS 95 at the genus level.**

| Method | Accuracy | Balanced Accuracy | F1-score Micro | F1-score Macro | F1-score Weighted | Precision Micro | Precision Macro | Precision Weighted | Recall Micro | Recall Macro | Recall Weighted | Jaccard Micro | Jaccard Macro | Jaccard Weighted |
|---|---|---|---|---|---|---|---|---|---|---|---|---|---|---|
| HiTaC | 83.64 | 76.31 | 83.64 | 66.46 | 82.44 | 83.64 | 66.64 | 83.47 | 83.64 | 68.60 | 83.64 | 71.88 | 64.26 | 78.77 |
| Microclass | 81.98 | 77.03 | 81.98 | 65.65 | 82.00 | 81.98 | 66.54 | 84.57 | 81.98 | 67.33 | 81.98 | 69.46 | 63.35 | 77.82 |
| TOP | 81.90 | 77.14 | 81.90 | 66.36 | 82.01 | 81.90 | 67.22 | 84.51 | 81.90 | 67.77 | 81.90 | 69.34 | 64.26 | 78.03 |
| BTOP | 81.34 | 76.49 | 81.34 | 65.20 | 81.90 | 81.34 | 66.72 | 85.37 | 81.34 | 66.52 | 81.34 | 68.55 | 62.74 | 77.37 |
| NBC50 | 81.11 | 76.34 | 81.11 | 66.84 | 81.36 | 81.11 | 67.76 | 84.16 | 81.11 | 68.27 | 81.11 | 68.22 | 64.62 | 77.33 |
| RDP50 | 80.87 | 75.77 | 80.87 | 66.39 | 81.24 | 80.87 | 67.35 | 84.26 | 80.87 | 67.94 | 80.87 | 67.88 | 64.12 | 77.18 |
| KTOP | 80.79 | 75.66 | 80.79 | 64.24 | 81.04 | 80.79 | 65.42 | 84.18 | 80.79 | 65.80 | 80.79 | 67.77 | 61.95 | 76.96 |
| Q1 | 80.71 | 72.72 | 80.71 | 61.90 | 80.29 | 80.71 | 62.53 | 82.14 | 80.71 | 63.40 | 80.71 | 67.66 | 59.61 | 76.35 |
| Q2_SK | 79.92 | 75.06 | 79.92 | 66.21 | 80.74 | 79.92 | 67.46 | 84.49 | 79.92 | 67.30 | 79.92 | 66.56 | 64.10 | 76.69 |
| CT1 | 79.68 | 71.08 | 79.68 | 60.03 | 79.56 | 79.68 | 60.70 | 81.82 | 79.68 | 61.67 | 79.68 | 66.23 | 57.79 | 75.70 |
| SINTAX50 | 79.60 | 73.91 | 79.60 | 66.19 | 80.28 | 79.60 | 67.59 | 83.46 | 79.60 | 67.14 | 79.60 | 66.12 | 63.93 | 76.43 |
| HiTaC_Filter | 79.60 | 71.69 | 79.60 | 66.33 | 79.95 | 79.60 | 67.35 | 82.59 | 79.60 | 67.23 | 79.60 | 66.12 | 64.22 | 76.28 |
| SPINGO | 77.00 | 71.48 | 77.00 | 64.78 | 78.61 | 77.00 | 66.65 | 83.22 | 77.00 | 65.10 | 77.00 | 62.60 | 62.49 | 74.39 |
| RDP80 | 76.60 | 72.35 | 76.60 | 66.35 | 78.14 | 76.60 | 68.40 | 82.46 | 76.60 | 66.59 | 76.60 | 62.08 | 64.01 | 73.95 |
| NBC80 | 76.05 | 71.77 | 76.05 | 65.86 | 77.78 | 76.05 | 67.98 | 82.32 | 76.05 | 66.06 | 76.05 | 61.35 | 63.48 | 73.54 |
| Metaxa2 | 75.81 | 68.64 | 75.81 | 61.53 | 77.38 | 75.81 | 63.13 | 81.97 | 75.81 | 62.35 | 75.81 | 61.04 | 59.23 | 72.80 |
| BLCA | 75.18 | 70.18 | 75.18 | 62.12 | 76.89 | 75.18 | 63.98 | 81.71 | 75.18 | 62.60 | 75.18 | 60.23 | 59.90 | 72.56 |
| Q2_VS | 72.65 | 57.03 | 72.65 | 52.13 | 72.15 | 72.65 | 53.08 | 74.63 | 72.65 | 53.78 | 72.65 | 57.05 | 49.89 | 68.29 |
| Q2_BLAST | 72.41 | 55.99 | 72.41 | 50.68 | 71.64 | 72.41 | 50.81 | 72.62 | 72.41 | 52.51 | 72.41 | 56.75 | 48.54 | 67.81 |
| CT2 | 71.15 | 51.83 | 71.15 | 47.39 | 70.43 | 71.15 | 47.66 | 72.26 | 71.15 | 49.14 | 71.15 | 55.21 | 45.39 | 66.76 |
| SINTAX80 | 70.91 | 64.77 | 70.91 | 62.10 | 73.62 | 70.91 | 64.97 | 79.61 | 70.91 | 61.41 | 70.91 | 54.93 | 59.73 | 69.22 |
| KNN | 46.01 | 24.46 | 46.01 | 25.16 | 49.58 | 46.01 | 28.20 | 58.94 | 46.01 | 24.25 | 46.01 | 29.88 | 23.56 | 45.31 |

**Table S73. Machine learning metrics computed for the dataset SP RDP ITS 95 at the species level.**

| Method | Accuracy | Balanced Accuracy | F1-score Micro | F1-score Macro | F1-score Weighted | Precision Micro | Precision Macro | Precision Weighted | Recall Micro | Recall Macro | Recall Weighted | Jaccard Micro | Jaccard Macro | Jaccard Weighted |
|---|---|---|---|---|---|---|---|---|---|---|---|---|---|---|
| BTOP | 0.32 | 0.38 | 0.32 | 0.18 | 0.26 | 0.32 | 0.16 | 0.24 | 0.32 | 0.21 | 0.32 | 0.16 | 0.16 | 0.22 |
| KTOP | 0.24 | 0.32 | 0.24 | 0.16 | 0.21 | 0.24 | 0.15 | 0.20 | 0.24 | 0.18 | 0.24 | 0.12 | 0.15 | 0.20 |
| HiTaC | 0.24 | 0.32 | 0.24 | 0.14 | 0.18 | 0.24 | 0.13 | 0.17 | 0.24 | 0.18 | 0.24 | 0.12 | 0.13 | 0.17 |
| Microclass | 0.24 | 0.32 | 0.24 | 0.14 | 0.18 | 0.24 | 0.13 | 0.17 | 0.24 | 0.18 | 0.24 | 0.12 | 0.13 | 0.17 |
| HiTaC_Filter | 0.16 | 0.22 | 0.16 | 0.18 | 0.16 | 0.16 | 0.18 | 0.16 | 0.16 | 0.18 | 0.16 | 0.08 | 0.18 | 0.16 |
| SINTAX50 | 0.16 | 0.22 | 0.16 | 0.15 | 0.16 | 0.16 | 0.15 | 0.16 | 0.16 | 0.15 | 0.16 | 0.08 | 0.15 | 0.16 |
| Q2_SK | 0.16 | 0.22 | 0.16 | 0.14 | 0.16 | 0.16 | 0.14 | 0.16 | 0.16 | 0.14 | 0.16 | 0.08 | 0.14 | 0.16 |
| TOP | 0.16 | 0.22 | 0.16 | 0.12 | 0.16 | 0.16 | 0.12 | 0.16 | 0.16 | 0.12 | 0.16 | 0.08 | 0.12 | 0.16 |
| NBC50 | 0.16 | 0.22 | 0.16 | 0.11 | 0.13 | 0.16 | 0.10 | 0.12 | 0.16 | 0.13 | 0.16 | 0.08 | 0.10 | 0.12 |
| RDP50 | 0.16 | 0.22 | 0.16 | 0.11 | 0.13 | 0.16 | 0.10 | 0.12 | 0.16 | 0.13 | 0.16 | 0.08 | 0.10 | 0.12 |
| Q2_BLAST | 0.08 | 0.11 | 0.08 | 0.10 | 0.08 | 0.08 | 0.10 | 0.08 | 0.08 | 0.10 | 0.08 | 0.04 | 0.10 | 0.08 |
| Q2_VS | 0.08 | 0.11 | 0.08 | 0.10 | 0.08 | 0.08 | 0.10 | 0.08 | 0.08 | 0.10 | 0.08 | 0.04 | 0.10 | 0.08 |
| SINTAX80 | 0.08 | 0.11 | 0.08 | 0.09 | 0.08 | 0.08 | 0.09 | 0.08 | 0.08 | 0.09 | 0.08 | 0.04 | 0.09 | 0.08 |
| NBC80 | 0.08 | 0.11 | 0.08 | 0.08 | 0.08 | 0.08 | 0.08 | 0.08 | 0.08 | 0.08 | 0.08 | 0.04 | 0.08 | 0.08 |
| RDP80 | 0.08 | 0.11 | 0.08 | 0.08 | 0.08 | 0.08 | 0.08 | 0.08 | 0.08 | 0.08 | 0.08 | 0.04 | 0.08 | 0.08 |
| SPINGO | 0.08 | 0.11 | 0.08 | 0.08 | 0.08 | 0.08 | 0.08 | 0.08 | 0.08 | 0.08 | 0.08 | 0.04 | 0.08 | 0.08 |
| BLCA | 0.08 | 0.11 | 0.08 | 0.07 | 0.08 | 0.08 | 0.07 | 0.08 | 0.08 | 0.07 | 0.08 | 0.04 | 0.07 | 0.08 |
| Q1 | 0.08 | 0.11 | 0.08 | 0.07 | 0.08 | 0.08 | 0.07 | 0.08 | 0.08 | 0.07 | 0.08 | 0.04 | 0.07 | 0.08 |
| CT1 | 0.00 | 0.00 | 0.00 | 0.00 | 0.00 | 0.00 | 0.00 | 0.00 | 0.00 | 0.00 | 0.00 | 0.00 | 0.00 | 0.00 |
| CT2 | 0.00 | 0.00 | 0.00 | 0.00 | 0.00 | 0.00 | 0.00 | 0.00 | 0.00 | 0.00 | 0.00 | 0.00 | 0.00 | 0.00 |
| KNN | 0.00 | 0.00 | 0.00 | 0.00 | 0.00 | 0.00 | 0.00 | 0.00 | 0.00 | 0.00 | 0.00 | 0.00 | 0.00 | 0.00 |
| Metaxa2 | 0.00 | 0.00 | 0.00 | 0.00 | 0.00 | 0.00 | 0.00 | 0.00 | 0.00 | 0.00 | 0.00 | 0.00 | 0.00 | 0.00 |

**Table S74. Machine learning metrics computed for the dataset SP RDP ITS 97 at the phylum level.**

| Method | Accuracy | Balanced Accuracy | F1-score Micro | F1-score Macro | F1-score Weighted | Precision Micro | Precision Macro | Precision Weighted | Recall Micro | Recall Macro | Recall Weighted | Jaccard Micro | Jaccard Macro | Jaccard Weighted |
|---|---|---|---|---|---|---|---|---|---|---|---|---|---|---|
| BTOP | 100.00 | 100.00 | 100.00 | 100.00 | 100.00 | 100.00 | 100.00 | 100.00 | 100.00 | 100.00 | 100.00 | 100.00 | 100.00 | 100.00 |
| CT1 | 100.00 | 100.00 | 100.00 | 100.00 | 100.00 | 100.00 | 100.00 | 100.00 | 100.00 | 100.00 | 100.00 | 100.00 | 100.00 | 100.00 |
| CT2 | 100.00 | 100.00 | 100.00 | 100.00 | 100.00 | 100.00 | 100.00 | 100.00 | 100.00 | 100.00 | 100.00 | 100.00 | 100.00 | 100.00 |
| HiTaC | 100.00 | 100.00 | 100.00 | 100.00 | 100.00 | 100.00 | 100.00 | 100.00 | 100.00 | 100.00 | 100.00 | 100.00 | 100.00 | 100.00 |
| HiTaC_Filter | 100.00 | 100.00 | 100.00 | 100.00 | 100.00 | 100.00 | 100.00 | 100.00 | 100.00 | 100.00 | 100.00 | 100.00 | 100.00 | 100.00 |
| KTOP | 100.00 | 100.00 | 100.00 | 100.00 | 100.00 | 100.00 | 100.00 | 100.00 | 100.00 | 100.00 | 100.00 | 100.00 | 100.00 | 100.00 |
| Microclass | 100.00 | 100.00 | 100.00 | 100.00 | 100.00 | 100.00 | 100.00 | 100.00 | 100.00 | 100.00 | 100.00 | 100.00 | 100.00 | 100.00 |
| NBC50 | 100.00 | 100.00 | 100.00 | 100.00 | 100.00 | 100.00 | 100.00 | 100.00 | 100.00 | 100.00 | 100.00 | 100.00 | 100.00 | 100.00 |
| NBC80 | 100.00 | 100.00 | 100.00 | 100.00 | 100.00 | 100.00 | 100.00 | 100.00 | 100.00 | 100.00 | 100.00 | 100.00 | 100.00 | 100.00 |
| Q2_BLAST | 100.00 | 100.00 | 100.00 | 100.00 | 100.00 | 100.00 | 100.00 | 100.00 | 100.00 | 100.00 | 100.00 | 100.00 | 100.00 | 100.00 |
| Q2_SK | 100.00 | 100.00 | 100.00 | 100.00 | 100.00 | 100.00 | 100.00 | 100.00 | 100.00 | 100.00 | 100.00 | 100.00 | 100.00 | 100.00 |
| RDP50 | 100.00 | 100.00 | 100.00 | 100.00 | 100.00 | 100.00 | 100.00 | 100.00 | 100.00 | 100.00 | 100.00 | 100.00 | 100.00 | 100.00 |
| RDP80 | 100.00 | 100.00 | 100.00 | 100.00 | 100.00 | 100.00 | 100.00 | 100.00 | 100.00 | 100.00 | 100.00 | 100.00 | 100.00 | 100.00 |
| SINTAX50 | 100.00 | 100.00 | 100.00 | 100.00 | 100.00 | 100.00 | 100.00 | 100.00 | 100.00 | 100.00 | 100.00 | 100.00 | 100.00 | 100.00 |
| SINTAX80 | 100.00 | 100.00 | 100.00 | 100.00 | 100.00 | 100.00 | 100.00 | 100.00 | 100.00 | 100.00 | 100.00 | 100.00 | 100.00 | 100.00 |
| TOP | 100.00 | 100.00 | 100.00 | 100.00 | 100.00 | 100.00 | 100.00 | 100.00 | 100.00 | 100.00 | 100.00 | 100.00 | 100.00 | 100.00 |
| Q2_VS | 99.93 | 99.98 | 99.93 | 85.71 | 99.96 | 99.93 | 85.71 | 100.00 | 99.93 | 85.70 | 99.93 | 99.86 | 85.70 | 99.93 |
| Q1 | 99.93 | 99.96 | 99.93 | 85.70 | 99.96 | 99.93 | 85.71 | 100.00 | 99.93 | 85.68 | 99.93 | 99.86 | 85.68 | 99.93 |
| KNN | 99.86 | 66.67 | 99.86 | 57.14 | 99.86 | 99.86 | 57.14 | 99.86 | 99.86 | 57.14 | 99.86 | 99.72 | 57.14 | 99.86 |
| Metaxa2 | 99.86 | 66.67 | 99.86 | 57.14 | 99.86 | 99.86 | 57.14 | 99.86 | 99.86 | 57.14 | 99.86 | 99.72 | 57.14 | 99.86 |
| BLCA | 94.51 | 53.31 | 94.51 | 50.01 | 97.07 | 94.51 | 57.14 | 99.86 | 94.51 | 45.69 | 94.51 | 89.59 | 45.69 | 94.51 |
| SPINGO | 0.00 | 0.00 | 0.00 | 0.00 | 0.00 | 0.00 | 0.00 | 0.00 | 0.00 | 0.00 | 0.00 | 0.00 | 0.00 | 0.00 |

**Table S75. Machine learning metrics computed for the dataset SP RDP ITS 97 at the class level.**

| Method | Accuracy | Balanced Accuracy | F1-score Micro | F1-score Macro | F1-score Weighted | Precision Micro | Precision Macro | Precision Weighted | Recall Micro | Recall Macro | Recall Weighted | Jaccard Micro | Jaccard Macro | Jaccard Weighted |
|---|---|---|---|---|---|---|---|---|---|---|---|---|---|---|
| BTOP | 99.93 | 99.96 | 99.93 | 99.94 | 99.93 | 99.93 | 99.92 | 99.93 | 99.93 | 99.96 | 99.93 | 99.86 | 99.88 | 99.86 |
| CT2 | 99.93 | 99.96 | 99.93 | 99.94 | 99.93 | 99.93 | 99.92 | 99.93 | 99.93 | 99.96 | 99.93 | 99.86 | 99.88 | 99.86 |
| HiTaC | 99.93 | 99.96 | 99.93 | 99.94 | 99.93 | 99.93 | 99.92 | 99.93 | 99.93 | 99.96 | 99.93 | 99.86 | 99.88 | 99.86 |
| HiTaC_Filter | 99.93 | 99.96 | 99.93 | 99.94 | 99.93 | 99.93 | 99.92 | 99.93 | 99.93 | 99.96 | 99.93 | 99.86 | 99.88 | 99.86 |
| KTOP | 99.93 | 99.96 | 99.93 | 99.94 | 99.93 | 99.93 | 99.92 | 99.93 | 99.93 | 99.96 | 99.93 | 99.86 | 99.88 | 99.86 |
| Microclass | 99.93 | 99.96 | 99.93 | 99.94 | 99.93 | 99.93 | 99.92 | 99.93 | 99.93 | 99.96 | 99.93 | 99.86 | 99.88 | 99.86 |
| NBC50 | 99.93 | 99.96 | 99.93 | 99.94 | 99.93 | 99.93 | 99.92 | 99.93 | 99.93 | 99.96 | 99.93 | 99.86 | 99.88 | 99.86 |
| NBC80 | 99.93 | 99.96 | 99.93 | 99.94 | 99.93 | 99.93 | 99.92 | 99.93 | 99.93 | 99.96 | 99.93 | 99.86 | 99.88 | 99.86 |
| Q2_SK | 99.93 | 99.96 | 99.93 | 99.94 | 99.93 | 99.93 | 99.92 | 99.93 | 99.93 | 99.96 | 99.93 | 99.86 | 99.88 | 99.86 |
| RDP50 | 99.93 | 99.96 | 99.93 | 99.94 | 99.93 | 99.93 | 99.92 | 99.93 | 99.93 | 99.96 | 99.93 | 99.86 | 99.88 | 99.86 |
| RDP80 | 99.93 | 99.96 | 99.93 | 99.94 | 99.93 | 99.93 | 99.92 | 99.93 | 99.93 | 99.96 | 99.93 | 99.86 | 99.88 | 99.86 |
| SINTAX50 | 99.93 | 99.96 | 99.93 | 99.94 | 99.93 | 99.93 | 99.92 | 99.93 | 99.93 | 99.96 | 99.93 | 99.86 | 99.88 | 99.86 |
| TOP | 99.93 | 99.96 | 99.93 | 99.94 | 99.93 | 99.93 | 99.92 | 99.93 | 99.93 | 99.96 | 99.93 | 99.86 | 99.88 | 99.86 |
| Q2_BLAST | 99.86 | 99.88 | 99.86 | 99.88 | 99.86 | 99.86 | 99.89 | 99.86 | 99.86 | 99.88 | 99.86 | 99.72 | 99.77 | 99.72 |
| CT1 | 99.86 | 99.88 | 99.86 | 94.90 | 99.89 | 99.86 | 94.92 | 99.93 | 99.86 | 94.88 | 99.86 | 99.72 | 94.81 | 99.79 |
| Q2_VS | 99.86 | 99.88 | 99.86 | 94.90 | 99.89 | 99.86 | 94.92 | 99.93 | 99.86 | 94.88 | 99.86 | 99.72 | 94.81 | 99.79 |
| SINTAX80 | 99.86 | 99.88 | 99.86 | 94.90 | 99.89 | 99.86 | 94.92 | 99.93 | 99.86 | 94.88 | 99.86 | 99.72 | 94.81 | 99.79 |
| Q1 | 99.79 | 99.86 | 99.79 | 94.90 | 99.86 | 99.79 | 94.92 | 99.93 | 99.79 | 94.87 | 99.79 | 99.58 | 94.79 | 99.72 |
| Metaxa2 | 99.72 | 89.35 | 99.72 | 84.90 | 99.75 | 99.72 | 84.92 | 99.79 | 99.72 | 84.88 | 99.72 | 99.44 | 84.81 | 99.65 |
| KNN | 98.10 | 82.41 | 98.10 | 80.98 | 98.88 | 98.10 | 84.91 | 99.78 | 98.10 | 78.29 | 98.10 | 96.27 | 78.22 | 98.04 |
| BLCA | 94.44 | 81.54 | 94.44 | 80.43 | 96.73 | 94.44 | 84.92 | 99.79 | 94.44 | 77.46 | 94.44 | 89.46 | 77.39 | 94.37 |
| SPINGO | 0.00 | 0.00 | 0.00 | 0.00 | 0.00 | 0.00 | 0.00 | 0.00 | 0.00 | 0.00 | 0.00 | 0.00 | 0.00 | 0.00 |

**Table S76. Machine learning metrics computed for the dataset SP RDP ITS 97 at the order level.**

| Method | Accuracy | Balanced Accuracy | F1-score Micro | F1-score Macro | F1-score Weighted | Precision Micro | Precision Macro | Precision Weighted | Recall Micro | Recall Macro | Recall Weighted | Jaccard Micro | Jaccard Macro | Jaccard Weighted |
|---|---|---|---|---|---|---|---|---|---|---|---|---|---|---|
| BTOP | 99.44 | 98.34 | 99.44 | 97.67 | 99.51 | 99.44 | 98.25 | 99.69 | 99.44 | 98.34 | 99.44 | 98.88 | 96.59 | 99.12 |
| KTOP | 99.44 | 98.34 | 99.44 | 97.67 | 99.51 | 99.44 | 98.25 | 99.69 | 99.44 | 98.34 | 99.44 | 98.88 | 96.59 | 99.12 |
| Microclass | 99.44 | 98.34 | 99.44 | 97.67 | 99.51 | 99.44 | 98.25 | 99.69 | 99.44 | 98.34 | 99.44 | 98.88 | 96.59 | 99.12 |
| NBC50 | 99.44 | 98.34 | 99.44 | 97.67 | 99.51 | 99.44 | 98.25 | 99.69 | 99.44 | 98.34 | 99.44 | 98.88 | 96.59 | 99.12 |
| NBC80 | 99.44 | 98.34 | 99.44 | 97.67 | 99.51 | 99.44 | 98.25 | 99.69 | 99.44 | 98.34 | 99.44 | 98.88 | 96.59 | 99.12 |
| Q2_SK | 99.44 | 98.34 | 99.44 | 97.67 | 99.51 | 99.44 | 98.25 | 99.69 | 99.44 | 98.34 | 99.44 | 98.88 | 96.59 | 99.12 |
| RDP50 | 99.44 | 98.34 | 99.44 | 97.67 | 99.51 | 99.44 | 98.25 | 99.69 | 99.44 | 98.34 | 99.44 | 98.88 | 96.59 | 99.12 |
| RDP80 | 99.44 | 98.34 | 99.44 | 97.67 | 99.51 | 99.44 | 98.25 | 99.69 | 99.44 | 98.34 | 99.44 | 98.88 | 96.59 | 99.12 |
| TOP | 99.44 | 98.34 | 99.44 | 97.67 | 99.51 | 99.44 | 98.25 | 99.69 | 99.44 | 98.34 | 99.44 | 98.88 | 96.59 | 99.12 |
| SINTAX50 | 99.37 | 98.27 | 99.37 | 96.11 | 99.48 | 99.37 | 96.71 | 99.69 | 99.37 | 96.74 | 99.37 | 98.74 | 95.01 | 99.05 |
| HiTaC | 99.37 | 98.14 | 99.37 | 97.34 | 99.44 | 99.37 | 97.85 | 99.63 | 99.37 | 98.14 | 99.37 | 98.74 | 95.99 | 99.00 |
| HiTaC_Filter | 99.37 | 98.14 | 99.37 | 96.04 | 99.47 | 99.37 | 96.71 | 99.69 | 99.37 | 96.61 | 99.37 | 98.74 | 94.88 | 99.05 |
| SINTAX80 | 99.30 | 98.16 | 99.30 | 96.08 | 99.47 | 99.30 | 96.75 | 99.75 | 99.30 | 96.63 | 99.30 | 98.60 | 94.94 | 99.05 |
| Q1 | 99.30 | 98.14 | 99.30 | 95.84 | 99.44 | 99.30 | 96.36 | 99.70 | 99.30 | 96.60 | 99.30 | 98.60 | 94.53 | 99.00 |
| Q2_VS | 99.30 | 96.71 | 99.30 | 94.37 | 99.44 | 99.30 | 94.80 | 99.70 | 99.30 | 95.20 | 99.30 | 98.60 | 93.13 | 99.07 |
| Q2_BLAST | 99.30 | 96.71 | 99.30 | 94.36 | 99.41 | 99.30 | 94.79 | 99.63 | 99.30 | 95.20 | 99.30 | 98.60 | 93.11 | 99.00 |
| CT1 | 99.23 | 97.98 | 99.23 | 95.69 | 99.34 | 99.23 | 96.23 | 99.56 | 99.23 | 96.45 | 99.23 | 98.46 | 94.24 | 98.79 |
| CT2 | 99.23 | 96.62 | 99.23 | 94.25 | 99.34 | 99.23 | 94.68 | 99.56 | 99.23 | 95.11 | 99.23 | 98.46 | 92.91 | 98.86 |
| Metaxa2 | 99.01 | 94.72 | 99.01 | 92.81 | 99.26 | 99.01 | 93.62 | 99.61 | 99.01 | 93.24 | 99.01 | 98.05 | 91.55 | 98.77 |
| KNN | 95.85 | 78.05 | 95.85 | 76.82 | 97.09 | 95.85 | 78.00 | 98.55 | 95.85 | 76.83 | 95.85 | 92.02 | 75.15 | 95.67 |
| BLCA | 93.87 | 90.37 | 93.87 | 89.59 | 96.14 | 93.87 | 92.02 | 99.48 | 93.87 | 88.96 | 93.87 | 88.45 | 87.23 | 93.56 |
| SPINGO | 0.00 | 0.00 | 0.00 | 0.00 | 0.00 | 0.00 | 0.00 | 0.00 | 0.00 | 0.00 | 0.00 | 0.00 | 0.00 | 0.00 |

**Table S77. Machine learning metrics computed for the dataset SP RDP ITS 97 at the family level.**

| Method | Accuracy | Balanced Accuracy | F1-score Micro | F1-score Macro | F1-score Weighted | Precision Micro | Precision Macro | Precision Weighted | Recall Micro | Recall Macro | Recall Weighted | Jaccard Micro | Jaccard Macro | Jaccard Weighted |
|---|---|---|---|---|---|---|---|---|---|---|---|---|---|---|
| TOP | 99.23 | 98.28 | 99.23 | 97.07 | 99.26 | 99.23 | 97.40 | 99.55 | 99.23 | 97.65 | 99.23 | 98.46 | 96.33 | 98.85 |
| BTOP | 99.16 | 97.95 | 99.16 | 96.34 | 99.21 | 99.16 | 96.78 | 99.52 | 99.16 | 96.71 | 99.16 | 98.33 | 95.61 | 98.79 |
| KTOP | 99.15 | 98.25 | 99.15 | 97.02 | 99.19 | 99.15 | 97.33 | 99.49 | 99.15 | 97.62 | 99.15 | 98.32 | 96.24 | 98.71 |
| Microclass | 99.08 | 98.23 | 99.08 | 96.98 | 99.12 | 99.08 | 97.28 | 99.44 | 99.08 | 97.60 | 99.08 | 98.19 | 96.16 | 98.59 |
| NBC50 | 99.08 | 98.23 | 99.08 | 96.98 | 99.12 | 99.08 | 97.28 | 99.44 | 99.08 | 97.60 | 99.08 | 98.19 | 96.16 | 98.59 |
| Q2_SK | 99.08 | 98.23 | 99.08 | 96.98 | 99.12 | 99.08 | 97.28 | 99.44 | 99.08 | 97.60 | 99.08 | 98.19 | 96.16 | 98.59 |
| RDP50 | 99.08 | 98.23 | 99.08 | 96.98 | 99.12 | 99.08 | 97.28 | 99.44 | 99.08 | 97.60 | 99.08 | 98.19 | 96.16 | 98.59 |
| RDP80 | 99.08 | 98.23 | 99.08 | 96.42 | 99.19 | 99.08 | 96.78 | 99.55 | 99.08 | 96.98 | 99.08 | 98.19 | 95.67 | 98.71 |
| NBC80 | 99.01 | 98.20 | 99.01 | 96.62 | 99.17 | 99.01 | 97.09 | 99.58 | 99.01 | 96.95 | 99.01 | 98.05 | 95.96 | 98.67 |
| SINTAX50 | 98.94 | 98.09 | 98.94 | 96.35 | 99.11 | 98.94 | 96.78 | 99.55 | 98.94 | 96.84 | 98.94 | 97.91 | 95.53 | 98.56 |
| HiTaC | 98.94 | 97.96 | 98.94 | 96.11 | 99.02 | 98.94 | 96.55 | 99.43 | 98.94 | 96.71 | 98.94 | 97.91 | 95.17 | 98.45 |
| HiTaC_Filter | 98.87 | 97.93 | 98.87 | 95.62 | 99.05 | 98.87 | 96.16 | 99.55 | 98.87 | 96.07 | 98.87 | 97.77 | 94.77 | 98.49 |
| SPINGO | 98.73 | 97.79 | 98.73 | 96.15 | 98.99 | 98.73 | 96.77 | 99.54 | 98.73 | 96.54 | 98.73 | 97.50 | 95.23 | 98.36 |
| CT1 | 98.66 | 95.24 | 98.66 | 93.44 | 98.66 | 98.66 | 93.84 | 98.95 | 98.66 | 94.03 | 98.66 | 97.36 | 92.36 | 97.98 |
| Q1 | 98.45 | 97.76 | 98.45 | 95.97 | 98.60 | 98.45 | 96.44 | 99.09 | 98.45 | 96.52 | 98.45 | 96.95 | 94.88 | 97.64 |
| SINTAX80 | 98.31 | 96.58 | 98.31 | 95.75 | 98.67 | 98.31 | 96.76 | 99.48 | 98.31 | 95.96 | 98.31 | 96.68 | 94.64 | 97.93 |
| Metaxa2 | 98.10 | 93.36 | 98.10 | 92.23 | 98.47 | 98.10 | 93.27 | 99.16 | 98.10 | 92.17 | 98.10 | 96.27 | 91.18 | 97.76 |
| Q2_VS | 97.39 | 93.75 | 97.39 | 91.87 | 97.63 | 97.39 | 93.12 | 98.43 | 97.39 | 91.97 | 97.39 | 94.92 | 90.16 | 96.19 |
| Q2_BLAST | 97.39 | 93.70 | 97.39 | 90.85 | 97.66 | 97.39 | 91.68 | 98.45 | 97.39 | 91.34 | 97.39 | 94.92 | 89.31 | 96.40 |
| CT2 | 96.41 | 89.39 | 96.41 | 88.18 | 96.59 | 96.41 | 90.06 | 97.68 | 96.41 | 88.25 | 96.41 | 93.07 | 85.96 | 94.80 |
| BLCA | 93.38 | 90.80 | 93.38 | 89.55 | 95.45 | 93.38 | 91.10 | 98.74 | 93.38 | 89.08 | 93.38 | 87.58 | 87.99 | 93.02 |
| KNN | 85.63 | 64.32 | 85.63 | 66.23 | 88.40 | 85.63 | 71.56 | 93.77 | 85.63 | 63.90 | 85.63 | 74.88 | 63.78 | 85.46 |

**Table S78. Machine learning metrics computed for the dataset SP RDP ITS 97 at the genus level.**

| Method | Accuracy | Balanced Accuracy | F1-score Micro | F1-score Macro | F1-score Weighted | Precision Micro | Precision Macro | Precision Weighted | Recall Micro | Recall Macro | Recall Weighted | Jaccard Micro | Jaccard Macro | Jaccard Weighted |
|---|---|---|---|---|---|---|---|---|---|---|---|---|---|---|
| HiTaC | 91.55 | 89.31 | 91.55 | 84.36 | 90.79 | 91.55 | 85.15 | 91.67 | 91.55 | 85.07 | 91.55 | 84.42 | 82.67 | 87.69 |
| Microclass | 91.06 | 89.41 | 91.06 | 84.24 | 90.74 | 91.06 | 85.25 | 91.99 | 91.06 | 84.72 | 91.06 | 83.58 | 82.49 | 87.23 |
| TOP | 90.99 | 89.33 | 90.99 | 84.40 | 90.42 | 90.99 | 85.49 | 91.58 | 90.99 | 84.86 | 90.99 | 83.46 | 82.70 | 87.12 |
| KTOP | 90.99 | 89.33 | 90.99 | 83.85 | 90.46 | 90.99 | 84.79 | 91.49 | 90.99 | 84.42 | 90.99 | 83.46 | 82.09 | 87.07 |
| BTOP | 90.85 | 89.60 | 90.85 | 84.53 | 90.38 | 90.85 | 85.55 | 91.38 | 90.85 | 84.90 | 90.85 | 83.24 | 82.87 | 86.93 |
| NBC50 | 90.56 | 88.93 | 90.56 | 83.93 | 90.21 | 90.56 | 84.75 | 91.37 | 90.56 | 84.48 | 90.56 | 82.75 | 82.24 | 86.91 |
| RDP50 | 90.49 | 88.82 | 90.49 | 84.21 | 90.25 | 90.49 | 85.12 | 91.41 | 90.49 | 84.60 | 90.49 | 82.64 | 82.49 | 86.89 |
| Q2_SK | 89.08 | 88.05 | 89.08 | 83.73 | 89.23 | 89.08 | 84.78 | 91.09 | 89.08 | 84.09 | 89.08 | 80.32 | 82.03 | 85.77 |
| SINTAX50 | 88.94 | 86.85 | 88.94 | 83.12 | 89.01 | 88.94 | 84.69 | 91.02 | 88.94 | 83.17 | 88.94 | 80.09 | 81.30 | 85.53 |
| CT1 | 87.89 | 80.70 | 87.89 | 73.99 | 87.24 | 87.89 | 74.78 | 88.41 | 87.89 | 74.70 | 87.89 | 78.39 | 72.07 | 83.55 |
| HiTaC_Filter | 87.68 | 85.02 | 87.68 | 82.36 | 88.63 | 87.68 | 84.13 | 91.30 | 87.68 | 82.07 | 87.68 | 78.06 | 80.66 | 85.16 |
| Q1 | 87.18 | 82.75 | 87.18 | 75.23 | 86.91 | 87.18 | 76.43 | 88.63 | 87.18 | 75.63 | 87.18 | 77.28 | 73.06 | 82.73 |
| SPINGO | 86.48 | 84.67 | 86.48 | 82.03 | 87.85 | 86.48 | 84.56 | 91.69 | 86.48 | 81.50 | 86.48 | 76.18 | 80.10 | 84.16 |
| NBC80 | 86.41 | 85.42 | 86.41 | 82.63 | 87.60 | 86.41 | 84.76 | 91.15 | 86.41 | 82.24 | 86.41 | 76.07 | 80.96 | 84.13 |
| RDP80 | 86.34 | 84.90 | 86.34 | 82.29 | 87.50 | 86.34 | 84.33 | 91.00 | 86.34 | 81.95 | 86.34 | 75.96 | 80.57 | 84.07 |
| Metaxa2 | 84.93 | 78.12 | 84.93 | 74.57 | 85.65 | 84.93 | 76.62 | 88.17 | 84.93 | 74.22 | 84.93 | 73.81 | 72.62 | 81.69 |
| BLCA | 82.96 | 79.28 | 82.96 | 75.61 | 84.27 | 82.96 | 78.68 | 88.81 | 82.96 | 74.92 | 82.96 | 70.88 | 73.47 | 80.50 |
| SINTAX80 | 82.25 | 78.99 | 82.25 | 77.74 | 83.93 | 82.25 | 80.18 | 88.36 | 82.25 | 77.07 | 82.25 | 69.86 | 76.00 | 80.48 |
| Q2_BLAST | 78.03 | 65.61 | 78.03 | 61.17 | 77.67 | 78.03 | 62.09 | 79.97 | 78.03 | 62.17 | 78.03 | 63.97 | 58.94 | 73.27 |
| Q2_VS | 77.89 | 65.86 | 77.89 | 62.48 | 77.83 | 77.89 | 64.02 | 80.80 | 77.89 | 63.06 | 77.89 | 63.78 | 60.26 | 73.33 |
| CT2 | 75.99 | 61.01 | 75.99 | 56.97 | 75.40 | 75.99 | 58.02 | 78.05 | 75.99 | 57.96 | 75.99 | 61.27 | 54.94 | 71.13 |
| KNN | 48.03 | 30.70 | 48.03 | 31.76 | 51.49 | 48.03 | 34.66 | 59.27 | 48.03 | 30.62 | 48.03 | 31.60 | 30.18 | 47.16 |

**Table S79. Machine learning metrics computed for the dataset SP RDP ITS 97 at the species level.**

| Method | Accuracy | Balanced Accuracy | F1-score Micro | F1-score Macro | F1-score Weighted | Precision Micro | Precision Macro | Precision Weighted | Recall Micro | Recall Macro | Recall Weighted | Jaccard Micro | Jaccard Macro | Jaccard Weighted |
|---|---|---|---|---|---|---|---|---|---|---|---|---|---|---|
| TOP | 3.66 | 4.94 | 3.66 | 2.75 | 3.54 | 3.66 | 2.71 | 3.50 | 3.66 | 2.84 | 3.66 | 1.87 | 2.71 | 3.50 |
| HiTaC | 3.66 | 4.89 | 3.66 | 2.75 | 3.60 | 3.66 | 2.73 | 3.58 | 3.66 | 2.80 | 3.66 | 1.87 | 2.72 | 3.56 |
| BTOP | 3.66 | 4.84 | 3.66 | 2.66 | 3.50 | 3.66 | 2.62 | 3.44 | 3.66 | 2.78 | 3.66 | 1.86 | 2.62 | 3.44 |
| Microclass | 3.59 | 4.84 | 3.59 | 2.73 | 3.48 | 3.59 | 2.70 | 3.45 | 3.59 | 2.81 | 3.59 | 1.83 | 2.70 | 3.45 |
| KTOP | 3.52 | 4.75 | 3.52 | 2.64 | 3.35 | 3.52 | 2.59 | 3.30 | 3.52 | 2.77 | 3.52 | 1.79 | 2.59 | 3.30 |
| Q2_SK | 3.45 | 4.65 | 3.45 | 3.01 | 3.39 | 3.45 | 2.99 | 3.37 | 3.45 | 3.06 | 3.45 | 1.76 | 2.99 | 3.37 |
| RDP50 | 3.45 | 4.65 | 3.45 | 2.88 | 3.39 | 3.45 | 2.86 | 3.37 | 3.45 | 2.93 | 3.45 | 1.76 | 2.86 | 3.37 |
| NBC50 | 3.45 | 4.65 | 3.45 | 2.86 | 3.39 | 3.45 | 2.84 | 3.37 | 3.45 | 2.91 | 3.45 | 1.76 | 2.84 | 3.37 |
| SINTAX50 | 3.17 | 4.27 | 3.17 | 2.84 | 3.05 | 3.17 | 2.79 | 3.00 | 3.17 | 2.95 | 3.17 | 1.61 | 2.79 | 3.00 |
| NBC80 | 3.03 | 4.08 | 3.03 | 2.94 | 3.00 | 3.03 | 2.93 | 2.99 | 3.03 | 2.96 | 3.03 | 1.54 | 2.93 | 2.99 |
| SPINGO | 3.03 | 4.08 | 3.03 | 2.82 | 2.93 | 3.03 | 2.79 | 2.90 | 3.03 | 2.91 | 3.03 | 1.54 | 2.79 | 2.90 |
| RDP80 | 2.96 | 3.99 | 2.96 | 2.88 | 2.93 | 2.96 | 2.86 | 2.92 | 2.96 | 2.90 | 2.96 | 1.50 | 2.86 | 2.92 |
| HiTaC_Filter | 2.89 | 3.89 | 2.89 | 2.92 | 2.84 | 2.89 | 2.90 | 2.82 | 2.89 | 2.97 | 2.89 | 1.46 | 2.90 | 2.82 |
| BLCA | 2.61 | 3.51 | 2.61 | 2.27 | 2.54 | 2.61 | 2.25 | 2.52 | 2.61 | 2.33 | 2.61 | 1.32 | 2.25 | 2.52 |
| SINTAX80 | 2.39 | 3.23 | 2.39 | 2.60 | 2.37 | 2.39 | 2.58 | 2.36 | 2.39 | 2.62 | 2.39 | 1.21 | 2.58 | 2.36 |
| Q1 | 1.69 | 2.28 | 1.69 | 1.40 | 1.62 | 1.69 | 1.37 | 1.58 | 1.69 | 1.46 | 1.69 | 0.85 | 1.37 | 1.58 |
| Q2_BLAST | 0.77 | 1.04 | 0.77 | 1.01 | 0.77 | 0.77 | 1.01 | 0.77 | 0.77 | 1.01 | 0.77 | 0.39 | 1.01 | 0.77 |
| Q2_VS | 0.70 | 0.95 | 0.70 | 0.92 | 0.70 | 0.70 | 0.92 | 0.70 | 0.70 | 0.92 | 0.70 | 0.35 | 0.92 | 0.70 |
| CT1 | 0.56 | 0.76 | 0.56 | 0.43 | 0.48 | 0.56 | 0.40 | 0.45 | 0.56 | 0.50 | 0.56 | 0.28 | 0.40 | 0.45 |
| Metaxa2 | 0.35 | 0.47 | 0.35 | 0.31 | 0.31 | 0.35 | 0.28 | 0.28 | 0.35 | 0.36 | 0.35 | 0.18 | 0.28 | 0.28 |
| CT2 | 0.21 | 0.28 | 0.21 | 0.28 | 0.21 | 0.21 | 0.28 | 0.21 | 0.21 | 0.28 | 0.21 | 0.11 | 0.28 | 0.21 |
| KNN | 0.00 | 0.00 | 0.00 | 0.00 | 0.00 | 0.00 | 0.00 | 0.00 | 0.00 | 0.00 | 0.00 | 0.00 | 0.00 | 0.00 |

**Table S80. Machine learning metrics computed for the dataset SP RDP ITS 99 at the phylum level.**

| Method | Accuracy | Balanced Accuracy | F1-score Micro | F1-score Macro | F1-score Weighted | Precision Micro | Precision Macro | Precision Weighted | Recall Micro | Recall Macro | Recall Weighted | Jaccard Micro | Jaccard Macro | Jaccard Weighted |
|---|---|---|---|---|---|---|---|---|---|---|---|---|---|---|
| BTOP | 100.00 | 100.00 | 100.00 | 100.00 | 100.00 | 100.00 | 100.00 | 100.00 | 100.00 | 100.00 | 100.00 | 100.00 | 100.00 | 100.00 |
| CT1 | 100.00 | 100.00 | 100.00 | 100.00 | 100.00 | 100.00 | 100.00 | 100.00 | 100.00 | 100.00 | 100.00 | 100.00 | 100.00 | 100.00 |
| CT2 | 100.00 | 100.00 | 100.00 | 100.00 | 100.00 | 100.00 | 100.00 | 100.00 | 100.00 | 100.00 | 100.00 | 100.00 | 100.00 | 100.00 |
| HiTaC | 100.00 | 100.00 | 100.00 | 100.00 | 100.00 | 100.00 | 100.00 | 100.00 | 100.00 | 100.00 | 100.00 | 100.00 | 100.00 | 100.00 |
| HiTaC_Filter | 100.00 | 100.00 | 100.00 | 100.00 | 100.00 | 100.00 | 100.00 | 100.00 | 100.00 | 100.00 | 100.00 | 100.00 | 100.00 | 100.00 |
| KTOP | 100.00 | 100.00 | 100.00 | 100.00 | 100.00 | 100.00 | 100.00 | 100.00 | 100.00 | 100.00 | 100.00 | 100.00 | 100.00 | 100.00 |
| Microclass | 100.00 | 100.00 | 100.00 | 100.00 | 100.00 | 100.00 | 100.00 | 100.00 | 100.00 | 100.00 | 100.00 | 100.00 | 100.00 | 100.00 |
| NBC50 | 100.00 | 100.00 | 100.00 | 100.00 | 100.00 | 100.00 | 100.00 | 100.00 | 100.00 | 100.00 | 100.00 | 100.00 | 100.00 | 100.00 |
| NBC80 | 100.00 | 100.00 | 100.00 | 100.00 | 100.00 | 100.00 | 100.00 | 100.00 | 100.00 | 100.00 | 100.00 | 100.00 | 100.00 | 100.00 |
| Q2_SK | 100.00 | 100.00 | 100.00 | 100.00 | 100.00 | 100.00 | 100.00 | 100.00 | 100.00 | 100.00 | 100.00 | 100.00 | 100.00 | 100.00 |
| RDP50 | 100.00 | 100.00 | 100.00 | 100.00 | 100.00 | 100.00 | 100.00 | 100.00 | 100.00 | 100.00 | 100.00 | 100.00 | 100.00 | 100.00 |
| RDP80 | 100.00 | 100.00 | 100.00 | 100.00 | 100.00 | 100.00 | 100.00 | 100.00 | 100.00 | 100.00 | 100.00 | 100.00 | 100.00 | 100.00 |
| SINTAX50 | 100.00 | 100.00 | 100.00 | 100.00 | 100.00 | 100.00 | 100.00 | 100.00 | 100.00 | 100.00 | 100.00 | 100.00 | 100.00 | 100.00 |
| SINTAX80 | 100.00 | 100.00 | 100.00 | 100.00 | 100.00 | 100.00 | 100.00 | 100.00 | 100.00 | 100.00 | 100.00 | 100.00 | 100.00 | 100.00 |
| TOP | 100.00 | 100.00 | 100.00 | 100.00 | 100.00 | 100.00 | 100.00 | 100.00 | 100.00 | 100.00 | 100.00 | 100.00 | 100.00 | 100.00 |
| Metaxa2 | 99.97 | 95.00 | 99.97 | 80.95 | 99.99 | 99.97 | 83.33 | 100.00 | 99.97 | 79.17 | 99.97 | 99.95 | 79.17 | 99.97 |
| Q2_BLAST | 99.90 | 99.96 | 99.90 | 83.32 | 99.95 | 99.90 | 83.33 | 100.00 | 99.90 | 83.30 | 99.90 | 99.80 | 83.30 | 99.90 |
| Q2_VS | 99.87 | 99.95 | 99.87 | 83.31 | 99.94 | 99.87 | 83.33 | 100.00 | 99.87 | 83.29 | 99.87 | 99.75 | 83.29 | 99.87 |
| KNN | 99.77 | 69.56 | 99.77 | 60.93 | 99.86 | 99.77 | 66.67 | 99.97 | 99.77 | 57.97 | 99.77 | 99.54 | 57.97 | 99.77 |
| Q1 | 99.75 | 99.90 | 99.75 | 83.29 | 99.87 | 99.75 | 83.33 | 100.00 | 99.75 | 83.25 | 99.75 | 99.49 | 83.25 | 99.75 |
| BLCA | 95.21 | 89.50 | 95.21 | 78.46 | 97.51 | 95.21 | 83.33 | 100.00 | 95.21 | 74.58 | 95.21 | 90.86 | 74.58 | 95.21 |
| SPINGO | 0.00 | 0.00 | 0.00 | 0.00 | 0.00 | 0.00 | 0.00 | 0.00 | 0.00 | 0.00 | 0.00 | 0.00 | 0.00 | 0.00 |

 

**Table S81. Machine learning metrics computed for the dataset SP RDP ITS 99 at the class level.**

| Method | Accuracy | Balanced Accuracy | F1-score Micro | F1-score Macro | F1-score Weighted | Precision Micro | Precision Macro | Precision Weighted | Recall Micro | Recall Macro | Recall Weighted | Jaccard Micro | Jaccard Macro | Jaccard Weighted |
|---|---|---|---|---|---|---|---|---|---|---|---|---|---|---|
| BTOP | 100.00 | 100.00 | 100.00 | 100.00 | 100.00 | 100.00 | 100.00 | 100.00 | 100.00 | 100.00 | 100.00 | 100.00 | 100.00 | 100.00 |
| HiTaC | 100.00 | 100.00 | 100.00 | 100.00 | 100.00 | 100.00 | 100.00 | 100.00 | 100.00 | 100.00 | 100.00 | 100.00 | 100.00 | 100.00 |
| HiTaC_Filter | 100.00 | 100.00 | 100.00 | 100.00 | 100.00 | 100.00 | 100.00 | 100.00 | 100.00 | 100.00 | 100.00 | 100.00 | 100.00 | 100.00 |
| KTOP | 100.00 | 100.00 | 100.00 | 100.00 | 100.00 | 100.00 | 100.00 | 100.00 | 100.00 | 100.00 | 100.00 | 100.00 | 100.00 | 100.00 |
| Microclass | 100.00 | 100.00 | 100.00 | 100.00 | 100.00 | 100.00 | 100.00 | 100.00 | 100.00 | 100.00 | 100.00 | 100.00 | 100.00 | 100.00 |
| NBC50 | 100.00 | 100.00 | 100.00 | 100.00 | 100.00 | 100.00 | 100.00 | 100.00 | 100.00 | 100.00 | 100.00 | 100.00 | 100.00 | 100.00 |
| NBC80 | 100.00 | 100.00 | 100.00 | 100.00 | 100.00 | 100.00 | 100.00 | 100.00 | 100.00 | 100.00 | 100.00 | 100.00 | 100.00 | 100.00 |
| Q2_SK | 100.00 | 100.00 | 100.00 | 100.00 | 100.00 | 100.00 | 100.00 | 100.00 | 100.00 | 100.00 | 100.00 | 100.00 | 100.00 | 100.00 |
| RDP50 | 100.00 | 100.00 | 100.00 | 100.00 | 100.00 | 100.00 | 100.00 | 100.00 | 100.00 | 100.00 | 100.00 | 100.00 | 100.00 | 100.00 |
| RDP80 | 100.00 | 100.00 | 100.00 | 100.00 | 100.00 | 100.00 | 100.00 | 100.00 | 100.00 | 100.00 | 100.00 | 100.00 | 100.00 | 100.00 |
| SINTAX50 | 100.00 | 100.00 | 100.00 | 100.00 | 100.00 | 100.00 | 100.00 | 100.00 | 100.00 | 100.00 | 100.00 | 100.00 | 100.00 | 100.00 |
| TOP | 100.00 | 100.00 | 100.00 | 100.00 | 100.00 | 100.00 | 100.00 | 100.00 | 100.00 | 100.00 | 100.00 | 100.00 | 100.00 | 100.00 |
| SINTAX80 | 99.97 | 99.99 | 99.97 | 96.15 | 99.99 | 99.97 | 96.15 | 100.00 | 99.97 | 96.14 | 99.97 | 99.95 | 96.14 | 99.97 |
| CT1 | 99.87 | 98.23 | 99.87 | 98.73 | 99.87 | 99.87 | 99.51 | 99.88 | 99.87 | 98.23 | 99.87 | 99.75 | 97.75 | 99.75 |
| Q2_VS | 99.70 | 99.50 | 99.70 | 95.74 | 99.76 | 99.70 | 95.83 | 99.82 | 99.70 | 95.67 | 99.70 | 99.39 | 95.35 | 99.52 |
| Q1 | 99.70 | 98.91 | 99.70 | 95.48 | 99.82 | 99.70 | 95.87 | 99.95 | 99.70 | 95.10 | 99.70 | 99.39 | 94.89 | 99.65 |
| Q2_BLAST | 99.70 | 98.17 | 99.70 | 94.98 | 99.76 | 99.70 | 95.83 | 99.82 | 99.70 | 94.39 | 99.70 | 99.39 | 94.07 | 99.52 |
| CT2 | 99.70 | 98.16 | 99.70 | 94.87 | 99.72 | 99.70 | 95.62 | 99.75 | 99.70 | 94.38 | 99.70 | 99.39 | 93.87 | 99.45 |
| Metaxa2 | 99.65 | 92.82 | 99.65 | 90.52 | 99.80 | 99.65 | 92.31 | 99.97 | 99.65 | 89.25 | 99.65 | 99.29 | 89.25 | 99.65 |
| KNN | 97.85 | 66.37 | 97.85 | 65.80 | 98.75 | 97.85 | 69.23 | 99.75 | 97.85 | 63.82 | 97.85 | 95.78 | 63.82 | 97.85 |
| BLCA | 95.18 | 91.29 | 95.18 | 89.76 | 97.36 | 95.18 | 92.31 | 99.95 | 95.18 | 87.78 | 95.18 | 90.81 | 87.78 | 95.18 |
| SPINGO | 0.00 | 0.00 | 0.00 | 0.00 | 0.00 | 0.00 | 0.00 | 0.00 | 0.00 | 0.00 | 0.00 | 0.00 | 0.00 | 0.00 |

**Table S82. Machine learning metrics computed for the dataset SP RDP ITS 99 at the order level.**

| Method | Accuracy | Balanced Accuracy | F1-score Micro | F1-score Macro | F1-score Weighted | Precision Micro | Precision Macro | Precision Weighted | Recall Micro | Recall Macro | Recall Weighted | Jaccard Micro | Jaccard Macro | Jaccard Weighted |
|---|---|---|---|---|---|---|---|---|---|---|---|---|---|---|
| BTOP | 99.97 | 99.97 | 99.97 | 99.98 | 99.97 | 99.97 | 99.99 | 99.98 | 99.97 | 99.97 | 99.97 | 99.95 | 99.97 | 99.95 |
| HiTaC | 99.97 | 99.97 | 99.97 | 99.98 | 99.97 | 99.97 | 99.99 | 99.97 | 99.97 | 99.97 | 99.97 | 99.95 | 99.97 | 99.95 |
| HiTaC_Filter | 99.97 | 99.97 | 99.97 | 99.98 | 99.97 | 99.97 | 99.99 | 99.97 | 99.97 | 99.97 | 99.97 | 99.95 | 99.97 | 99.95 |
| Microclass | 99.97 | 99.97 | 99.97 | 99.98 | 99.97 | 99.97 | 99.99 | 99.97 | 99.97 | 99.97 | 99.97 | 99.95 | 99.97 | 99.95 |
| KTOP | 99.95 | 99.97 | 99.95 | 99.98 | 99.95 | 99.95 | 99.99 | 99.95 | 99.95 | 99.97 | 99.95 | 99.90 | 99.95 | 99.90 |
| NBC50 | 99.95 | 99.97 | 99.95 | 99.98 | 99.95 | 99.95 | 99.99 | 99.95 | 99.95 | 99.97 | 99.95 | 99.90 | 99.95 | 99.90 |
| RDP50 | 99.95 | 99.97 | 99.95 | 99.98 | 99.95 | 99.95 | 99.99 | 99.95 | 99.95 | 99.97 | 99.95 | 99.90 | 99.95 | 99.90 |
| TOP | 99.95 | 99.97 | 99.95 | 99.98 | 99.95 | 99.95 | 99.99 | 99.95 | 99.95 | 99.97 | 99.95 | 99.90 | 99.95 | 99.90 |
| Q2_SK | 99.95 | 99.97 | 99.95 | 98.82 | 99.96 | 99.95 | 98.83 | 99.97 | 99.95 | 98.80 | 99.95 | 99.90 | 98.80 | 99.92 |
| SINTAX50 | 99.95 | 99.97 | 99.95 | 98.82 | 99.96 | 99.95 | 98.83 | 99.97 | 99.95 | 98.80 | 99.95 | 99.90 | 98.80 | 99.92 |
| SINTAX80 | 99.92 | 99.96 | 99.92 | 98.81 | 99.95 | 99.92 | 98.83 | 99.97 | 99.92 | 98.80 | 99.92 | 99.85 | 98.79 | 99.90 |
| NBC80 | 99.92 | 99.96 | 99.92 | 98.81 | 99.94 | 99.92 | 98.83 | 99.95 | 99.92 | 98.80 | 99.92 | 99.85 | 98.78 | 99.87 |
| RDP80 | 99.92 | 99.96 | 99.92 | 98.81 | 99.94 | 99.92 | 98.83 | 99.95 | 99.92 | 98.80 | 99.92 | 99.85 | 98.78 | 99.87 |
| Q1 | 99.39 | 99.22 | 99.39 | 98.31 | 99.54 | 99.39 | 98.58 | 99.70 | 99.39 | 98.07 | 99.39 | 98.79 | 97.83 | 99.10 |
| CT1 | 99.29 | 97.29 | 99.29 | 95.31 | 99.33 | 99.29 | 96.30 | 99.43 | 99.29 | 95.06 | 99.29 | 98.59 | 93.68 | 98.73 |
| Q2_VS | 99.04 | 96.20 | 99.04 | 95.44 | 99.13 | 99.04 | 96.24 | 99.27 | 99.04 | 95.08 | 99.04 | 98.09 | 93.86 | 98.41 |
| Q2_BLAST | 98.91 | 95.71 | 98.91 | 95.15 | 99.02 | 98.91 | 96.22 | 99.19 | 98.91 | 94.60 | 98.91 | 97.84 | 93.35 | 98.21 |
| Metaxa2 | 98.66 | 87.63 | 98.66 | 88.26 | 99.14 | 98.66 | 90.69 | 99.70 | 98.66 | 86.61 | 98.66 | 97.35 | 86.61 | 98.63 |
| CT2 | 98.45 | 89.42 | 98.45 | 88.31 | 98.59 | 98.45 | 91.04 | 98.87 | 98.45 | 87.37 | 98.45 | 96.95 | 85.54 | 97.58 |
| BLCA | 95.16 | 94.55 | 95.16 | 95.16 | 97.20 | 95.16 | 97.67 | 99.91 | 95.16 | 93.46 | 95.16 | 90.76 | 93.45 | 95.14 |
| KNN | 93.16 | 56.50 | 93.16 | 58.16 | 95.37 | 93.16 | 61.62 | 97.97 | 93.16 | 55.84 | 93.16 | 87.19 | 55.83 | 93.13 |
| SPINGO | 0.00 | 0.00 | 0.00 | 0.00 | 0.00 | 0.00 | 0.00 | 0.00 | 0.00 | 0.00 | 0.00 | 0.00 | 0.00 | 0.00 |

**Table S83. Machine learning metrics computed for the dataset SP RDP ITS 99 at the family level.**

| Method | Accuracy | Balanced Accuracy | F1-score Micro | F1-score Macro | F1-score Weighted | Precision Micro | Precision Macro | Precision Weighted | Recall Micro | Recall Macro | Recall Weighted | Jaccard Micro | Jaccard Macro | Jaccard Weighted |
|---|---|---|---|---|---|---|---|---|---|---|---|---|---|---|
| BTOP | 99.97 | 99.94 | 99.97 | 99.97 | 99.97 | 99.97 | 100.00 | 99.98 | 99.97 | 99.94 | 99.97 | 99.95 | 99.94 | 99.95 |
| HiTaC | 99.97 | 99.94 | 99.97 | 99.97 | 99.97 | 99.97 | 100.00 | 99.97 | 99.97 | 99.94 | 99.97 | 99.95 | 99.94 | 99.95 |
| HiTaC_Filter | 99.95 | 99.94 | 99.95 | 99.55 | 99.96 | 99.95 | 99.58 | 99.97 | 99.95 | 99.52 | 99.95 | 99.90 | 99.52 | 99.92 |
| Microclass | 99.95 | 99.92 | 99.95 | 99.93 | 99.95 | 99.95 | 99.95 | 99.95 | 99.95 | 99.92 | 99.95 | 99.90 | 99.88 | 99.90 |
| KTOP | 99.92 | 99.92 | 99.92 | 99.93 | 99.92 | 99.92 | 99.95 | 99.93 | 99.92 | 99.92 | 99.92 | 99.85 | 99.87 | 99.85 |
| NBC50 | 99.92 | 99.92 | 99.92 | 99.93 | 99.92 | 99.92 | 99.95 | 99.93 | 99.92 | 99.92 | 99.92 | 99.85 | 99.87 | 99.85 |
| RDP50 | 99.92 | 99.92 | 99.92 | 99.93 | 99.92 | 99.92 | 99.95 | 99.93 | 99.92 | 99.92 | 99.92 | 99.85 | 99.87 | 99.85 |
| TOP | 99.92 | 99.92 | 99.92 | 99.87 | 99.93 | 99.92 | 99.85 | 99.93 | 99.92 | 99.92 | 99.92 | 99.85 | 99.77 | 99.86 |
| Q2_SK | 99.90 | 99.87 | 99.90 | 99.49 | 99.92 | 99.90 | 99.53 | 99.95 | 99.90 | 99.46 | 99.90 | 99.80 | 99.41 | 99.85 |
| NBC80 | 99.87 | 99.87 | 99.87 | 99.51 | 99.92 | 99.87 | 99.58 | 99.97 | 99.87 | 99.45 | 99.87 | 99.75 | 99.45 | 99.85 |
| SINTAX50 | 99.87 | 99.87 | 99.87 | 99.51 | 99.92 | 99.87 | 99.58 | 99.97 | 99.87 | 99.45 | 99.87 | 99.75 | 99.45 | 99.85 |
| SPINGO | 99.87 | 99.87 | 99.87 | 99.51 | 99.92 | 99.87 | 99.58 | 99.97 | 99.87 | 99.45 | 99.87 | 99.75 | 99.45 | 99.85 |
| RDP80 | 99.87 | 99.87 | 99.87 | 99.49 | 99.91 | 99.87 | 99.53 | 99.95 | 99.87 | 99.45 | 99.87 | 99.75 | 99.41 | 99.83 |
| SINTAX80 | 99.80 | 99.73 | 99.80 | 99.44 | 99.88 | 99.80 | 99.58 | 99.97 | 99.80 | 99.32 | 99.80 | 99.60 | 99.32 | 99.77 |
| Q1 | 99.06 | 98.67 | 99.06 | 98.18 | 99.21 | 99.06 | 98.24 | 99.42 | 99.06 | 98.27 | 99.06 | 98.14 | 97.36 | 98.52 |
| CT1 | 97.95 | 91.67 | 97.95 | 90.84 | 97.99 | 97.95 | 91.94 | 98.31 | 97.95 | 90.91 | 97.95 | 95.98 | 88.42 | 96.60 |
| Metaxa2 | 96.22 | 84.42 | 96.22 | 86.31 | 97.59 | 96.22 | 89.67 | 99.29 | 96.22 | 84.07 | 96.22 | 92.72 | 84.07 | 96.20 |
| Q2_VS | 96.10 | 85.90 | 96.10 | 85.98 | 96.45 | 96.10 | 87.93 | 97.23 | 96.10 | 85.19 | 96.10 | 92.49 | 82.96 | 94.29 |
| Q2_BLAST | 95.56 | 84.96 | 95.56 | 85.19 | 96.09 | 95.56 | 87.84 | 97.28 | 95.56 | 84.26 | 95.56 | 91.50 | 81.85 | 93.75 |
| BLCA | 95.13 | 95.48 | 95.13 | 96.60 | 97.08 | 95.13 | 99.17 | 99.91 | 95.13 | 95.08 | 95.13 | 90.72 | 95.08 | 95.12 |
| CT2 | 93.81 | 77.40 | 93.81 | 78.02 | 94.44 | 93.81 | 82.26 | 96.00 | 93.81 | 76.45 | 93.81 | 88.35 | 73.78 | 91.58 |
| KNN | 80.84 | 45.20 | 80.84 | 48.58 | 85.31 | 80.84 | 55.78 | 92.72 | 80.84 | 45.01 | 80.84 | 67.84 | 45.01 | 80.81 |

**Table S84. Machine learning metrics computed for the dataset SP RDP ITS 99 at the genus level.**

| Method | Accuracy | Balanced Accuracy | F1-score Micro | F1-score Macro | F1-score Weighted | Precision Micro | Precision Macro | Precision Weighted | Recall Micro | Recall Macro | Recall Weighted | Jaccard Micro | Jaccard Macro | Jaccard Weighted |
|---|---|---|---|---|---|---|---|---|---|---|---|---|---|---|
| TOP | 99.52 | 99.64 | 99.52 | 99.05 | 99.58 | 99.52 | 99.11 | 99.69 | 99.52 | 99.04 | 99.52 | 99.04 | 98.89 | 99.27 |
| HiTaC | 99.49 | 99.48 | 99.49 | 99.08 | 99.51 | 99.49 | 99.15 | 99.61 | 99.49 | 99.12 | 99.49 | 98.99 | 98.88 | 99.17 |
| Microclass | 99.44 | 99.45 | 99.44 | 98.96 | 99.48 | 99.44 | 99.04 | 99.58 | 99.44 | 98.97 | 99.44 | 98.89 | 98.75 | 99.11 |
| BTOP | 99.37 | 99.49 | 99.37 | 98.86 | 99.42 | 99.37 | 98.89 | 99.51 | 99.37 | 98.89 | 99.37 | 98.76 | 98.66 | 99.02 |
| KTOP | 99.37 | 99.42 | 99.37 | 98.82 | 99.43 | 99.37 | 98.90 | 99.55 | 99.37 | 98.83 | 99.37 | 98.74 | 98.60 | 99.02 |
| RDP50 | 99.32 | 99.38 | 99.32 | 98.88 | 99.39 | 99.32 | 99.02 | 99.58 | 99.32 | 98.90 | 99.32 | 98.64 | 98.64 | 98.97 |
| NBC50 | 99.29 | 99.37 | 99.29 | 98.86 | 99.35 | 99.29 | 98.98 | 99.52 | 99.29 | 98.89 | 99.29 | 98.59 | 98.61 | 98.91 |
| Q2_SK | 99.24 | 99.36 | 99.24 | 99.15 | 99.38 | 99.24 | 99.30 | 99.64 | 99.24 | 99.12 | 99.24 | 98.49 | 98.91 | 98.95 |
| SINTAX50 | 99.21 | 99.35 | 99.21 | 98.89 | 99.39 | 99.21 | 99.05 | 99.69 | 99.21 | 98.87 | 99.21 | 98.44 | 98.64 | 98.96 |
| SPINGO | 98.99 | 99.24 | 98.99 | 99.26 | 99.27 | 98.99 | 99.55 | 99.73 | 98.99 | 99.12 | 98.99 | 97.99 | 99.03 | 98.79 |
| RDP80 | 98.86 | 99.12 | 98.86 | 99.03 | 99.23 | 98.86 | 99.30 | 99.76 | 98.86 | 98.88 | 98.86 | 97.74 | 98.79 | 98.71 |
| NBC80 | 98.81 | 99.08 | 98.81 | 99.01 | 99.22 | 98.81 | 99.31 | 99.79 | 98.81 | 98.84 | 98.81 | 97.65 | 98.75 | 98.68 |
| HiTaC_Filter | 98.78 | 98.83 | 98.78 | 98.99 | 99.19 | 98.78 | 99.47 | 99.80 | 98.78 | 98.71 | 98.78 | 97.60 | 98.68 | 98.68 |
| SINTAX80 | 97.97 | 98.08 | 97.97 | 98.30 | 98.60 | 97.97 | 99.03 | 99.72 | 97.97 | 97.96 | 97.97 | 96.02 | 97.96 | 97.90 |
| Q1 | 95.54 | 90.51 | 95.54 | 88.93 | 95.49 | 95.54 | 89.74 | 96.04 | 95.54 | 89.11 | 95.54 | 91.46 | 87.22 | 93.34 |
| BLCA | 94.32 | 95.26 | 94.32 | 95.69 | 95.99 | 94.32 | 97.30 | 99.07 | 94.32 | 94.91 | 94.32 | 89.25 | 94.75 | 94.16 |
| CT1 | 92.22 | 79.41 | 92.22 | 75.50 | 91.66 | 92.22 | 75.87 | 92.05 | 92.22 | 76.72 | 92.22 | 85.56 | 72.74 | 88.53 |
| Metaxa2 | 89.05 | 73.05 | 89.05 | 74.58 | 91.32 | 89.05 | 77.84 | 94.73 | 89.05 | 72.78 | 89.05 | 80.26 | 72.72 | 88.90 |
| Q2_VS | 81.42 | 58.70 | 81.42 | 57.74 | 81.20 | 81.42 | 59.06 | 82.86 | 81.42 | 58.21 | 81.42 | 68.66 | 54.94 | 76.70 |
| Q2_BLAST | 80.15 | 55.87 | 80.15 | 55.13 | 79.88 | 80.15 | 56.89 | 81.52 | 80.15 | 55.41 | 80.15 | 66.88 | 51.89 | 74.96 |
| CT2 | 77.08 | 47.32 | 77.08 | 46.66 | 76.91 | 77.08 | 48.79 | 79.21 | 77.08 | 46.70 | 77.08 | 62.71 | 43.72 | 72.02 |
| KNN | 47.98 | 15.58 | 47.98 | 17.03 | 53.01 | 47.98 | 20.14 | 62.89 | 47.98 | 15.56 | 47.98 | 31.57 | 15.56 | 47.96 |

**Table S85. Machine learning metrics computed for the dataset SP RDP ITS 99 at the species level.**

| Method | Accuracy | Balanced Accuracy | F1-score Micro | F1-score Macro | F1-score Weighted | Precision Micro | Precision Macro | Precision Weighted | Recall Micro | Recall Macro | Recall Weighted | Jaccard Micro | Jaccard Macro | Jaccard Weighted |
|---|---|---|---|---|---|---|---|---|---|---|---|---|---|---|
| HiTaC | 72.98 | 77.61 | 72.98 | 66.47 | 71.65 | 72.98 | 66.12 | 71.59 | 72.98 | 67.79 | 72.98 | 57.45 | 65.69 | 70.56 |
| BTOP | 72.59 | 76.78 | 72.59 | 65.60 | 71.13 | 72.59 | 65.17 | 71.02 | 72.59 | 67.05 | 72.59 | 56.97 | 64.78 | 70.03 |
| Microclass | 72.32 | 76.87 | 72.32 | 65.90 | 70.72 | 72.32 | 65.42 | 70.32 | 72.32 | 67.38 | 72.32 | 56.64 | 65.15 | 69.72 |
| TOP | 72.29 | 76.98 | 72.29 | 65.95 | 70.96 | 72.29 | 65.64 | 70.92 | 72.29 | 67.25 | 72.29 | 56.61 | 65.16 | 69.86 |
| KTOP | 71.63 | 76.35 | 71.63 | 65.39 | 70.06 | 71.63 | 64.89 | 69.70 | 71.63 | 66.89 | 71.63 | 55.81 | 64.56 | 68.96 |
| NBC50 | 68.59 | 73.81 | 68.59 | 63.59 | 66.98 | 68.59 | 63.06 | 66.51 | 68.59 | 65.10 | 68.59 | 52.20 | 62.85 | 66.05 |
| RDP50 | 68.29 | 73.51 | 68.29 | 63.36 | 66.73 | 68.29 | 62.86 | 66.30 | 68.29 | 64.82 | 68.29 | 51.85 | 62.64 | 65.81 |
| SINTAX50 | 68.24 | 73.34 | 68.24 | 65.11 | 67.26 | 68.24 | 64.80 | 67.13 | 68.24 | 66.10 | 68.24 | 51.79 | 64.51 | 66.50 |
| Q2_SK | 66.89 | 72.37 | 66.89 | 63.74 | 65.80 | 66.89 | 63.35 | 65.48 | 66.89 | 64.81 | 66.89 | 50.26 | 63.21 | 65.15 |
| SPINGO | 65.73 | 71.03 | 65.73 | 64.70 | 65.31 | 65.73 | 64.65 | 65.56 | 65.73 | 65.24 | 65.73 | 48.95 | 64.23 | 64.68 |
| NBC80 | 64.06 | 69.53 | 64.06 | 62.43 | 63.15 | 64.06 | 62.12 | 62.96 | 64.06 | 63.35 | 64.06 | 47.12 | 61.96 | 62.60 |
| RDP80 | 63.90 | 69.36 | 63.90 | 62.49 | 63.03 | 63.90 | 62.18 | 62.82 | 63.90 | 63.37 | 63.90 | 46.95 | 62.04 | 62.51 |
| HiTaC_Filter | 63.45 | 68.82 | 63.45 | 63.45 | 63.15 | 63.45 | 63.45 | 63.47 | 63.45 | 63.89 | 63.45 | 46.46 | 63.07 | 62.62 |
| BLCA | 63.07 | 67.98 | 63.07 | 60.21 | 62.32 | 63.07 | 60.01 | 62.43 | 63.07 | 61.10 | 63.07 | 46.06 | 59.58 | 61.47 |
| SINTAX80 | 57.90 | 63.39 | 57.90 | 59.76 | 57.73 | 57.90 | 59.79 | 58.01 | 57.90 | 60.06 | 57.90 | 40.74 | 59.44 | 57.31 |
| Q1 | 36.91 | 39.50 | 36.91 | 32.19 | 34.93 | 36.91 | 31.41 | 34.14 | 36.91 | 34.02 | 36.91 | 22.63 | 31.33 | 33.90 |
| CT1 | 27.10 | 28.42 | 27.10 | 21.62 | 24.50 | 27.10 | 20.67 | 23.47 | 27.10 | 23.93 | 27.10 | 15.67 | 20.59 | 23.28 |
| Metaxa2 | 23.12 | 24.39 | 23.12 | 22.58 | 22.69 | 23.12 | 22.45 | 22.71 | 23.12 | 23.05 | 23.12 | 13.07 | 22.27 | 22.32 |
| Q2_VS | 7.25 | 8.11 | 7.25 | 7.81 | 7.11 | 7.25 | 7.74 | 7.04 | 7.25 | 7.97 | 7.25 | 3.76 | 7.74 | 7.04 |
| Q2_BLAST | 5.75 | 6.43 | 5.75 | 6.22 | 5.65 | 5.75 | 6.17 | 5.61 | 5.75 | 6.33 | 5.75 | 2.96 | 6.17 | 5.61 |
| CT2 | 2.94 | 3.24 | 2.94 | 3.18 | 2.90 | 2.94 | 3.16 | 2.89 | 2.94 | 3.21 | 2.94 | 1.49 | 3.16 | 2.89 |
| KNN | 0.00 | 0.00 | 0.00 | 0.00 | 0.00 | 0.00 | 0.00 | 0.00 | 0.00 | 0.00 | 0.00 | 0.00 | 0.00 | 0.00 |

Table S86. Machine learning metrics computed for the dataset SP RDP ITS 100 at the phylum level.

| Method | Accuracy | Balanced Accuracy | F1-score Micro | F1-score Macro | F1-score Weighted | Precision Micro | Precision Macro | Precision Weighted | Recall Micro | Recall Macro | Recall Weighted | Jaccard Micro | Jaccard Macro | Jaccard Weighted |
|---|---|---|---|---|---|---|---|---|---|---|---|---|---|---|
| BTOP | 100.00 | 100.00 | 100.00 | 100.00 | 100.00 | 100.00 | 100.00 | 100.00 | 100.00 | 100.00 | 100.00 | 100.00 | 100.00 | 100.00 |
| CT1 | 100.00 | 100.00 | 100.00 | 100.00 | 100.00 | 100.00 | 100.00 | 100.00 | 100.00 | 100.00 | 100.00 | 100.00 | 100.00 | 100.00 |
| CT2 | 100.00 | 100.00 | 100.00 | 100.00 | 100.00 | 100.00 | 100.00 | 100.00 | 100.00 | 100.00 | 100.00 | 100.00 | 100.00 | 100.00 |
| HiTaC | 100.00 | 100.00 | 100.00 | 100.00 | 100.00 | 100.00 | 100.00 | 100.00 | 100.00 | 100.00 | 100.00 | 100.00 | 100.00 | 100.00 |
| HiTaC_Filter | 100.00 | 100.00 | 100.00 | 100.00 | 100.00 | 100.00 | 100.00 | 100.00 | 100.00 | 100.00 | 100.00 | 100.00 | 100.00 | 100.00 |
| KTOP | 100.00 | 100.00 | 100.00 | 100.00 | 100.00 | 100.00 | 100.00 | 100.00 | 100.00 | 100.00 | 100.00 | 100.00 | 100.00 | 100.00 |
| Metaxa2 | 100.00 | 100.00 | 100.00 | 100.00 | 100.00 | 100.00 | 100.00 | 100.00 | 100.00 | 100.00 | 100.00 | 100.00 | 100.00 | 100.00 |
| Microclass | 100.00 | 100.00 | 100.00 | 100.00 | 100.00 | 100.00 | 100.00 | 100.00 | 100.00 | 100.00 | 100.00 | 100.00 | 100.00 | 100.00 |
| Q1 | 100.00 | 100.00 | 100.00 | 100.00 | 100.00 | 100.00 | 100.00 | 100.00 | 100.00 | 100.00 | 100.00 | 100.00 | 100.00 | 100.00 |
| Q2_BLAST | 100.00 | 100.00 | 100.00 | 100.00 | 100.00 | 100.00 | 100.00 | 100.00 | 100.00 | 100.00 | 100.00 | 100.00 | 100.00 | 100.00 |
| Q2_SK | 100.00 | 100.00 | 100.00 | 100.00 | 100.00 | 100.00 | 100.00 | 100.00 | 100.00 | 100.00 | 100.00 | 100.00 | 100.00 | 100.00 |
| Q2_VS | 100.00 | 100.00 | 100.00 | 100.00 | 100.00 | 100.00 | 100.00 | 100.00 | 100.00 | 100.00 | 100.00 | 100.00 | 100.00 | 100.00 |
| RDP50 | 100.00 | 100.00 | 100.00 | 100.00 | 100.00 | 100.00 | 100.00 | 100.00 | 100.00 | 100.00 | 100.00 | 100.00 | 100.00 | 100.00 |
| RDP80 | 100.00 | 100.00 | 100.00 | 100.00 | 100.00 | 100.00 | 100.00 | 100.00 | 100.00 | 100.00 | 100.00 | 100.00 | 100.00 | 100.00 |
| SINTAX50 | 100.00 | 100.00 | 100.00 | 100.00 | 100.00 | 100.00 | 100.00 | 100.00 | 100.00 | 100.00 | 100.00 | 100.00 | 100.00 | 100.00 |
| SINTAX80 | 100.00 | 100.00 | 100.00 | 100.00 | 100.00 | 100.00 | 100.00 | 100.00 | 100.00 | 100.00 | 100.00 | 100.00 | 100.00 | 100.00 |
| TOP | 100.00 | 100.00 | 100.00 | 100.00 | 100.00 | 100.00 | 100.00 | 100.00 | 100.00 | 100.00 | 100.00 | 100.00 | 100.00 | 100.00 |
| KNN | 99.81 | 53.42 | 99.81 | 48.15 | 99.88 | 99.81 | 50.00 | 99.96 | 99.81 | 46.74 | 99.81 | 99.63 | 46.74 | 99.81 |
| BLCA | 0.00 | 0.00 | 0.00 | 0.00 | 0.00 | 0.00 | 0.00 | 0.00 | 0.00 | 0.00 | 0.00 | 0.00 | 0.00 | 0.00 |
| SPINGO | 0.00 | 0.00 | 0.00 | 0.00 | 0.00 | 0.00 | 0.00 | 0.00 | 0.00 | 0.00 | 0.00 | 0.00 | 0.00 | 0.00 |

**Table S87. Machine learning metrics computed for the dataset SP RDP ITS 100 at the class level.**

| Method | Accuracy | Balanced Accuracy | F1-score Micro | F1-score Macro | F1-score Weighted | Precision Micro | Precision Macro | Precision Weighted | Recall Micro | Recall Macro | Recall Weighted | Jaccard Micro | Jaccard Macro | Jaccard Weighted |
|---|---|---|---|---|---|---|---|---|---|---|---|---|---|---|
| BTOP | 100.00 | 100.00 | 100.00 | 100.00 | 100.00 | 100.00 | 100.00 | 100.00 | 100.00 | 100.00 | 100.00 | 100.00 | 100.00 | 100.00 |
| HiTaC | 100.00 | 100.00 | 100.00 | 100.00 | 100.00 | 100.00 | 100.00 | 100.00 | 100.00 | 100.00 | 100.00 | 100.00 | 100.00 | 100.00 |
| HiTaC_Filter | 100.00 | 100.00 | 100.00 | 100.00 | 100.00 | 100.00 | 100.00 | 100.00 | 100.00 | 100.00 | 100.00 | 100.00 | 100.00 | 100.00 |
| KTOP | 100.00 | 100.00 | 100.00 | 100.00 | 100.00 | 100.00 | 100.00 | 100.00 | 100.00 | 100.00 | 100.00 | 100.00 | 100.00 | 100.00 |
| Microclass | 100.00 | 100.00 | 100.00 | 100.00 | 100.00 | 100.00 | 100.00 | 100.00 | 100.00 | 100.00 | 100.00 | 100.00 | 100.00 | 100.00 |
| Q2_SK | 100.00 | 100.00 | 100.00 | 100.00 | 100.00 | 100.00 | 100.00 | 100.00 | 100.00 | 100.00 | 100.00 | 100.00 | 100.00 | 100.00 |
| RDP50 | 100.00 | 100.00 | 100.00 | 100.00 | 100.00 | 100.00 | 100.00 | 100.00 | 100.00 | 100.00 | 100.00 | 100.00 | 100.00 | 100.00 |
| RDP80 | 100.00 | 100.00 | 100.00 | 100.00 | 100.00 | 100.00 | 100.00 | 100.00 | 100.00 | 100.00 | 100.00 | 100.00 | 100.00 | 100.00 |
| SINTAX50 | 100.00 | 100.00 | 100.00 | 100.00 | 100.00 | 100.00 | 100.00 | 100.00 | 100.00 | 100.00 | 100.00 | 100.00 | 100.00 | 100.00 |
| SINTAX80 | 100.00 | 100.00 | 100.00 | 100.00 | 100.00 | 100.00 | 100.00 | 100.00 | 100.00 | 100.00 | 100.00 | 100.00 | 100.00 | 100.00 |
| TOP | 100.00 | 100.00 | 100.00 | 100.00 | 100.00 | 100.00 | 100.00 | 100.00 | 100.00 | 100.00 | 100.00 | 100.00 | 100.00 | 100.00 |
| Q1 | 99.99 | 100.00 | 99.99 | 100.00 | 99.99 | 99.99 | 100.00 | 99.99 | 99.99 | 100.00 | 99.99 | 99.99 | 99.99 | 99.99 |
| CT1 | 99.93 | 99.97 | 99.93 | 99.97 | 99.93 | 99.93 | 99.97 | 99.93 | 99.93 | 99.97 | 99.93 | 99.86 | 99.94 | 99.86 |
| Metaxa2 | 99.88 | 96.39 | 99.88 | 93.41 | 99.94 | 99.88 | 93.55 | 99.99 | 99.88 | 93.28 | 99.88 | 99.76 | 93.28 | 99.88 |
| Q2_VS | 99.74 | 99.64 | 99.74 | 96.46 | 99.75 | 99.74 | 96.51 | 99.77 | 99.74 | 96.42 | 99.74 | 99.48 | 96.16 | 99.51 |
| Q2_BLAST | 99.73 | 99.63 | 99.73 | 96.46 | 99.75 | 99.73 | 96.51 | 99.77 | 99.73 | 96.42 | 99.73 | 99.47 | 96.16 | 99.50 |
| CT2 | 99.67 | 99.46 | 99.67 | 96.34 | 99.69 | 99.67 | 96.44 | 99.70 | 99.67 | 96.25 | 99.67 | 99.34 | 95.92 | 99.38 |
| KNN | 98.15 | 57.88 | 98.15 | 57.92 | 98.91 | 98.15 | 61.29 | 99.77 | 98.15 | 56.01 | 98.15 | 96.37 | 56.01 | 98.15 |
| BLCA | 0.00 | 0.00 | 0.00 | 0.00 | 0.00 | 0.00 | 0.00 | 0.00 | 0.00 | 0.00 | 0.00 | 0.00 | 0.00 | 0.00 |
| SPINGO | 0.00 | 0.00 | 0.00 | 0.00 | 0.00 | 0.00 | 0.00 | 0.00 | 0.00 | 0.00 | 0.00 | 0.00 | 0.00 | 0.00 |

**Table S88. Machine learning metrics computed for the dataset SP RDP ITS 100 at the order level.**

| Method | Accuracy | Balanced Accuracy | F1-score Micro | F1-score Macro | F1-score Weighted | Precision Micro | Precision Macro | Precision Weighted | Recall Micro | Recall Macro | Recall Weighted | Jaccard Micro | Jaccard Macro | Jaccard Weighted |
|---|---|---|---|---|---|---|---|---|---|---|---|---|---|---|
| BTOP | 100.00 | 100.00 | 100.00 | 100.00 | 100.00 | 100.00 | 100.00 | 100.00 | 100.00 | 100.00 | 100.00 | 100.00 | 100.00 | 100.00 |
| HiTaC | 100.00 | 100.00 | 100.00 | 100.00 | 100.00 | 100.00 | 100.00 | 100.00 | 100.00 | 100.00 | 100.00 | 100.00 | 100.00 | 100.00 |
| HiTaC_Filter | 100.00 | 100.00 | 100.00 | 100.00 | 100.00 | 100.00 | 100.00 | 100.00 | 100.00 | 100.00 | 100.00 | 100.00 | 100.00 | 100.00 |
| KTOP | 100.00 | 100.00 | 100.00 | 100.00 | 100.00 | 100.00 | 100.00 | 100.00 | 100.00 | 100.00 | 100.00 | 100.00 | 100.00 | 100.00 |
| Microclass | 100.00 | 100.00 | 100.00 | 100.00 | 100.00 | 100.00 | 100.00 | 100.00 | 100.00 | 100.00 | 100.00 | 100.00 | 100.00 | 100.00 |
| Q2_SK | 100.00 | 100.00 | 100.00 | 100.00 | 100.00 | 100.00 | 100.00 | 100.00 | 100.00 | 100.00 | 100.00 | 100.00 | 100.00 | 100.00 |
| RDP50 | 100.00 | 100.00 | 100.00 | 100.00 | 100.00 | 100.00 | 100.00 | 100.00 | 100.00 | 100.00 | 100.00 | 100.00 | 100.00 | 100.00 |
| RDP80 | 100.00 | 100.00 | 100.00 | 100.00 | 100.00 | 100.00 | 100.00 | 100.00 | 100.00 | 100.00 | 100.00 | 100.00 | 100.00 | 100.00 |
| SINTAX50 | 100.00 | 100.00 | 100.00 | 100.00 | 100.00 | 100.00 | 100.00 | 100.00 | 100.00 | 100.00 | 100.00 | 100.00 | 100.00 | 100.00 |
| SINTAX80 | 100.00 | 100.00 | 100.00 | 100.00 | 100.00 | 100.00 | 100.00 | 100.00 | 100.00 | 100.00 | 100.00 | 100.00 | 100.00 | 100.00 |
| TOP | 100.00 | 100.00 | 100.00 | 100.00 | 100.00 | 100.00 | 100.00 | 100.00 | 100.00 | 100.00 | 100.00 | 100.00 | 100.00 | 100.00 |
| Q1 | 99.96 | 98.99 | 99.96 | 97.98 | 99.96 | 99.96 | 97.95 | 99.96 | 99.96 | 98.02 | 99.96 | 99.93 | 97.93 | 99.93 |
| CT1 | 99.79 | 96.50 | 99.79 | 95.55 | 99.79 | 99.79 | 95.59 | 99.79 | 99.79 | 95.55 | 99.79 | 99.59 | 95.07 | 99.60 |
| Metaxa2 | 99.69 | 94.19 | 99.69 | 93.67 | 99.83 | 99.69 | 94.12 | 99.97 | 99.69 | 93.26 | 99.69 | 99.39 | 93.26 | 99.69 |
| Q2_VS | 99.05 | 95.17 | 99.05 | 94.94 | 99.18 | 99.05 | 96.01 | 99.36 | 99.05 | 94.24 | 99.05 | 98.11 | 93.32 | 98.48 |
| Q2_BLAST | 98.96 | 92.88 | 98.96 | 92.73 | 99.11 | 98.96 | 93.98 | 99.31 | 98.96 | 91.97 | 98.96 | 97.94 | 90.97 | 98.35 |
| CT2 | 98.73 | 89.11 | 98.73 | 89.55 | 98.91 | 98.73 | 91.94 | 99.17 | 98.73 | 88.24 | 98.73 | 97.49 | 87.19 | 98.00 |
| KNN | 94.37 | 54.20 | 94.37 | 56.59 | 96.34 | 94.37 | 60.78 | 98.68 | 94.37 | 53.67 | 94.37 | 89.34 | 53.67 | 94.37 |
| BLCA | 0.00 | 0.00 | 0.00 | 0.00 | 0.00 | 0.00 | 0.00 | 0.00 | 0.00 | 0.00 | 0.00 | 0.00 | 0.00 | 0.00 |
| SPINGO | 0.00 | 0.00 | 0.00 | 0.00 | 0.00 | 0.00 | 0.00 | 0.00 | 0.00 | 0.00 | 0.00 | 0.00 | 0.00 | 0.00 |

**Table S89. Machine learning metrics computed for the dataset SP RDP ITS 100 at the family level.**

| Method | Accuracy | Balanced Accuracy | F1-score Micro | F1-score Macro | F1-score Weighted | Precision Micro | Precision Macro | Precision Weighted | Recall Micro | Recall Macro | Recall Weighted | Jaccard Micro | Jaccard Macro | Jaccard Weighted |
|---|---|---|---|---|---|---|---|---|---|---|---|---|---|---|
| BTOP | 100.00 | 100.00 | 100.00 | 100.00 | 100.00 | 100.00 | 100.00 | 100.00 | 100.00 | 100.00 | 100.00 | 100.00 | 100.00 | 100.00 |
| HiTaC | 100.00 | 100.00 | 100.00 | 100.00 | 100.00 | 100.00 | 100.00 | 100.00 | 100.00 | 100.00 | 100.00 | 100.00 | 100.00 | 100.00 |
| HiTaC_Filter | 100.00 | 100.00 | 100.00 | 100.00 | 100.00 | 100.00 | 100.00 | 100.00 | 100.00 | 100.00 | 100.00 | 100.00 | 100.00 | 100.00 |
| KTOP | 100.00 | 100.00 | 100.00 | 100.00 | 100.00 | 100.00 | 100.00 | 100.00 | 100.00 | 100.00 | 100.00 | 100.00 | 100.00 | 100.00 |
| Microclass | 100.00 | 100.00 | 100.00 | 100.00 | 100.00 | 100.00 | 100.00 | 100.00 | 100.00 | 100.00 | 100.00 | 100.00 | 100.00 | 100.00 |
| SINTAX50 | 100.00 | 100.00 | 100.00 | 100.00 | 100.00 | 100.00 | 100.00 | 100.00 | 100.00 | 100.00 | 100.00 | 100.00 | 100.00 | 100.00 |
| SPINGO | 100.00 | 100.00 | 100.00 | 100.00 | 100.00 | 100.00 | 100.00 | 100.00 | 100.00 | 100.00 | 100.00 | 100.00 | 100.00 | 100.00 |
| TOP | 100.00 | 100.00 | 100.00 | 100.00 | 100.00 | 100.00 | 100.00 | 100.00 | 100.00 | 100.00 | 100.00 | 100.00 | 100.00 | 100.00 |
| Q2_SK | 99.99 | 100.00 | 99.99 | 99.65 | 100.00 | 99.99 | 99.66 | 100.00 | 99.99 | 99.65 | 99.99 | 99.99 | 99.65 | 99.99 |
| RDP50 | 99.99 | 100.00 | 99.99 | 99.65 | 100.00 | 99.99 | 99.66 | 100.00 | 99.99 | 99.65 | 99.99 | 99.99 | 99.65 | 99.99 |
| RDP80 | 99.99 | 99.82 | 99.99 | 99.54 | 99.99 | 99.99 | 99.66 | 100.00 | 99.99 | 99.48 | 99.99 | 99.98 | 99.48 | 99.99 |
| SINTAX80 | 99.98 | 99.96 | 99.98 | 99.63 | 99.99 | 99.98 | 99.66 | 100.00 | 99.98 | 99.61 | 99.98 | 99.95 | 99.61 | 99.98 |
| Q1 | 99.83 | 98.12 | 99.83 | 97.75 | 99.82 | 99.83 | 97.73 | 99.81 | 99.83 | 97.79 | 99.83 | 99.66 | 97.58 | 99.68 |
| CT1 | 99.47 | 94.39 | 99.47 | 94.10 | 99.45 | 99.47 | 94.23 | 99.45 | 99.47 | 94.06 | 99.47 | 98.95 | 93.47 | 99.02 |
| Metaxa2 | 99.22 | 92.31 | 99.22 | 92.64 | 99.54 | 99.22 | 93.45 | 99.88 | 99.22 | 91.99 | 99.22 | 98.44 | 91.99 | 99.22 |
| Q2_VS | 96.37 | 86.84 | 96.37 | 87.49 | 96.84 | 96.37 | 89.87 | 97.68 | 96.37 | 86.54 | 96.37 | 92.99 | 84.68 | 94.68 |
| Q2_BLAST | 96.14 | 85.71 | 96.14 | 86.55 | 96.72 | 96.14 | 89.49 | 97.72 | 96.14 | 85.41 | 96.14 | 92.58 | 83.29 | 94.40 |
| CT2 | 95.76 | 81.91 | 95.76 | 84.00 | 96.46 | 95.76 | 88.57 | 97.57 | 95.76 | 81.63 | 95.76 | 91.86 | 79.69 | 93.92 |
| KNN | 84.75 | 45.98 | 84.75 | 49.49 | 89.04 | 84.75 | 56.21 | 95.75 | 84.75 | 45.82 | 84.75 | 73.54 | 45.82 | 84.75 |
| BLCA | 0.00 | 0.00 | 0.00 | 0.00 | 0.00 | 0.00 | 0.00 | 0.00 | 0.00 | 0.00 | 0.00 | 0.00 | 0.00 | 0.00 |

**Table S90. Machine learning metrics computed for the dataset SP RDP ITS 100 at the genus level.**

| Method | Accuracy | Balanced Accuracy | F1-score Micro | F1-score Macro | F1-score Weighted | Precision Micro | Precision Macro | Precision Weighted | Recall Micro | Recall Macro | Recall Weighted | Jaccard Micro | Jaccard Macro | Jaccard Weighted |
|---|---|---|---|---|---|---|---|---|---|---|---|---|---|---|
| BTOP | 100.00 | 100.00 | 100.00 | 100.00 | 100.00 | 100.00 | 100.00 | 100.00 | 100.00 | 100.00 | 100.00 | 100.00 | 100.00 | 100.00 |
| HiTaC | 100.00 | 100.00 | 100.00 | 100.00 | 100.00 | 100.00 | 100.00 | 100.00 | 100.00 | 100.00 | 100.00 | 100.00 | 100.00 | 100.00 |
| KTOP | 100.00 | 100.00 | 100.00 | 100.00 | 100.00 | 100.00 | 100.00 | 100.00 | 100.00 | 100.00 | 100.00 | 100.00 | 100.00 | 100.00 |
| Microclass | 99.99 | 99.99 | 99.99 | 100.00 | 99.99 | 99.99 | 100.00 | 99.99 | 99.99 | 99.99 | 99.99 | 99.99 | 99.99 | 99.99 |
| TOP | 99.99 | 99.99 | 99.99 | 99.99 | 99.99 | 99.99 | 100.00 | 99.99 | 99.99 | 99.99 | 99.99 | 99.99 | 99.99 | 99.99 |
| SINTAX50 | 99.99 | 99.98 | 99.99 | 99.91 | 100.00 | 99.99 | 99.92 | 100.00 | 99.99 | 99.90 | 99.99 | 99.99 | 99.90 | 99.99 |
| SPINGO | 99.98 | 99.97 | 99.98 | 99.90 | 99.99 | 99.98 | 99.92 | 100.00 | 99.98 | 99.89 | 99.98 | 99.96 | 99.89 | 99.98 |
| Q2_SK | 99.97 | 99.87 | 99.97 | 99.80 | 99.97 | 99.97 | 99.81 | 99.98 | 99.97 | 99.79 | 99.97 | 99.94 | 99.77 | 99.95 |
| RDP50 | 99.96 | 99.88 | 99.96 | 99.80 | 99.96 | 99.96 | 99.81 | 99.96 | 99.96 | 99.79 | 99.96 | 99.93 | 99.77 | 99.93 |
| HiTaC_Filter | 99.95 | 99.89 | 99.95 | 99.82 | 99.97 | 99.95 | 99.83 | 99.99 | 99.95 | 99.81 | 99.95 | 99.90 | 99.81 | 99.95 |
| RDP80 | 99.91 | 99.70 | 99.91 | 99.66 | 99.94 | 99.91 | 99.73 | 99.97 | 99.91 | 99.62 | 99.91 | 99.83 | 99.59 | 99.90 |
| SINTAX80 | 99.76 | 99.53 | 99.76 | 99.58 | 99.86 | 99.76 | 99.75 | 99.99 | 99.76 | 99.45 | 99.76 | 99.53 | 99.45 | 99.76 |
| Q1 | 98.97 | 95.51 | 98.97 | 95.49 | 98.92 | 98.97 | 95.83 | 98.96 | 98.97 | 95.43 | 98.97 | 97.97 | 94.69 | 98.21 |
| CT1 | 98.16 | 89.35 | 98.16 | 88.94 | 97.95 | 98.16 | 89.02 | 97.88 | 98.16 | 89.28 | 98.16 | 96.38 | 87.59 | 96.83 |
| Metaxa2 | 97.68 | 87.58 | 97.68 | 88.28 | 98.35 | 97.68 | 89.35 | 99.13 | 97.68 | 87.51 | 97.68 | 95.47 | 87.51 | 97.68 |
| Q2_VS | 84.27 | 57.74 | 84.27 | 57.93 | 84.36 | 84.27 | 60.28 | 86.12 | 84.27 | 57.70 | 84.27 | 72.82 | 54.91 | 79.63 |
| Q2_BLAST | 83.62 | 54.73 | 83.62 | 55.23 | 83.76 | 83.62 | 58.25 | 85.78 | 83.62 | 54.69 | 83.62 | 71.85 | 51.70 | 78.68 |
| CT2 | 82.90 | 51.40 | 82.90 | 53.84 | 83.51 | 82.90 | 60.63 | 86.41 | 82.90 | 51.36 | 82.90 | 70.79 | 48.51 | 77.99 |
| KNN | 57.61 | 17.43 | 57.61 | 19.13 | 62.89 | 57.61 | 23.20 | 73.90 | 57.61 | 17.42 | 57.61 | 40.46 | 17.42 | 57.61 |
| BLCA | 0.00 | 0.00 | 0.00 | 0.00 | 0.00 | 0.00 | 0.00 | 0.00 | 0.00 | 0.00 | 0.00 | 0.00 | 0.00 | 0.00 |

**Table S91. Machine learning metrics computed for the dataset SP RDP ITS 100 at the species level.**

| Method | Accuracy | Balanced Accuracy | F1-score Micro | F1-score Macro | F1-score Weighted | Precision Micro | Precision Macro | Precision Weighted | Recall Micro | Recall Macro | Recall Weighted | Jaccard Micro | Jaccard Macro | Jaccard Weighted |
|---|---|---|---|---|---|---|---|---|---|---|---|---|---|---|
| BTOP | 99.99 | 99.99 | 99.99 | 99.99 | 99.99 | 99.99 | 99.99 | 100.00 | 99.99 | 99.99 | 99.99 | 99.99 | 99.99 | 99.99 |
| HiTaC | 99.97 | 99.99 | 99.97 | 99.98 | 99.97 | 99.97 | 99.98 | 99.98 | 99.97 | 99.99 | 99.97 | 99.94 | 99.97 | 99.95 |
| TOP | 99.56 | 99.50 | 99.56 | 99.43 | 99.51 | 99.56 | 99.50 | 99.60 | 99.56 | 99.50 | 99.56 | 99.12 | 99.24 | 99.28 |
| KTOP | 99.45 | 99.48 | 99.45 | 99.37 | 99.39 | 99.45 | 99.45 | 99.54 | 99.45 | 99.48 | 99.45 | 98.90 | 99.14 | 99.11 |
| Microclass | 99.38 | 99.14 | 99.38 | 99.07 | 99.25 | 99.38 | 99.13 | 99.28 | 99.38 | 99.14 | 99.38 | 98.77 | 98.84 | 98.94 |
| HiTaC_Filter | 97.78 | 97.80 | 97.78 | 98.12 | 98.24 | 97.78 | 98.75 | 99.10 | 97.78 | 97.78 | 97.78 | 95.66 | 97.78 | 97.78 |
| SINTAX50 | 97.50 | 97.08 | 97.50 | 97.32 | 97.83 | 97.50 | 98.00 | 98.70 | 97.50 | 97.07 | 97.50 | 95.13 | 96.86 | 97.22 |
| SPINGO | 96.77 | 96.43 | 96.77 | 97.01 | 97.52 | 96.77 | 98.16 | 98.97 | 96.77 | 96.42 | 96.77 | 93.74 | 96.42 | 96.77 |
| RDP50 | 96.19 | 93.95 | 96.19 | 93.57 | 95.47 | 96.19 | 93.81 | 95.50 | 96.19 | 93.94 | 96.19 | 92.67 | 92.69 | 94.30 |
| Q2_SK | 95.33 | 93.22 | 95.33 | 93.51 | 95.60 | 95.33 | 94.39 | 96.53 | 95.33 | 93.21 | 95.33 | 91.09 | 92.80 | 94.72 |
| RDP80 | 93.02 | 89.86 | 93.02 | 90.09 | 93.13 | 93.02 | 91.08 | 94.13 | 93.02 | 89.85 | 93.02 | 86.96 | 89.20 | 92.00 |
| SINTAX80 | 86.18 | 85.13 | 86.18 | 86.39 | 87.82 | 86.18 | 89.10 | 91.41 | 86.18 | 85.12 | 86.18 | 75.71 | 85.11 | 86.16 |
| Q1 | 83.71 | 73.71 | 83.71 | 72.29 | 81.98 | 83.71 | 72.42 | 82.05 | 83.71 | 73.70 | 83.71 | 71.99 | 69.80 | 78.91 |
| CT1 | 82.65 | 70.33 | 82.65 | 68.46 | 80.39 | 82.65 | 68.33 | 80.17 | 82.65 | 70.32 | 82.65 | 70.44 | 65.78 | 77.14 |
| Metaxa2 | 77.98 | 66.12 | 77.98 | 67.30 | 79.50 | 77.98 | 69.70 | 82.59 | 77.98 | 66.11 | 77.98 | 63.90 | 66.10 | 77.97 |
| Q2_VS | 10.90 | 9.90 | 10.90 | 9.81 | 10.70 | 10.90 | 10.00 | 11.01 | 10.90 | 9.90 | 10.90 | 5.76 | 9.50 | 10.19 |
| Q2_BLAST | 9.27 | 8.10 | 9.27 | 8.08 | 9.17 | 9.27 | 8.39 | 9.60 | 9.27 | 8.10 | 9.27 | 4.86 | 7.76 | 8.65 |
| CT2 | 8.53 | 6.86 | 8.53 | 7.80 | 9.49 | 8.53 | 9.83 | 11.79 | 8.53 | 6.86 | 8.53 | 4.45 | 6.61 | 8.01 |
| KNN | 0.07 | 0.01 | 0.07 | 0.01 | 0.07 | 0.07 | 0.01 | 0.08 | 0.07 | 0.01 | 0.07 | 0.03 | 0.01 | 0.07 |
| BLCA | 0.00 | 0.00 | 0.00 | 0.00 | 0.00 | 0.00 | 0.00 | 0.00 | 0.00 | 0.00 | 0.00 | 0.00 | 0.00 | 0.00 |
